# Deciphering lung adenocarcinoma evolution and the role of LINE-1 retrotransposition

**DOI:** 10.1101/2025.03.14.643063

**Authors:** Tongwu Zhang, Wei Zhao, Christopher Wirth, Marcos Díaz-Gay, Jinhu Yin, Monia Cecati, Francesca Marchegiani, Phuc H. Hoang, Charles Leduc, Marina K. Baine, William D. Travis, Lynette M. Sholl, Philippe Joubert, Jian Sang, John P. McElderry, Alyssa Klein, Azhar Khandekar, Caleb Hartman, Jennifer Rosenbaum, Frank J. Colón-Matos, Mona Miraftab, Monjoy Saha, Olivia W. Lee, Kristine M. Jones, Neil E. Caporaso, Maria Pik Wong, Kin Chung Leung, Chao Agnes Hsiung, Chih-Yi Chen, Eric S. Edell, Jacobo Martínez Santamaría, Matthew B. Schabath, Sai S. Yendamuri, Marta Manczuk, Jolanta Lissowska, Beata Świątkowska, Anush Mukeria, Oxana Shangina, David Zaridze, Ivana Holcatova, Dana Mates, Sasa Milosavljevic, Milan Savic, Yohan Bossé, Bonnie E. Gould Rothberg, David C. Christiani, Valerie Gaborieau, Paul Brennan, Geoffrey Liu, Paul Hofman, Robert Homer, Soo-Ryum Yang, Angela C. Pesatori, Dario Consonni, Lixing Yang, Bin Zhu, Jianxin Shi, Kevin Brown, Nathaniel Rothman, Stephen J. Chanock, Ludmil B. Alexandrov, Jiyeon Choi, Maurizio Cardelli, Qing Lan, Martin A. Nowak, David C. Wedge, Maria Teresa Landi

## Abstract

Understanding lung cancer evolution can identify tools for intercepting its growth. In a landscape analysis of 1024 lung adenocarcinomas (LUAD) with deep whole-genome sequencing integrated with multiomic data, we identified 542 LUAD that displayed diverse clonal architecture. In this group, we observed an interplay between mobile elements, endogenous and exogenous mutational processes, distinct driver genes, and epidemiological features. Our results revealed divergent evolutionary trajectories based on tobacco smoking exposure, ancestry, and sex. LUAD from smokers showed an abundance of tobacco-related C:G>A:T driver mutations in *KRAS* plus short subclonal diversification. LUAD in never smokers showed early occurrence of copy number alterations and *EGFR* mutations associated with SBS5 and SBS40a mutational signatures. Tumors harboring *EGFR* mutations exhibited long latency, particularly in females of European-ancestry (EU_N). In EU_N, *EGFR* mutations preceded the occurrence of other driver genes, including *TP53* and *RBM10.* Tumors from Asian never smokers showed a short clonal evolution and presented with heterogeneous repetitive patterns for the inferred mutational order. Importantly, we found that the mutational signature ID2 is a marker of a previously unrecognized mechanism for LUAD evolution. Tumors with ID2 showed short latency and high L1 retrotransposon activity linked to L1 promoter demethylation. These tumors exhibited an aggressive phenotype, characterized by increased genomic instability, elevated hypoxia scores, low burden of neoantigens, propensity to develop metastasis, and poor overall survival. Re-activated L1 retrotransposition-induced mutagenesis can contribute to the origin of the mutational signature ID2, including through the regulation of the transcriptional factor *ZNF695*, a member of the KZFP family. The complex nature of LUAD evolution creates both challenges and opportunities for screening and treatment plans.

## INTRODUCTION

Lung cancer ranks as the second most prevalent cancer type and stands as the leading cause of cancer-related death worldwide^1^. Among lung cancers, adenocarcinoma (LUAD) is the most prominent histological subtype^2–5^. Understanding tumor evolution could identify tools that could halt its progression or suggest strategies for treatment. Several studies have investigated evolution of different cancer types^6–10^, including LUAD^10–13^. However, these studies have mostly relied on targeted or whole exome sequencing analyses, which limit the investigation of complex genomic alterations such as transposable elements.

Long interspersed nuclear elements-1 (LINE-1 or L1) have emerged as critical players in cancer biology due to their ability to retrotranspose, creating new insertions in the genome^14–18^. While L1 retrotransposons are typically epigenetically silenced in somatic cells, they become aberrantly active in many human malignancies inducing DNA double-strand breaks that can lead to genome instability with local and long-range chromosomal rearrangements, small and large segmental copy-number alterations, and subclonal copy-number heterogeneity^17,19–21^. The activity of L1 retrotransposition has been implicated in promoting heterogeneity in multiple cancer types^22,23^, but its role in shaping LUAD evolution remains underexplored, largely because of the lack of WGS data. Besides our previous work in 232 lung cancer in never-smokers (LCINS)^24^, the only LUAD WGS-based evolution study including smokers and nonsmokers to date included 36 LUAD samples from the Pan-Cancer Analysis of Whole Genomes^9^ (PCAWG). Moreover, previous studies have not investigated LUAD evolutionary trajectories across demographic or exposure scenarios.

We examined 1024 LUAD tumors from the Sherlock*-Lung* study^24,25^, a large integrative study that uses mutational signatures and other genomic features to complement questionnaire, imaging, and exposure assessment approaches to identify potential factors contributing to lung tumorigenesis. Here, we focused on 542 tumors that showed evidence of multiple subclones through deep whole-genome sequencing. Through multiomics analysis, we identified different tumor evolutionary trajectories based on ancestry, tobacco smoking exposure, and sex, and found an aggressive tumor subgroup characterized by specific mutational processes. Furthermore, we characterized the contribution of L1 transposons to mutational processes and tumor evolution. Specifically, we identified *ZNF695*, a member of the KZFP family of transcription factors, as a possible driver of L1 retrotransposition in LUAD. These findings provide valuable insights into how L1 contributes to LUAD progression and heterogeneity.

## RESULTS

### Lung cancer whole-genome sequencing analysis and sample selection

We analyzed deep whole-genome sequencing (mean read depth 81.7x) and other omics data of 1024 LUAD from the Sherlock-*Lung* study^24^; samples were collected from multiple centers across four continents^25^, including 286 smokers, 737 never-smokers, and 1 with unknown smoking status. To ensure adequate statistical power for identifying subclone architectures and constructing lung cancer evolutionary histories, we utilized a metric known as NRPCC (number of reads per tumor chromosomal copy)^26^ to select the appropriate samples for clonal evolution analyses. As expected, our ability to detect subclones was found to depend on the NRPCC, rather than the number of identified mutations (**Supplementary Fig. 1**). By setting a threshold of NRPCC>10, we could reliably identify subclones with a cancer cell fraction (CCF) exceeding 40%. Overall, approximately 53.2% of the LUAD samples exhibited evidence of at least one subclone expansion, which is higher than the LUAD tumors (mostly from smokers) in the PCAWG study (47.4%), possibly due to the higher sequencing depth in Sherlock-*Lung*. We selected 542 LUAD tumor samples with NRPCC>10, purity>0.3 and high-quality copy number profiles since we need clonality data for the downstream tumor evolution analyses (**Supplementary Fig. 2**; **Methods**). We grouped these tumors into four categories based on the patients’ tobacco smoking status and genetic ancestry (**Fig. 1a**; **Supplementary Table 1; Methods)**, including East Asian-ancestry never-smokers, mostly from Taiwan and Hong Kong (AS_N, n=180), European-ancestry never-smokers, mostly from Europe and North America (EU_N, n=184), European-ancestry smokers mostly from Europe and North America (EU_S, n=120), and an indeterminate group (“Others”). The “Others” group comprised 6 African-ancestry individuals, 16 individuals who did not cluster with 1KGP EUR, EAS or AFR reference panels, and 36 from either East Asian or European-ancestry with uncertain tobacco exposure, based on the presence of SBS4 mutational signature and tobacco smoking history.

**Fig. 1:**
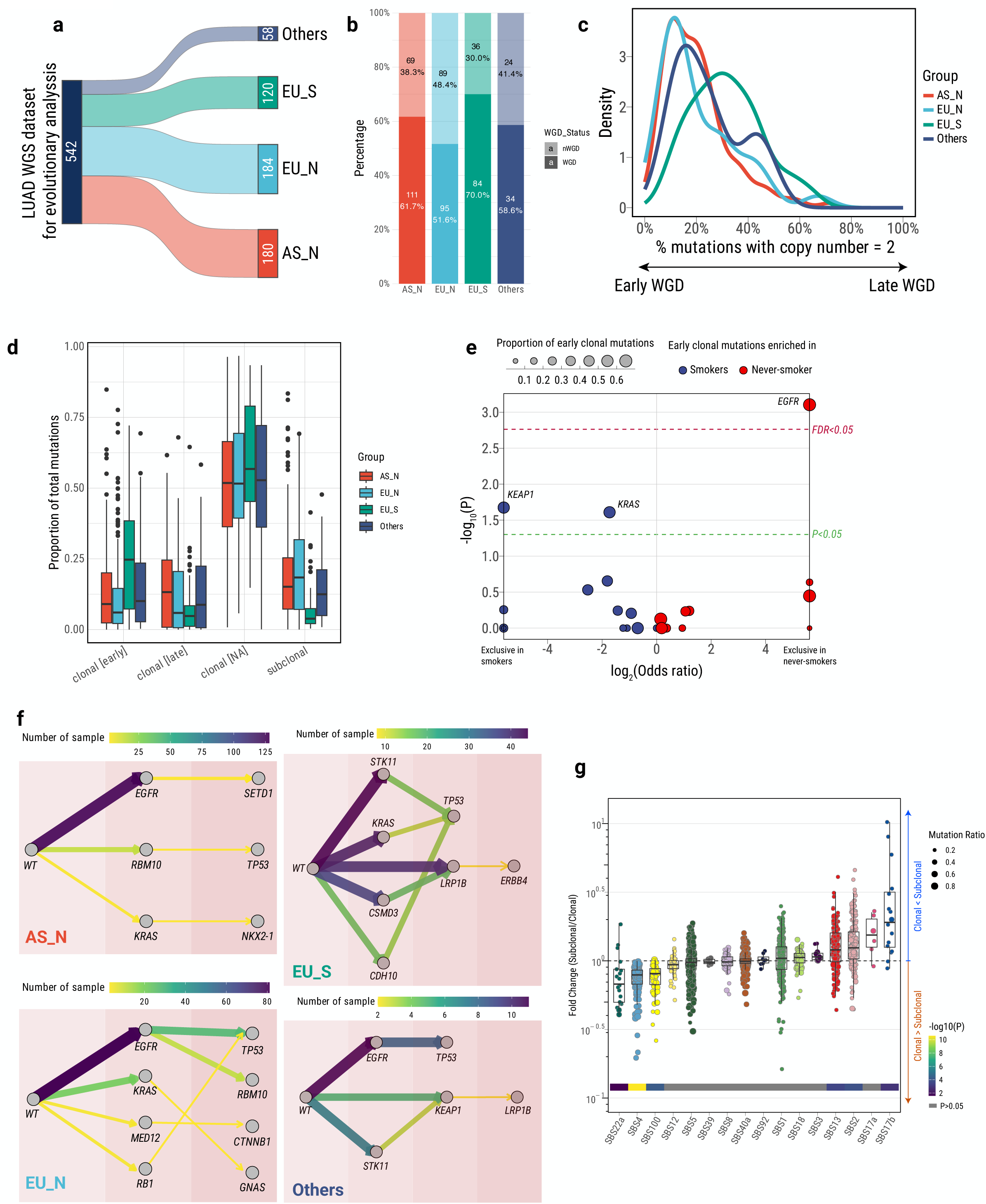
Evolutionary dynamics of lung cancer. **a**) Sankey diagram illustrating high-clonality WGS data summary from Sherlock*-Lung*. **b**) Proportion of tumor samples exhibiting whole genome doubling (WGD) across AS_N, EU_N, EU_S and “Others”. **c**) Distribution of the percentage of mutations with a copy number 2, considering only mutations attributed to clock-like signatures (SBS1 and SBS5). **d**) Box plots depicting the proportion of total mutations attributed to different clonal statuses. **e**) Enrichment of early clonal mutations within driver genes in never-smokers and smokers. For each cancer driver gene, a Fisher’s exact test was performed on a 2×2 contingency table (binary variables: smoking status and early clonal mutation status). The significance thresholds for P < 0.05 (green) and FDR < 0.05 (red), calculated using the Benjamini– Hochberg method, are indicated by dashed lines. **f**) Evolutionary models displaying the recurrent temporal order of driver genes, as inferred by the ASCETIC framework. The color represents the number of samples harboring specific mutation orders. **g**) Dynamic mutational processes during clonal and subclonal tumor evolution. Fold changes between relative proportions of clonal and subclonal mutations attributed to individual mutational signatures. Each point represents a tumor sample, and points are colored by mutational signature. P-values from the Wilcoxon rank-sum test are displayed at the bottom of the boxplots.

### Timing genomic changes in lung cancer

We determined the temporal order of genomic changes in LUAD using whole-genome sequencing data (**Methods**) by assessing the relative timing of somatic mutations and copy number alterations. Whole-genome duplication (WGD)^27^ occurs in 61.7% of AS_N, 51.6% of EU_N, and 70.0% of EU_S tumors (**Fig. 1b; Supplementary Fig. 3)**, with WGD events occurring earlier in never-smokers compared to smokers (two-sided Wilcoxon test, P=5.11e-11; **Fig. 1c**). Chromosomal gains generally occur later in smokers (**Supplementary Fig. 4)**. Notably, early copy number gains on chromosomes 21 and 22 across all groups, and chromosome X in EU_S suggest a role in LUAD tumorigenesis. LUAD from smokers had many early clonal mutations (median 24.7% vs. 9.0% and 6.0% in AS_N and EU_N, respectively), including driver mutations in *KRAS* (**Supplementary Fig. 5**), the dominant oncogene in this group. *KRAS* mutations were enriched with the C:G>A:T type, typically found in the SBS4 signature of tobacco smoking (**Supplementary Fig. 6**). Tumors from never-smokers had a higher proportion of subclonal mutations than those from smokers: 15.1% and 18.4% of somatic mutations in AS_N and EU_N, respectively, compared to 3.9% in EU_S (linear regression, P=2.61e-13 after adjusting for ancestry and tumor purity; **Fig. 1d; Supplementary Table 2**). However, we observed a dominant presence of early clonal *EGFR* driver mutations in never-smokers, accounting for 66.7% in AS_N and 74.7% in EU_N *EGFR*-mutation positive samples, while no *EGFR* driver mutations were observed as early clonal in EU_S (never smokers *vs.* smokers, FDR=0.025; **Fig. 1e** and **Supplementary Fig. 5**). Overall, 81.4% of *EGFR* mutations were attributed to the mutational signatures SBS5 and SBS40a (83.6% in AS_N, 85.8% in EU_N and 35.3% in EU_S; never smokers vs. smokers, p=0.011; **Supplementary Fig. 6**). Both SBS5 and SBS40 are likely the result of endogenous processes^28–32^, predominantly observed in never smokers^33^. This finding highlights the significant role of *EGFR* mutations in the early stages of tumor evolution among never smokers, likely driven by endogenous processes. After multiple testing correction, there was no enrichment of early clonal driver mutations in LCINS compared to EU_S for other cancer driver genes (*e.g.*, *TP53* or *KRAS*).

Driver mutations undergo non-random selection, often leading to similar mutation order across various tumors. We used an agony-based inference framework^34^ to infer these repeated mutational orders. AS_N tumors exhibited large intratumor heterogeneity (ITH), with only a few repeated mutational orders (**Fig. 1f**). In contrast, EU_N showed a clear order *EGFR*-to-*TP53*, which was also observed in the “others” group. Notably, AS_N exhibited higher mutation frequencies for both *EGFR* (72.1% vs. 43.5% in EU_N) and *TP53* (31.1% vs. 25.0% in EU_N) (**Supplementary Fig. 7a**). Additionally, *EGFR* and *TP53* mutations significantly co-occurred in both the AS_N (Fisher’s exact test: P = 0.033, OR = 2.35) and EU_N (P = 0.025, OR = 2.27) groups (**Supplementary Fig. 7b)**. The likely *EGFR* → *TP53* trajectory difference between AS_N and EU_N were confirmed by comparing cancer cell fractions (CCF) in tumors (**Supplementary Fig. 7c)**. *EGFR* appeared to be also earlier than *RBM10* in EU_N, as previously observed^34,35^. Notably, in *TP53* mutation-positive LCINS (N=245), *EGFR* (N=173) and *RBM10* (N=15) mutations were mutually exclusive (P=0.002 in AS_N and P=0.004 in EU_N). In EU_S, *KRAS, STK11*, and *CDH10* mutations preceded *TP53* mutations, while *CSMD*3 mutations preceded *LRP1B* mutations (**Fig. 1f**). These findings emphasize the distinct evolutionary trajectories in LUAD, shaped by factors such as smoking history and ancestry, and the critical role of founder mutations like *EGFR, KRAS*, and *STK11* in the development of LUAD.

### Timing exogenous and endogenous mutational processes

We described the patterns of mutational signatures in all Sherlock-*Lung* tumors in Diaz-Gay et al., MedxRiv 2024. By stratifying the contributing mutations based on their clonal status, we timed mutational signatures, which identified changes in the mutational spectra between clonal and subclonal states and between early and late clonal stages, providing insights into the evolutionary changes driven by dynamic endogenous and exogenous mutational processes. As previously shown for the SBS4 signature^9,36,37^, in our dataset several exogenous mutational processes were found to be predominantly active during the clonal stage of tumorigenesis, as shown by higher proportion of contributing mutations being clonal. These include mutational signatures SBS22a in AS_N (aristolochic acid; median fold change of contributing subclonal/clonal mutations 0.68, IQR 0.49–0.86; two-sided Wilcoxon test with Benjamini–Hochberg correction, FDR=0.026) and SBS4 and SBS100 in EU_S (tobacco smoking associated signatures; SBS4: median fold change 0.79, IQR 0.70–0.88, FDR=6.97e-07; SBS100^38^: median fold change 0.81, IQR 0.70-0.91, FDR=2.21e-03; **Fig. 1g)**. Interestingly, SBS92, which was recently found to be associated with tobacco smoking^39^, showed both clonal and subclonal mutations. Among the signatures with subclonal mutations, both the endogenous mutational processes associated with APOBEC deaminases (SBS2: median fold change 1.24 and IQR 0.95–1.54, FDR=2.33e-03; SBS13: median fold change 1.20 and IQR 0.90–1.51, FDR=2.95e-03) and signatures SBS17a (unknown etiology, median fold change 1.55, IQR 1.16–1.93, FDR=0.65) and SBS17b (unknown etiology, median fold change 1.91, IQR 0.95–2.86, FDR=5.37e-03) had the highest subclonal mutation proportions. However, most APOBEC-related mutations occurred during the clonal stage (**Supplementary Fig. 8**). Intriguingly, in the AS_N tumors, SBS17a was completely lacking and SBS17b was present in only a single tumor, suggesting endogenous or exposure-specific mutational processes in our cohort of European-ancestry (Supplementary Fig. 9), which needs to be further investigated. The comparison between early clonal and late clonal mutations^9^ in mutational signatures (**Supplementary Fig. 10**) identified similar temporal trends to the comparison between clonal and subclonal mutations across the four groups (*e.g.*, SBS4 and SBS22a were enriched with early clonal vs. late clonal mutations in EU_S and AS_N, respectively, **Supplementary Fig. 11**). The analysis of Indel (ID) signatures showed interesting patterns (**Supplementary Fig. 12**), e.g., DNA repair deficiency-associated ID8 and ID6^29^ were enriched in subclonal mutations, although we could not separate the signatures across smoking or ancestry groups, perhaps because of the low numbers of indels.

### Factors influencing lung tumor latency

By focusing on stably accumulating processes such as CpG>TpG mutations, which can act as a molecular clock, we can gain insights into temporal patterns, including tumor latency^9,24^. We define tumor latency as the difference between the age at diagnosis and the estimated age at the appearance of the most recent common ancestor (MRCA) (**Methods; Supplementary Fig. 13**). As expected, the CpG>TpG mutational burden was significantly associated with the age at diagnosis overall (P=2.09e-05; **Supplementary Fig. 14**). Mapping molecular timing estimates to chronological time under different scenarios of increased CpG>TpG mutation rates revealed a similar tumor latency among the three major groups (latency based on constant mutation rate; AS_N=17.0 yrs; EU_N=14.9 yrs; EU_S=14.7 yrs; **Supplementary Fig. 15a-b; Supplementary Table 3**). Of note, tumors in the AS_N cohort had an earlier age at onset (AS_N=63.0 yrs; EU_N=67.9 yrs; EU_S=66.6 yrs; AS_N vs. EU_N P=0.00012; AS_N vs EU_S P=0.003; **Supplementary Fig. 15c**) and an earlier MRCA estimated age when compared to the other groups (**Supplementary Fig. 15d**).

Tumors with shorter latency are more aggressive but clonal, making them easier to target therapeutically, while those with longer latency have greater subclonal diversity, providing a longer detection window but posing therapeutic challenges. Understanding tumor latency is crucial for improving detection and therapy. Thus, we investigated the impact of driver gene mutations on tumor latency. Among the 50 identified driver genes (**Fig. 2a; Supplementary Table 4**), tumors harboring *EGFR* driver mutations exhibited a significantly longer tumor latency than those without *EGFR* driver mutations, with a median of approximately 6.3 more years from the time of subclonal diversification in comparison to tumors with no *EGFR* driver mutations (FDR=8.25e-05). When LCINS were stratified by ancestry, *EGFR*-mutation positive tumors in AS_N and EU_N had average latency increases of approximately 6.2 (P=0.086), and 8.9 years (P=0.00031), respectively, vs. *EGFR* mutation-negative tumors **Supplementary Fig. 16a**). In contrast, tumors with *KRAS* mutations showed a significantly shorter latency in comparison to tumors lacking *KRAS* driver mutations (approximately -5.2 median years; FDR=0.028), particularly in tumors from never smokers (-10 and -9.2 years in EU_N and AS_N, respectively, **Supplementary Fig. 16b**, although based on only 3 AS_N and 21 EU_N). Intriguingly, tumors from female patients had longer latency (median 3.3 more years vs. males) (P=0.0034), mostly in the EU_N subgroup (about 6.4 years) (**Supplementary Fig. 16c**). Multivariable analyses suggest that both *EGFR* mutations and female sex were independently associated with longer latency, but these associations reached statistical significance only in the EU_N subgroup (**Fig. 2b-c**). No difference by sex was observed between tumors with and without *KRAS* mutations (e.g., in EU_S, P=0.83 and 0.62, respectively).

**Fig. 2:**
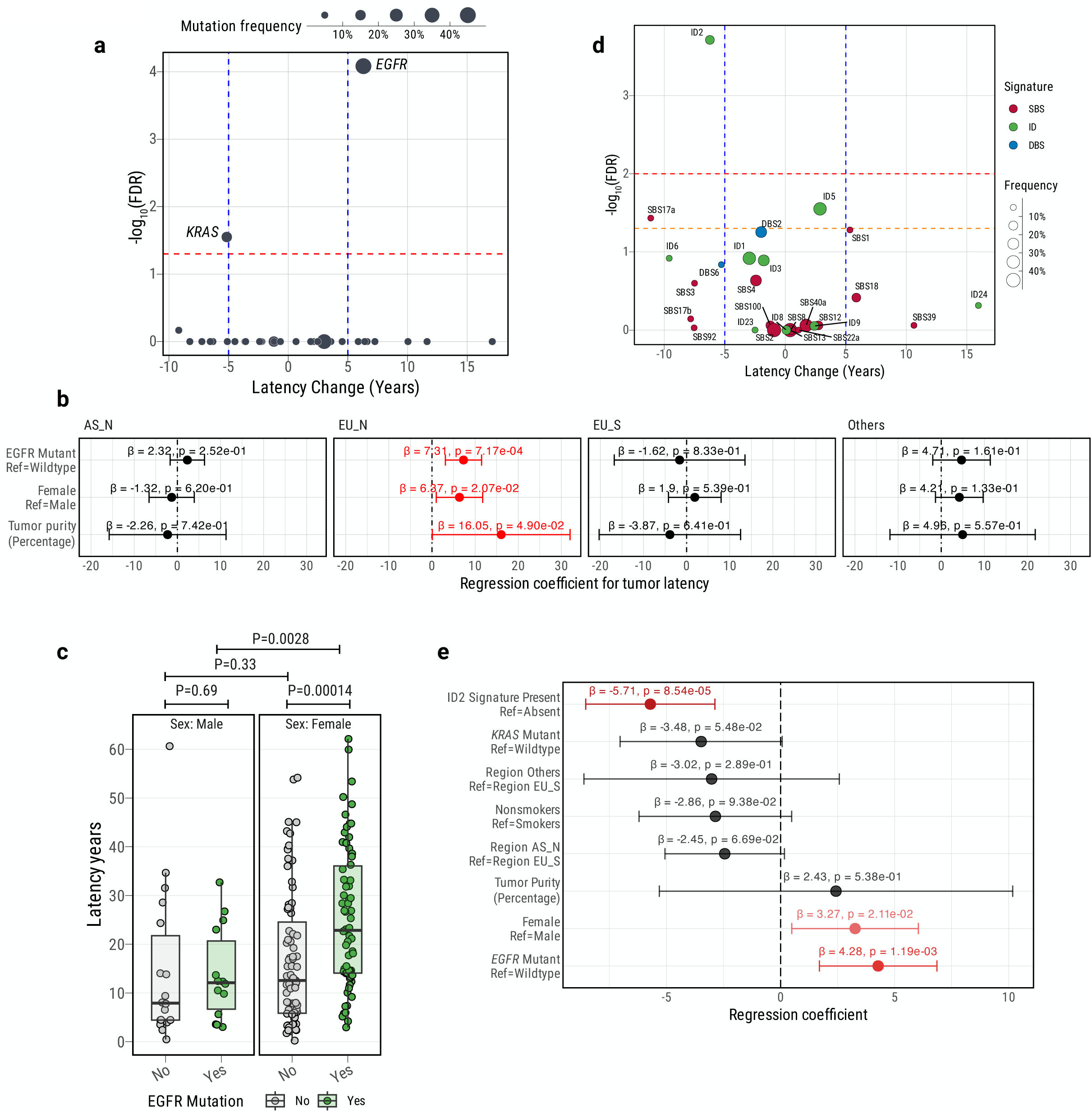
Features associated with lung tumor latency. **a**) Associations between tumor latency and driver gene mutation status. The vertical blue dashed line indicates a 5-year latency difference between driver gene wild type and mutant groups, and the horizontal line represents the significance threshold (FDR<0.05). **b**) Forest plot illustrating associations between tumor latency and *EGFR* mutation status, adjusted for sex and tumor purity (percentage of cancer cells within a tumor sample). P-values and regression coefficients with 95% confidence intervals (CIs) are provided for each variable. Significant associations are in red. **c**) Box plots displaying estimated tumor latency separated by *EGFR* mutation status and sex. **d**) Associations between tumor latency and the presence of specific mutational signatures. The vertical blue dashed line represents a 5-year latency difference between tumors with and without each mutational signature, and the horizontal line indicates the significance threshold (FDR<0.05). **e**) A multivariate regression analysis examining the relationship between tumor latency and various factors, including sex, ancestry, smoking, *EGFR* mutations, *KRAS* mutations, and mutational signature ID2. Statistically significant associations (P<0.05) are denoted in red.

We investigated the association between mutational processes and tumor latency^40^. Among the single base substitutions, the SBS17a signature was nominally associated with shorter latency, regardless of tobacco smoking exposure (**Fig. 2d; Supplementary Fig. 17**). Importantly, the mutational signature ID2 was found to be significantly associated with shorter tumor latency (median 10.5 years, approximately 6.2 median years shorter in comparison with tumors lacking the ID2 signature) after correcting for testing all signature types (FDR = 1.94e-04; **Fig. 2d; Supplementary Table 5**). Similarly, we observed a significant negative correlation between the number of deletions attributed to ID2 and tumor latency (R=-0.15; P=0.0042; **Supplementary Fig. 18**). Previous studies have reported the presence of mutational signature ID2 in LUAD (mostly from smokers) with a frequency of 34%^29,30^. In our study, the frequency of mutational signature ID2 was 11.1%, 16.8%, and 35.8% for AS_N, EU_N, and EU_S, respectively (**Supplementary Fig. 19**). Overall, 83.7% of ID2 deletions were clonal. Consistent with a previous report^41^, we observed that ID2 deletions were associated with late replication timing in lung cancer cell lines (**Methods; Supplementary Fig. 20**). An increased mutation rate related to late replication timing^42^ could contribute to the accumulation of this type of indel.

In a multivariable analysis including sex, ancestry, smoking, *EGFR* mutations, *KRAS* mutations, and mutational signature ID2 in relation to tumor latency (**Fig. 2e**), ID2 had the strongest effect size, followed by *EGFR* mutations and sex, suggesting an independent impact of these factors on LUAD evolution.

### Multi-omic characterization of mutational signature ID2-positive lung tumors

The ID2 signature has an unknown etiology, although it has been found elevated in cancer types with defective DNA mismatch repair and microsatellite instability^29^, which were absent in our samples. To gain insights into the underlying factors contributing to the occurrence of mutational signature ID2, we investigated multiple variables in tumors exhibiting this signature. Short tumor latency suggests aggressive tumor behavior. Thus, we tested the association between tumors with mutational signature ID2 and elevated expression of eight tumor proliferation markers^43^ using RNA-Seq data. As hypothesized, tumors exhibiting ID2 signatures showed significantly higher expression levels of these markers (**Fig. 3a**; e.g., *MKI67*: fold change = 1.2; FDR=9.35e-13; **Supplementary Fig. 21**), and a strong correlation between these markers’ expression levels and the number of ID2-attributed deletions (**Fig. 3b**; *MKI67*: R=0.48; P=8.88e-16). This association was not observed in paired non-tumor lung tissue samples (**Supplementary Fig. 22**). Pathway analyses using QIAGEN Ingenuity Pathway Analysis (IPA) based on RNA-Seq data confirmed the activation of cell proliferation and cell cycle regulation pathways (e.g., Cell Cycle Checkpoints Z=9.43; P=5.23e-31; **Supplementary Fig. 23a**). These findings were further corroborated by Gene Set Enrichment Analyses (**Supplementary Fig. 23b-d**). Furthermore, we found that tumors with the ID2 signature were associated with poor survival (HR=1.8, P=0.0021) and the occurrence of metastases (Fisher’s exact test: OR=2.5, P=5.58e-03; Logistic regression: P=0.039 after adjust tumor purity, age, sex and *TP53* mutation status) (**Fig. 3c-d; Supplementary Fig. 23e-h**).

**Fig. 3:**
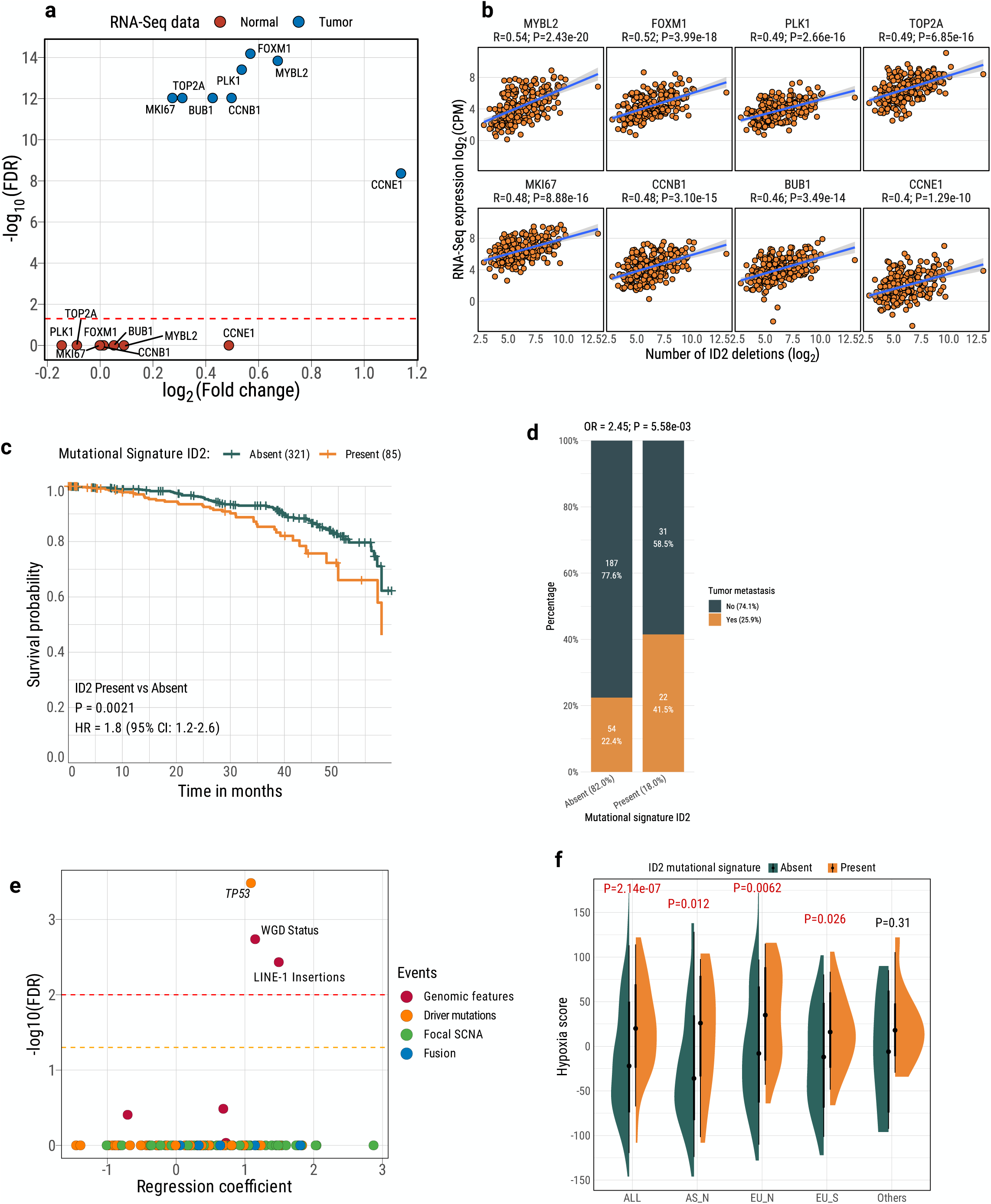
Characterization of tumors with mutational signature ID2. **a**) Relationships between mutational signature ID2 presence and gene expression of tumor proliferation markers, analyzed from tumor and normal tissue RNA-Seq data. The horizontal line represents the significance threshold (FDR<0.05). **b**) Pearson correlation between the number of deletions attributed to mutational signature ID2 and the gene expression of tumor proliferation markers. Pearson correlation coefficients and corresponding p-values are displayed above the plots. **c**) Kaplan-Meier survival curves for overall survival, stratified by the presence of mutational signature ID2. Significance p-values and Hazard Ratios (HRs) were calculated using two-sided Cox proportional-hazards regression, adjusting for age, sex, smoking, and tumor stage. Numbers in brackets indicate the number of patients. **d**) Enrichment of tumor metastasis in tumors with mutational signature ID2. Odds ratios and p-values from the Fisher exact test are shown above the plot. **e**) Enrichment of genomic alterations (WGD=whole genome doubling; SCNA=somatic copy number alterations) in tumors with mutational signature ID2, determined through logistic regression and adjusted for ancestry, sex, smoking status, age, and tumor purity. The horizontal lines represent significance thresholds (FDR<0.05 in orange and FDR<0.01 in red). **f**) Distribution of estimated hypoxia scores between tumors with ID2 signatures and those without. P-values from the Wilcoxon rank-sum test are displayed above the plot.

A multivariable logistic regression model including age, sex, ancestry, smoking, and tumor purity as covariates showed that *TP53* driver mutations, WGD status and LINE-1 retrotransposon insertions were the most significant genomic events enriched in ID2 tumors after multiple testing correction (**Fig. 3e)**. In fact, *TP53* mutations were associated with the number of ID2 deletions (**Supplementary Fig. 24**). *TP53* was downregulated in ID2-positive tumors in comparison with ID2-negative tumors (IPA upstream transcriptional regulator analysis; Z=-6.81; P=1.81e-31), indicating a high level of genomic instability^44^ (**Supplementary Fig. 25**). Hypoxia has been suggested to play a crucial role in influencing the genomic, mutational, and evolutionary characteristics of cancers^45^ and could develop because of the rapid tumor growth that outstrips the oxygen supply^46,47^. Consistently, we found a considerably elevated hypoxia score in tumors exhibiting the ID2 signature in comparison to ID2-negative tumors across all three major groups (**Methods; Fig. 3f**).

A shorter tumor latency may be partly due to poor immune editing. To explore this hypothesis, we estimated the neoantigens predicted from mutation data and calculated the propensity of each SBS and ID mutational signature to generate neoantigens. Intriguingly, ID2 showed the least propensity among all signatures (**Supplementary Fig. 26**), which indicates a diminished predisposition for immunoediting, likely contributing to the observed shorter latency and accelerated tumor proliferation. Furthermore, immune cell decomposition analysis using various RNA-Seq-based methods^48^ consistently revealed that tumors with ID2 signatures exhibited a marked depletion of T cells and myeloid dendritic cells in their microenvironment compared to other tumors (**Supplementary Fig. 27**). These findings suggest that impaired immune surveillance may play a critical role in the aggressive progression of ID2-positive tumors.

### Frequent somatic L1 retrotransposition in LUAD and its contribution to mutational signature ID2

LINE-1 insertions were strongly associated with ID2-positive tumors (**Fig. 3e**), independent of *TP53* mutations (association in *TP53* mutant tumors: P=0.000129; OR = 5.57; in *TP53* wild-type tumors: P=0.00211; OR=3.73). When considering the ID2 mutational signature status, we found that among 50 curated hallmark pathways from the mSigDB gene sets^49^ and 658 pathways from the KEGG MEDICUS database as well as a genome-wide survey of genes involved in L1 retrotransposition control from CRISPR/Cas9 screening in two distinct human cell lines^50,51^, the L1-regulated gene sets exhibited significant enrichment in the ID2-positive tumors (**Supplementary Fig. 28**), ranking immediately after the top cell proliferation pathways (**Supplementary Fig. 23b-d**).Previous studies have highlighted the role of L1 retrotransposition in initiating and driving tumorigenesis in humans^52^, predominantly in tumors originating from epithelial tissues, including lung cancer^14^. In light of this, we conducted an investigation into L1 activities from germline and somatic sources in lung cancer and their impact on tumor evolution and ID2 signatures, leveraging our deep whole genome sequencing data (examples of germline and somatic sources of L1 retrotransposition in **Fig. 4a-b**; **Methods**). Our analysis revealed a significantly higher number of somatic L1 insertions in tumors from smokers compared to those from never-smokers (P=4.87e-07), as previously shown^53^. Overall, among the somatic L1 insertions with a known source, the majority were attributable to a preexisting germline L1 element (70.4%). A germline source suggests a role of this retrotransposition in the initiation (first clonal expansion) of the evolutionary trajectory. Interestingly, AS_N tumors exhibited a higher proportion of L1 retrotransposition originating from a somatic source of L1 elements compared with the other two groups (AS_N: 21.0%, EU_N: 8.3%, and EU_S: 4.3%; **Fig. 4c**; **Supplementary Table 6**). Consistent with previous reports^15,17^, we identified several master germline sources of L1 elements across all groups, including chr22q12.1 (28.9%), chrXp22.2 (14.2%), chr2q24.1 (4.6%), chr14q23.1 (4.4%), and others (**Fig. 4c**).

**Fig. 4:**
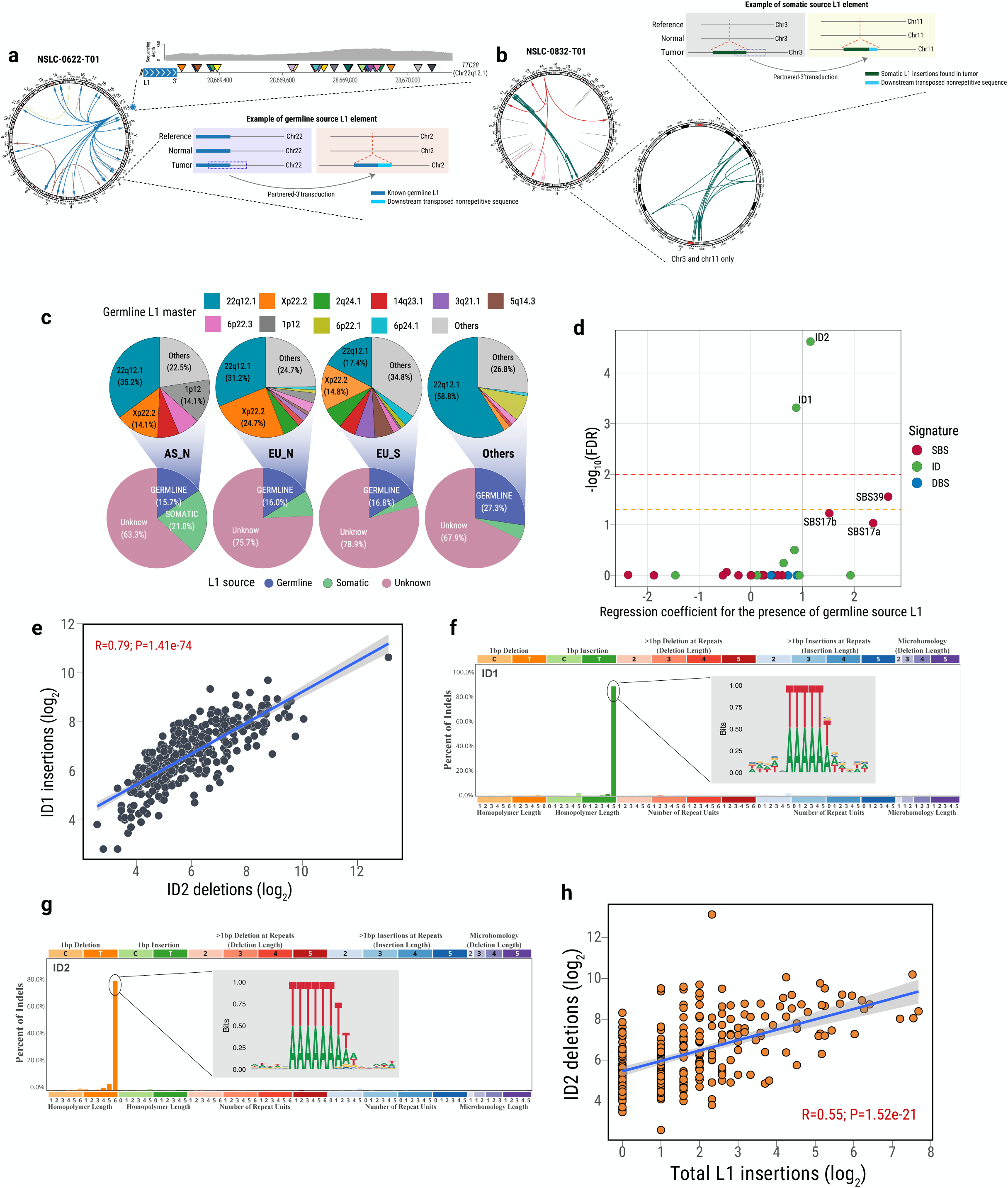
Association between L1 retrotransposition and mutational signatures ID2 and ID1. **a**) This panel illustrates a sample (NSLC-0622-T01) as an example of a tumor harboring L1 insertions from a germline source. In the Circos plot, an arrow indicates the direction from the location of L1 elements in the human germline genome to the position of L1 somatic insertions in the tumor genome. The gray line without an arrow in the Circos plot indicates L1 insertions with an unknown source. L1 retrotranspositions originating from the master L1 on chromosome 22q12.1 (highlighted in blue in the Circos plot) are zoomed in. Different partnered 3’ transductions are highlighted by triangles with various colors on chromosome 22, accompanied by a sequence depth plot (gray bars). A specific partnered 3’ transduction (including the L1 repetitive element and the adjacent unique sequence) between chromosome 22 and chromosome 2 serves as an example of one germline-source L1 retrotransposition. **b**) This section presents another example (tumor sample NSLC-0832-T01) predominantly characterized by somatic-source L1 insertions. In the Circos plot, a green arrow highlights multiple L1 retrotranspositions detected solely in the tumor genome. For improved visualization of these somatic-source L1 insertions, a zoomed-in view specifically focusing on the somatic retrotransposition between chromosome 3 and chromosome 11 is provided in the second Circos plot. Additionally, a specific partnered 3’ transduction serves to elucidate somatic-source L1 retrotransposition. **c**) Distribution of retrotransposable sources of L1 insertions. The bottom pie chart displays the percentage of L1 insertions retrotransposed from germline, somatic, and unknown L1 elements. The top pie chart show the proportion of L1 insertions originating from specific germline L1 masters. **d**) Enrichment of mutational signatures in tumors with germline source L1 insertions. The horizontal lines indicate the significance threshold FDR < 0.05 in orange and FDR < 0.01 in red. **e**) Pearson correlation between deletions attributed to signature ID2 and insertions attributed to signature ID1. Pearson correlation coefficients and corresponding p-values are displayed in the plot. **f-g**) Mutational signature profiles and motifs for mutational signature ID1 and ID2, respectively. **h**) Pearson correlation between deletions attributed to ID2 and total somatic L1 insertions. Pearson correlation coefficients and corresponding p-values are shown in the plot.

L1 retrotransposon activity in tumors is associated with genomic instability and can contribute to tumorigenesis by inducing DNA damage^54–56^ and/or disrupting the coding sequences of driver genes^57^. Thus, we explored the association between the number of L1 retrotransposon insertions and mutational processes using logistic regression (**Methods**). As expected, we observed a significant association between L1 insertions from a germline source and mutational signature ID2 (FDR=2.8e-05), and also with ID1 (FDR=5.71e-04; **Fig. 4d**). Intriguingly, there were strong correlations among the numbers of indels assigned to these two signatures (**Fig. 4e**), suggesting a shared underlying etiology. Mutational signatures ID1 and ID2 are characterized by 1 base pair insertions and deletions, respectively. They occur in similar mutation contexts within polymers of at least 5 As or Ts. Motif analysis of all the indels assigned to mutational signature ID1 and ID2, confirmed the minimum homopolymer lengths of 5 and 6 Ts, respectively (**Fig. 4f-g**). Even with a lower number of L1 insertions from a somatic source, we observed a significant association of L1 insertions with the ID2 signature (**Supplementary Fig. 29**). Additionally, we observed a strong positive correlation between the number of L1 insertions and the number of deletions attributed to mutational signature ID2 across all three major groups (Overall: R=0.55; P=1.52e-21; AS_N: R=0.37; P=3.62e-04; EU_N: R=0.46; P=1.58e-06; EU_S: R=0.44; P=3.06e-03; **Fig. 4h**). We validated this association in an independent TCGA LUAD dataset (R=0.55; P=4.1e-08; **Methods; Supplementary Fig. 30**). Furthermore, in the tumors exhibiting the mutational signature ID2 (vs. tumors without ID2), we observed a stronger enrichment of L1 insertions from a germline source (OR=3.57 P=1.23e-08) than somatic transposed L1 insertions (OR=2.33, P=1.94e-04; **Supplementary Fig. 31**). These findings suggest a likely contribution of L1 retrotransposition to the development of the mutational signature ID2 and its associated early clonal indels and short tumor latency. Consistently, tumor samples harboring L1 insertions, particularly from a germline source, showed a shorter tumor latency (**Supplementary Fig. 32**).

### Reactivation of L1 elements through DNA hypomethylation in ID2-positive tumors

The intricate process of retrotransposition involves the synthesis and insertion of new L1 copies into the genome through targeted site-primed reverse transcription (TPRT)^58^. During TPRT, the endonuclease activity of ORF2p, encoded by the L1 ORF2 gene, cleaves target DNA or other TPRT intermediates, resulting in DNA double-strand breaks (DSB)^55^ or DNA lesions that subsequently activate DNA repair pathways (e.g., non-homologous end-joining)^59–65^. Remarkably, ORF2p specifically identifies and cleaves a consensus sequence of 5’-TTTTT-3’ (canonical endonuclease motifs) within the genome^16,66,67^, which bears intriguing similarities to the mutational signature ID1 and ID2 motifs. To prevent activation of L1, epigenetic suppression, including methylation of L1 promoters, is known to play an important role^68–70^. We hypothesized that L1 is reactivated through promoter demethylation. Reactivated L1 ORF2-encoded endonuclease, coupled with DNA repair mechanisms, could contribute to the emergence of single base pair indels^65^ like ID2 across the tumor genome (**Fig. 5a**).

**Fig. 5:**
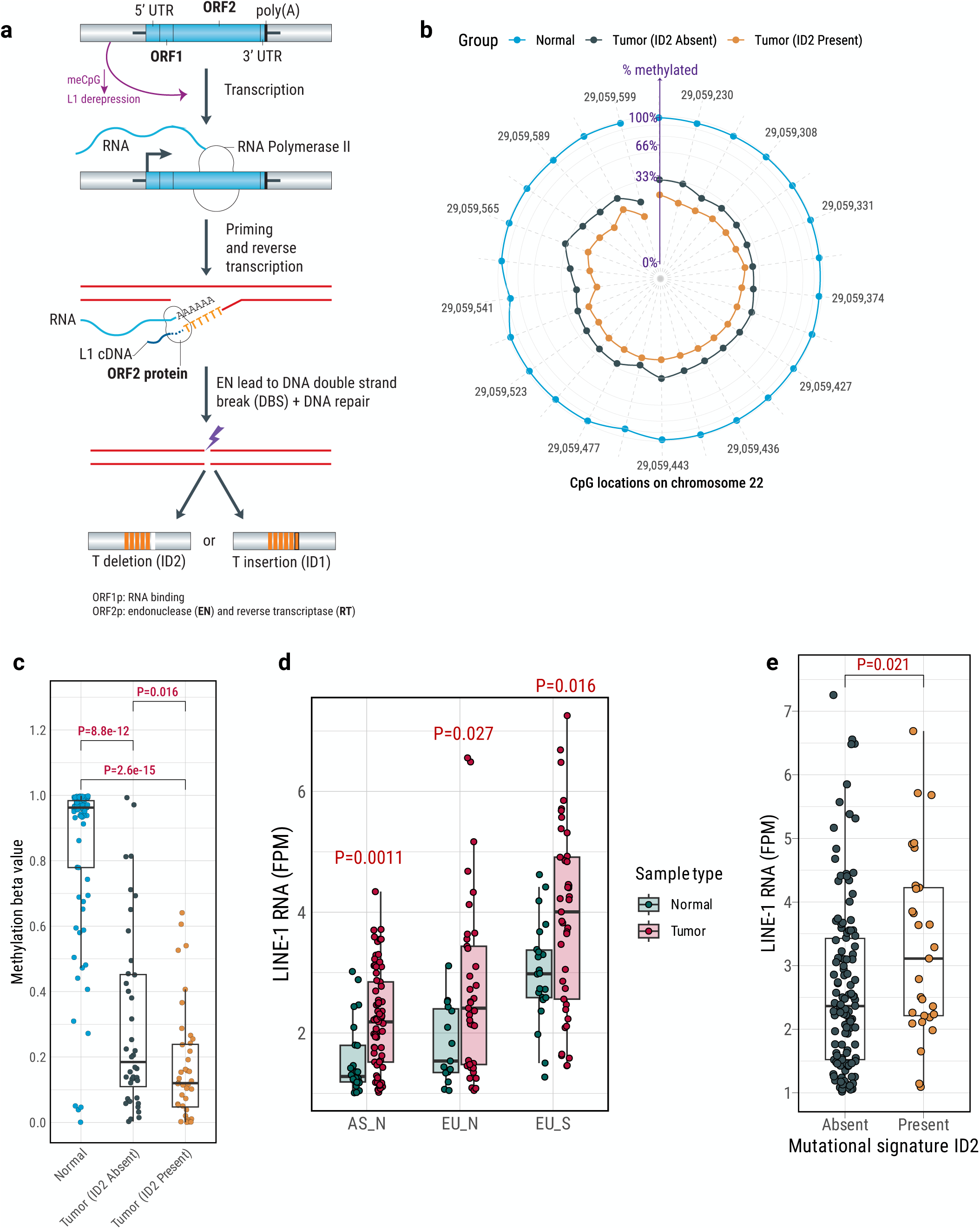
Activation of germline L1 retrotransposition due to DNA demethylation of L1 promoter. **a**) Diagram illustrating the transposon mobilization mechanism for long interspersed element 1 (L1). Adapted from a publication by Levin and Moran^58^, this mechanism depicts non-LTR retrotransposons mobilizing through target-site-primed reverse transcription (TPRT). ORF2-encoded endonuclease generates a single-strand ‘nick’ in genomic DNA, freeing a 3′-OH used to prime RNA reverse transcription. Demethylated CpG in the L1 promoter region (top purple arrow) is hypothesized to activate L1 retrotransposition. Endonuclease activity, coupled with DNA repair mechanisms, might lead to one-base pair deletions or insertions at polymer A/T regions. **b**) Validation of DNA methylation levels in the promoter region of germline L1 insertions from germline source in chr22q12.1, conducted via targeted bisulfite sequencing. The polar plot represents the median methylation level across the genome locations of the CpG island on chr22q12.1, stratified by normal lung samples (N=80), tumor samples without ID2 signature (N=40), and tumors samples with ID2 signature (N=40). Control samples were designed to represent 0%, 33.3%, 66.6%, and 100% methylation levels at each CpG site as shown on the y-axis. **c**) Box plot shows DNA median methylation levels across the genome locations of the CpG island on chr22q12.1, stratified by the sample type and ID2 status. **d**) Total L1 RNA expression estimated from RNA-Seq data, stratified by sample type and group. **e**) Total L1 RNA expression differs between tumors with and without the ID2 signature.

In support of the hypothesis of DNA hypomethylation contributing to L1 retrotransposition, DNA methylation array data revealed that the large majority (95.4%) of differentially methylated CpG probes displayed a global lower methylation level in tumor samples with ID2 signatures compared to those without ID2 (FDR<0.01, **Supplementary Fig. 33**). This observation is also consistent with previously found tobacco smoking association with global CpG hypomethylation in lung and other tissues^71–73^ and hypomethylation of L1 elements in the blood of smokers of cigarettes ^74^ or e-cigarettes^75^. In fact, we found the ID2 signature more frequently in tumors from smokers. Furthermore, to estimate the methylation status of the L1 promoter, we performed targeted bisulfite sequencing of the most active germline L1 element on chr22q12.1 in 80 tumors (40 ID2 positive and 40 ID2 negative; **Methods**). Consistent with our hypothesis, the L1 promoter region displayed notable demethylation in tumor samples characterized by high frequencies of L1 insertions and ID2 deletions. In contrast, L1 promoter regions remained hypermethylated in matched normal samples. Tumor samples lacking ID2 deletions had intermediate levels of methylation (**Fig. 5b-c**). A negative correlation was observed between CpG methylation level and mutations attributed to mutational signature ID2 (**Supplementary Fig. 34**). Further analyses revealed higher levels of both global and locus-specific L1 RNA expression in tumor tissues compared to normal tissues across all three major groups (EU_N, AS_N, and EU_S; **Fig. 5d; Supplementary Fig. 35**), as well as significantly higher expression in tumors with ID2 compared to those without ID2 (**P = 0.021**; **Fig. 5e**). As previously suggested^74^, individuals with a history of smoking in our study exhibited enhanced L1 RNA expression in both tumor and normal tissues in comparison with never smokers. Importantly, the L1 RNA expression was higher even in individuals who had quit tobacco smoking, suggesting a long term effect of tobacco smoking-related L1 retrotransposition (**Supplementary Fig. 36**), possibly through long-lasting epigenomic or transcriptional changes that continue to influence L1 retrotransposition activity even after smoking cessation^76^. Collectively, these findings illustrate a role for reactivated L1 elements in inducing the mutational signature ID2 in LUAD.

### *ZNF695* expression is linked to L1 retrotransposition and ID2 signatures in LUAD

To further investigate factors that may affect the L1 promoter methylation in tumors, we analyzed the Kruppel-associated box zinc finger proteins (KRAB-ZFPs/KZFPs), the largest family of transcription factors in humans, known to play an essential role in the recognition and transcriptional silencing of transposable elements, including L1^77^. KZFPs can induce cell type specific L1 repression^78^. To identify which KZFPs could be involved in germline L1 retrotransposition activation in LUAD, and in particular in LUAD enriched with the ID2 signature, we conducted analyses of RNA-Seq data for 471 previously identified KZFP genes^78^. The majority of expressed KZFP genes (390/471) displayed a positive correlation with each other (**Supplementary Fig. 37**). We observed that 77.4% of KZFP genes (n=302) exhibited significantly differential expression in tumors compared to the normal lung tissue (62.9% up-regulated and 37.1% down-regulated; **Supplementary Fig. 38a**), with the largest fold increase observed for *ZNF695* (Fold change=4.8; FDR=1.73e-45). Notably, 43 of these 302 genes also exhibited differential expression in tumors from smokers compared to never smokers, including *ZNF695* (Fold Change=2.7; FDR=0.01; **Supplementary Fig. 38b-d**), suggesting that tobacco smoking could have a role in modifying the levels of these genes. Among all KZFP genes, *ZNF695* was the most significant gene up-regulated in tumors with ID2 signature compared to those without ID2 (Fold Change=3.5, FDR=1.14e-08; **Fig. 6a-b**), even adjusting for other covariables (**Supplementary Fig. 39**). Furthermore, expression of *ZNF695* in tumors showed the most significant positive correlation with the deletions attributed to ID2 signatures (R=0.47; P=1.11e-14; **Fig. 6c-d**). Similar patterns were found between the *ZNF695* expression and tumor L1 insertions (**Supplementary Fig. 40**). Moreover, the *ZNF695* binding motif was enriched in the sequences (1kb) surrounding the demethylated CpG probes in the promoters of germline L1 compared to random sequences (P=0.0028; FDR=0.016). We also observed a negative correlation between the expression of *ZNF695* and methylation level of L1 promoter in chr22q12.1 in tumor tissues but not in normal (**Fig. 6e**). As expected, ZNF695 expression was strongly correlated with the expression of tumor proliferating markers (**Supplementary Fig. 41**). Furthermore, *ZNF695* expression was elevated in lung epithelial cells but was low in immune cell types, suggesting a potential inverse relationship with immunogenicity (**Supplementary Fig. 42a**). Intriguingly, type I pneumocytes, the well differentiated alveolar epithelial cells involved in oxygen exchange, showed much lower *ZNF695* expression than the stem-like type II pneumocytes, from which lung adenocarcinoma may originate^79^, supporting the hypothesis of a *ZNF695* role in adenocarcinoma genesis. This hypothesis was further supported by our lung single-nucleus multiomics study profiling gene expression and chromatin accessibility^80^. In this study, *ZNF695* expression was highest in a proliferating sub-population of alveolar type 2 (AT2pro) cells among other normal lung cell types identified (**Supplementary Fig. 42b**). Cells with detectable *ZNF695* expression also exhibited significantly higher L1 expression (P=0.04; **Supplementary Fig. 42c**), further supporting an association between *ZNF695* and LINE-1 activity.

**Fig. 6:**
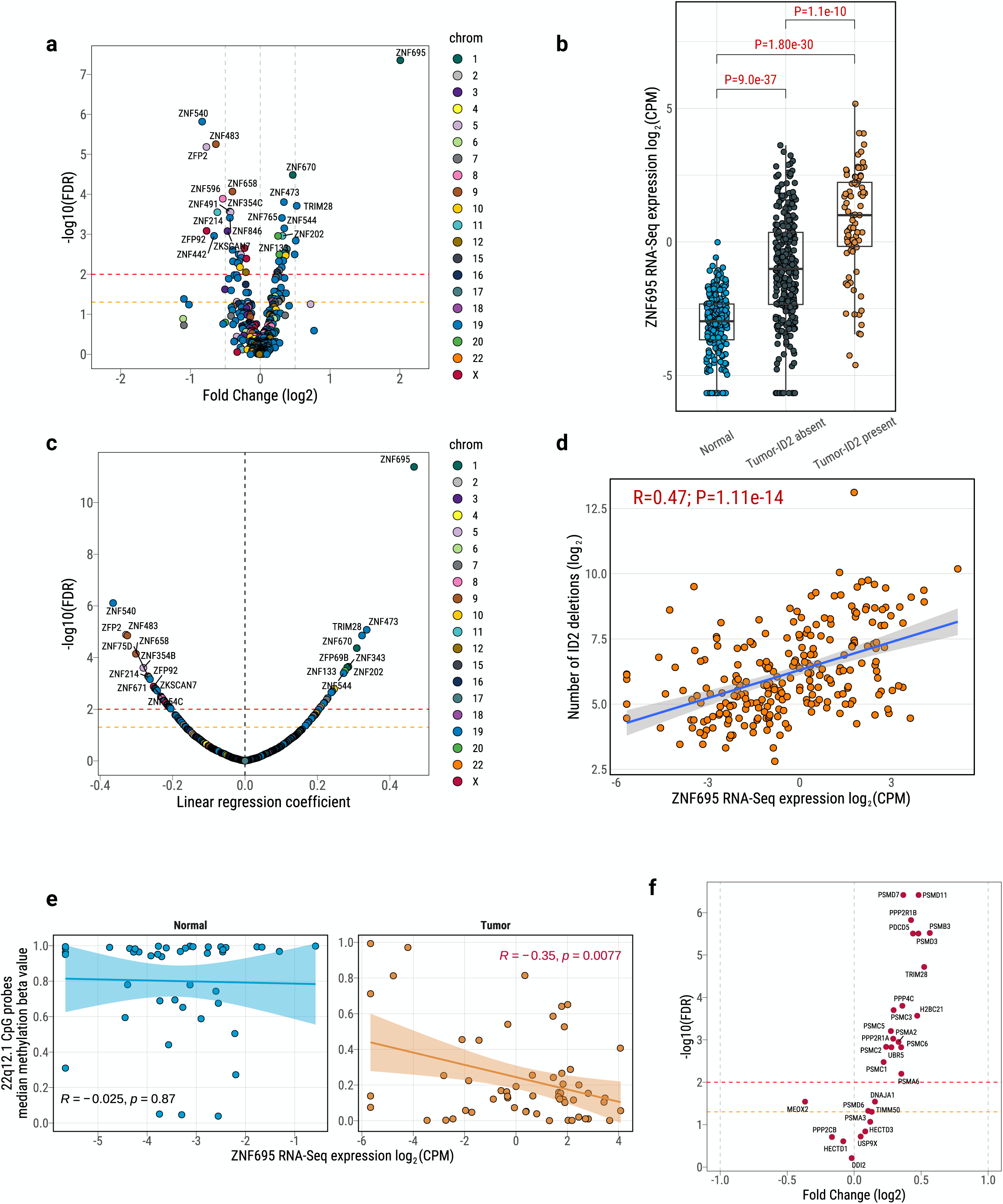
*ZNF695* upregulation in tumors and its association with mutational signature ID2. **a)** Analysis of differentially expressed KZFP protein coding genes between tumors with ID2 signatures and those without. Horizontal dashed lines represent significance thresholds (FDR < 0.05 in orange and FDR < 0.01 in red). The top 20 significant genes are annotated with gene names. **b**) Box plots illustrate the differential expression of *ZNF695* among normal tissue or blood, tumors without ID2 signatures, and tumors with ID2 signatures. **c**) Pearson correlations between KZFP protein-coding gene expression and indels attributed to mutational signature ID2. Horizontal dashed lines indicate significance thresholds (FDR < 0.05 in orange and FDR < 0.01 in red). The top 20 significant genes are annotated with gene names. **d**) Correlation between *ZNF695* expression and deletions attributed to mutational signature ID2. Pearson correlation coefficients and corresponding p-values are displayed above the plot. **e**) Correlation between ZNF695 RNA-Seq expression and the median of DNA methylation levels across the genome locations of the CpG island on chr22q12.1. Pearson correlation coefficients and corresponding p-values are displayed. **f**). Differentially expressed ZNF695 target genes identified between tumors with and without mutational signature ID2.

To further confirm the link between *ZNF695* expression and L1 retrotransposition/ID2 signatures, we examined the association between ID2 signatures and the transcriptional activity of *ZNF695*, as indicated by the expression of its target genes. Specifically, we identified 28 potential *ZNF695* target genes from curated databases^81,82^ and found that 79% of these were differentially expressed between tumors with and without the ID2 signature (**Fig. 6f)**. Additionally, analysis of 70 downstream target genes regulated by KRAB-ZFPs^83^ revealed that 67% were differentially expressed with respect to ID2 status (**Supplementary Fig. 43)**. These findings strongly support a role for KRAB-ZFPs, including ZNF695, in the transcriptional regulation associated with ID2/LINE1 activity.

We extracted all RNA sequencing reads corresponding to *ZNF695* and examined junction reads along with sequencing coverage. Intriguingly, we observed elevated levels of non-canonical transcripts in LUAD tumors compared to their paired normal samples (**Supplementary Fig. 44**). Furthermore, we noted increased expression of non-canonical transcripts in tumors exhibiting mutational signature ID2 compared to those without ID2. Canonical transcripts are translated into zinc-finger proteins that can recruit DNA methyltransferases, which maintain the DNA methylation that is essential for repressing germline L1 retrotransposition activity^84^. In contrast, non-canonical transcripts, lacking the DNA-binding zinc-finger domain, may result in decreased methylation levels at the L1 locations due to the dominant negative effect (**Supplementary Fig. 45)**. This observation suggests an interesting hypothesis of a dosage competition effect for non-canonical transcripts in ID2-positive tumors.

## DISCUSSION

In this comprehensive study of lung adenocarcinoma evolution, we utilized deep whole-genome sequencing and multi-omics analyses on high-clonality samples to determine the relative timing of various genomic changes and examine their associations with tobacco exposure, demographic factors, and clinical outcomes.

Major cancer driver genes, such as *EGFR* (dominant in never smokers) and *KRAS* (in smokers) have been found to be mutated in histologically normal lung tissues^85^ and in adenocarcinoma in situ (AIS)^35^. We report here that they are mutated very early in LUAD evolution. The timing of the mutational signatures show that the signatures associated with exogenous exposures, such as SBS4 and SBS100 (tobacco smoking), mostly include clonal mutations, reflecting an early impact on tumor initiation, while endogenous-related signatures (APOBEC, SBS17a/b) appear later in the evolution. Taken together, one could expect to see early clonal driver gene mutations enriched with early occurring mutational patterns. Indeed, in smokers’ tumors we found *KRAS* oncogene hotspots enriched with C:G>A:T transversions, the most frequent mutational type associated with tobacco smoking exposure. However, *EGFR* mutations in never smokers were mostly attributed to age or unknown endogenous processes (SBS5/SBS40a signatures). Moreover, we observed that tumors harboring these driver genes have profoundly different latency, with slow and fast subclonal diversification in *EGFR*-mutant and *KRAS*-mutant tumors, respectively. These results suggest that there could be two divergent lung adenocarcinoma evolutionary trajectories between smokers and never smokers.

Specifically, in never smokers, the epithelial tissue and related cancer driver genes accumulate mutations with increasing age^86^. Mutant clones compete with one another^87^ until one or a few clones (for example carrying *EGFR* mutations) acquires a fitness advantage and expands. The selection and expansion do not appear to be due to exogenous mutational processes, but they may be affected by a still unknown, likely endogenous, mutagenic process, although generating a low number of mutations (e.g., SBS40a) or a promotion effect, e.g., by stimulating inflammation^24,85^. The competition among subclones^87^ continues during tumor progression, maintained by the lack of strong mutagenic effects facilitating specific subclones’ positive selection and by poor immunoediting effects due to the low mutational burden^88^. Together, these factors contribute to the long latency, which was particularly evident in tumors from female patients, possibly reflecting sex-differences in immune responses^89^. This suggests that future or ongoing (e.g., NCT05164757) lung cancer screening programs for female nonsmokers could consider long intervals between scans.

On the contrary, in smokers, the age-related accumulation of mutations is increased by the strong mutagenic effect of tobacco smoking over time. Once *KRAS*-associated clones are positively selected (*KRAS* has enrichment of C:G>A:T mutations in the hotspot), or sensitized through smoke-induced epigenomic changes^90^, they rapidly expand, facilitated by the genomic instability, chromosomal rearrangements, frequent double strand breaks associated with LINE 1 retrotransposition and other tobacco smoking-related factors. In fact, tumors harboring *KRAS* mutations have much shorter subclonal diversification than *KRAS*-negative tumors. In both never smokers and smokers of European descent, *EGFR* and *KRAS* appear to precede the acquisition of *TP53* alterations, which in turn increase genomic instability and tumor growth.

Intriguingly, tumors in the Asian-ancestry LCINS samples appeared to follow yet another evolutionary pattern: AS_N tumors had much higher *EGFR* mutation frequency than EU_N tumors, often co-occurring with *TP53* alterations. AS_N *EGFR*-mutation positive tumors had a short clonal evolution (early age at MRCA occurrence), and presented with heterogeneous repetitive patterns of mutational order. These findings suggest that multiple competing forces are acting in these tumors. Of note, although SBS22a appears to be mostly clonal, EGFR and other major cancer driver genes in the Taiwanese samples are not enriched with the typical T:A>A:T mutation patterns of SBS22a. Thus, it is unclear whether this signature reflects a causative link between aristolochic acid exposure and LCINS carcinogenesis as has been observed in mice models of lung cancer^91^ and other tumor types^92^, or just the exposure to aristolochic acid in this population.

Importantly, we found that the mutational signature ID2 is a marker of a novel mechanism for LUAD tumor evolution, which, if validated, could be added to the arsenal of precision medicine tools for prognosis and treatment. The indels contributing to the mutational signature ID2 were mostly clonal, thus present at tumor initiation. Similarly, ID2 was strongly enriched in tumors with short tumor latency. Both the associations of ID2 with tumor initiation and short latency were independent from major driver genes or demographic features affecting tumor evolution. This signature, which was two-to-three times more frequent in smokers than nonsmokers, was enriched in fast growing tumors with poor prognosis. Crucially, we showed that this signature, previously of unknown etiology, could be a by-product of the activation of L1 retrotransposition, often from germline sources. The aggressive phenotype of ID2-positive tumors could be due to L1-related genomic instability^21,93–95^, oncogene activation^96^, tumor suppressor inactivation^19^, metabolic reprogramming^97^, and immune modulation^98^. L1 is the most abundant transposon element family in humans^99^, occupying ∼17% of genomic DNA. L1 could be reactivated in the genome through demethylation consequent to tobacco smoking exposure^76,100–102^ and reduced epigenetic regulation. We found that *ZNF695*, a transcription factor upregulated in ID2-positive tumors, and located on chr.1q, a typically amplified region in lung cancers, was enriched with non-canonical transcripts lacking the DNA-binding zinc-finger domain and unable to recruit DNA methyltransferases, likely resulting in decreased methylation levels at the L1 locations due to a dominant negative effect. While these findings provide strong correlative evidence linking *ZNF695* to L1 regulation, further experiments, such as CRISPR-based functional studies and single-cell approaches, will be required to validate this relationship and elucidate the mechanistic role of ZNF695 in regulating L1 retrotransposition as well as its potential as a therapeutic target. Of note, previous studies may have failed to recognize the important role of L1 retrotransposition on LUAD evolution because they were largely based on whole exome sequencing data (*e.g.*, TRACERx), which cannot capture non-coding genomic regions, including transposable elements.

Different evolutionary trajectories may impact clinical course and treatment responses. The longer latency of EGFR-mutant tumors may allow the accumulation of mutations increasing resistance to tyrosine kinase inhibitors (e.g., the EGFR T790M mutation)^103,104^, highlighting the potential value of upfront combination therapies to suppress treatment resistance mediated by the outgrowth of competing clones under selective therapeutic pressure. At the same time, the long latency of EGFR-mutant cancers provides a wide time window for tumor early detection. In contrast, the short subclonal diversification seen in *KRAS*-mutation positive tumors suggests a more focused and potent KRAS-directed therapeutic strategy. Given the distinctly aggressive phenotype of ID2-positive LUADs, the presence of ID2 signature could be explored as a potential biomarker to escalate therapies. While the low mutation-related neoantigen in ID2-positive LUADs could lead to poor immune recognition and may limit the clinical efficacy of currently available immune checkpoint inhibitors, the enrichment of Helper T Cells in the tumor microenvironment suggests that novel immunotherapies harnessing these existing T cell subsets could represent a more effective strategy. L1, the ORF2 protein, and transposable elements are the object of increasing research into their role in stabilizing the genome, their dynamic relationship with DNA methylation^105^ and their potential to be therapeutically targeted^67,106^, possibly exploiting tumor-specific antigens derived from transposable elements^107^. These findings highlight the need for future research to explore the relationship between tumor latency, genomic changes, transposable elements, and clinical outcomes in LUAD.

In summary, our findings provide new insights into LUAD tumorigenesis, underscoring the role of demographic features, exposures, and repetitive elements in shaping distinct evolutionary trajectories.

## METHODS

### Collection of Lung Cancer Samples

In the Sherlock-*Lung* study, we identified 1024 patients with histologically confirmed lung adenocarcinoma from multiple institutions (**Supplementary Table 1**). The fresh frozen tumor tissue with matched whole blood samples or fresh frozen normal lung tissue sampled, when possible, at least 3 cm from the tumor were collected from these treatment-naïve patients. We followed rigorous criteria for sample inclusion, similar to previous publications^24^: 1) Sequencing coverage. We maintained a minimum average sequencing coverage of >40x for tumor samples and >25x for normal samples. 2) Contamination and Relatedness. Cross-sample contamination was limited to <1% by Conpair^108^, and detected relatedness was maintained <0.2 by Somailer^109^; 3) Copy number analysis. Subjects with abnormal copy number profiles in normal samples were excluded, as determined by Battenberg^110^. 4) Mutational signatures. Tumor samples exhibiting mutational signatures SBS7 (two samples, associated with ultraviolet light exposure) and SBS31 (one sample, associated with platinum chemotherapy) were removed from the analysis. 5) WGS quality control. Tumor samples with a total genomic alteration count of <100 or <1000 along with NRPCC (the number of reads per clonal copy)^26^ <10 were excluded. 6) Multiple-region sequencing. In cases where multiple regions of a tumor were sequenced, only one high-purity tumor sample was included to avoid redundancy. These stringent criteria were applied consistently to ensure the robustness and reliability of the data collected for the Sherlock-*Lung* study.

Since NCI exclusively received de-identified samples and data from collaborating centers, had no direct contact or interaction with the study subjects, and did not utilize or generate any identifiable private information, Sherlock-*Lung* has been classified as “Not Human Subject Research (NHSR)” in accordance with the Federal Common Rule (45 CFR 46; eCFR.gov).

### Specimen Processing

The methods for specimen collection and processing have been detailed in our prior publication^24^. Genomic DNA was extracted from fresh frozen tissue following the protocols outlined in the QIAmp DNA Mini Kit (Qiagen) user manual. The concentration of DNA was determined using a Nanodrop spectrophotometer. Subsequently, DNA quantification, normalization, and fragmentation were performed using the same procedure as our standard WGS process^24^. Each DNA sample was required to meet minimum mass and concentration thresholds and had to exhibit no signs of contamination or profile discordance as indicated by the Identifiler assay. Samples meeting these requirements were then subdivided into aliquots, with mass adjusted to meet the specific requirements for downstream assay processing.

### Whole genome sequencing processing

Whole genome sequencing library construction followed our previously reported protocol^24^. The Broad Institute (https://www.broadinstitute.org) conducted WGS sequencing on the Illumina HiSeq X and NovaSeq6000 platforms following Illumina protocols for 2×150bp paired-end sequencing. FASTQ files were generated after Illumina base-calling. Paired FASTQ were converted into unmapped BAM using the GATK pipeline, available at https://github.com/gatk-workflows/seq-format-conversion. Subsequently, the unmapped BAM files were further processed using GATK on the cloud-based platform TERRA workspaces (https://app.terra.bio). The sequence data were aligned to the human reference genome GRCh38, and the resulting aligned BAM files were then transferred to the NIH HPC system (https://hpc.nih.gov) for subsequent downstream analyses.

For the public WGS data, the preprocessed aligned BAM/CRAM files (including unmapped reads) were first converted back to FASTQ files using Bazam (v.1.0.1)^111^ to retain the sequencing lane and read group information and then processed using the same pipeline as for the Sherlock-*Lung* WGS dataset. As a validation dataset to examine the relationship between ZNF695, mutational signature ID2, and L1 insertions, we downloaded newly released raw BAM files of TCGA LUAD samples from the NCI GDC data portal (https://portal.gdc.cancer.gov/). We excluded tumor samples previously analyzed as part of the Sherlock-*Lung* dataset, resulting in a total of 258 TCGA LUAD samples for validation. We followed the same analytical pipelines as in the Sherlock-*Lung* dataset.

### Define high-quality WGS dataset for tumor evolution analysis

As indicated in the PCAWG study^26^, our ability to detect subclones was determined by the NRPCC value, rather than the quantity of identified mutations^26^. Therefore, for tumor evolution analysis, we selected a set of 542 LUAD tumor samples with high-quality WGS data as our discovery set. The selection was based on the following criteria. Firstly, we required a tumor purity greater than 0.3, indicating a substantial proportion of tumor cells within the samples. Secondly, we set a minimum NRPCC value of 10 to ensure the robustness of subclone detection throughout tumor evolution analysis. Lastly, we only included samples with high-quality copy number profiles generated by Battenberg^110^, without any involvement of refitting steps, thereby eliminating the potential somatic copy number artifacts. We used VerifyBamID2 to project germline genetic principal components for these samples into the principal component space defined by the 1000 Genome Project (1KGR) reference panel (**Supplementary Fig. 2**). We labeled samples that clustered with the 1KGP EUR sample as “European-ancestry” (n=335); those that clustered with the 1KGP EAS sample as “East Asian-ancestry” (n=185); and those that clustered with the 1KGP AFR sample as “African-ancestry” (n=6). These analyzes included 400 non-smokers and 141 smokers, and 362 females and 180 males.

### Identification of Genome-Wide Somatic Alterations, including SNVs, SVs, and transposable elements

The somatic mutation calling was performed using our established bioinformatics pipeline as previously described^24^. Our approach involved applying four distinct mutation calling algorithms for tumor-normal paired analysis, including Strelka (v.2.9.10), MuTect, MuTect2, and TNscope implemented in the Sentieon’s genomics software (v.202010.01). We employed an ensemble method to merge the results from these different callers followed by additional filtering to reduce false positive calling. The final mutation calls for both SNVs and indels were required to meet the following criteria: 1) read depth >12 in tumor samples and >6 in normal samples; 2) variant allele frequency <0.02 in normal samples, and 3) overall allele frequency (AF) <0.001 in multiple genetics databases including 1000 Genomes (phase 3 v.5), gnomAD exomes (v.2.1.1) and gnomAD genomes (v.3.0). For indel calling, we retained only variants identified by at least three algorithms. The IntOGen pipeline (v.2020.02.0123)^112^, which combines seven state-of-the-art computational methods, was employed with default parameters to detect signals of positive selection in the mutational patterns of driver genes across the cohort.

We employed the Battenberg algorithm (v.2.2.9)^110^ to conduct analyses of somatic copy number alterations (SCNA). Initial SCNA profiles were generated, followed by an assessment of the clonality of each segment, purity, and ploidy. Any SCNA profile determined to have low-quality after manual inspection underwent a refitting process. This process required new tumor purity and ploidy inputs, either estimated by ccube (v.1.0)^113^ or recalculated from local copy number status. The Battenberg refitting procedures were iteratively executed until the final SCNA profile was established and met the criteria of manual validation check. GISTIC (v.2.0) was used to identify the recurrent copy number alterations at the gene level based on the major clonal copy number for each segmentation. Meerkat (v.0.189)^114^ and Manta (v.1.6.0)^115^ were applied with recommended filtering for identifying structural variants (SVs), and the union set of these two callers was merged as the final SV dataset.

To identify putative transposable elements (TE), we utilized a pipeline called TraFiC-mem (v.1.1.0) (Transposome Finder in Cancer)^15^ retaining only TE insertions that passed the default filtering criteria. The source element of L1 retrotransposition can be classified into two categories^15,17^: germline and somatic source L1 elements. Germline source L1 elements are identified by unique DNA regions retrotransposed somatically elsewhere in the cancer genome from a single locus of a reference full-length L1 element or a putative non-reference polymorphic L1 element. Somatic source L1 elements are those derived from the downstream region of a putative L1 event present in the tumor genome but absent in the matched normal genome.

Collectively, we define the overall somatic alterations as the summary of SNVs, indels, SVs, and TE insertions.

### Defining driver mutations

To identify driver mutations within the set of identified driver genes, we implemented a rigorous and multifaceted strategy, considering multiple criteria: (a) the presence of truncating mutations specifically in genes annotated as tumor suppressors, (b) the recurrence of missense mutations in a minimum of 3 samples, (c) mutations designated as “Likely drivers” with a boostDM score^116^ exceeding 0.5, (d) mutations categorized as either “Oncogenic” or “Likely Oncogenic” based on the criteria established by OncoKB^117^, an expert-guided precision oncology knowledge base, (e) mutations previously recognized as drivers in the TCGA MC3 drivers paper^118^, and (f) missense mutations characterized as “likely pathogenic” in genes that are annotated as tumor suppressors, as described in Cheng et al^119^. Any mutation meeting one or more of these criteria will be recognized as a potential driver mutation. This comprehensive approach ensures a thorough and exhaustive identification of driver mutations within the context of the identified driver genes.

### Timing mutations and copy number gains using MutationTimeR

We employed the R package MutationTimeR (v1.00.2) to estimate the timing of somatic mutations relative to clonal and subclonal copy number states and to calculate the relative timing of copy number gains^9^. This analysis utilized all somatic mutations, including single-nucleotide variants (SNVs) and indels, alongside the final copy number profiles as input data for the MutationTimeR algorithm.

MutationTimeR integrates mutation allele frequencies, local copy number states, and tumor purity to determine whether mutations occurred before or after copy number gains. The algorithm identifies whether mutations are present on alleles in chromosome regions with gains or losses. Using allele-specific copy number profiles, it assigns each mutation to a specific copy number state.

For diploid regions without copy number gains, MutationTimeR classifies mutations as clonal if they are present on one copy per cell or as subclonal if present on fewer than one copy per cell. In regions with copy number gains (e.g., bi-allelic gains), clonal mutations are further classified into two temporal categories based on their relationship to the copy number event:

1. **Early Clonal Mutations**: Mutations occurring before the copy number gain and present on all tumor cell copies (≥2 copies per cell). These mutations are clonal and often represent early driver events.
2. **Late Clonal Mutations**: Mutations occurring after the copy number gain, present on only one copy per cell.

Using Bayesian inference, MutationTimeR estimates the likelihood of mutations occurring in the context of specific copy number changes, enabling precise temporal classification.

For cancer driver genes, we calculated the proportion of each mutational clonality category across all samples. Additionally, we aggregated clonality information for each mutational signature by considering the probability of each mutation type being assigned to its respective signature.

### Infer the temporal patterns for the driver genes

To reconstruct robust tumor evolution models, we utilized the ASCETIC framework (Agony-baSed Cancer EvoluTion InferenCe) developed by Fontana et al.^34^. ASCETIC’s methodology involves the reconstruction of robust tumor evolution models for individual patients, which are subsequently integrated into a comprehensive cancer-specific evolution model. Specifically, for each group (AS_N, EU_N, EU_S and others), we generated a binary matrix, where a value of 1 indicated the presence of driver mutations in the sample and 0 indicated their absence. Furthermore, mean Cancer Cell Fraction (CCF) and Variant Allele Frequency (VAF) data were leveraged to estimate the temporal progression of each group, serving as input for ASCETIC. The ASCETIC algorithm was then executed with a total of 100 resampling iterations to reliably determine the timing of the driver genes. The outcome of this analysis yielded an evolutionary model illustrating the chronological acquisition of driver genes during cancer evolution.

Additionally, we manually examine the evolutionary trajectory by conducting a mutually exclusive analysis and comparing the order of cancer cell fractions between two mutations. For instance, to validate the *EGFR* → *TP53* trajectory difference between AS_N and EU_N, we directly compared cancer cell fractions (CCF) in tumors with detectable subclones (CCF < 0.8), which approximate mutation ordering. We defined a potential *EGFR* → *TP53* trajectory when: CCF*_EGFR_* - CCF*_TP53_* > 0.2, or *EGFR* is in the right side of clonal peak (higher CCF) and *TP53* is in the left side of clonal peak or subclonal peak (lower CCF). A Fisher’s exact test was performed to determine whether the EGFR → TP53 trajectory was enriched in EU_N.

### Chronological time analysis to estimate tumor latency

To determine the temporal expansion of tumors, we utilized a methodology of chronological time analysis derived from previous studies^9,24^, including the estimation of the emergence of the most recent common ancestor (MRCA) and the tumor latency between the MRCA and the age at diagnosis (**Supplementary Fig. 13**). The approach assumes a constant mutation rate per year, particularly for CpG>TpG mutations caused by the spontaneous deamination of 5-methylcytosine to thymine at CpG dinucleotides, which exhibit clock-like behavior. In summary, a hierarchical Bayesian linear regression model was utilized to establish the relationship between the age at diagnosis and the scaled number of clock-like CpG>TpG mutations occurring in an NpCpG context (SBS1). This model accounted for key variables, including tumor ploidy, tumor purity, and subclonal architecture, as described in the PCAWG evolutionary study^9^ (https://github.com/gerstung-lab/PCAWG-11/).

For each tumor, the age at the occurrence of the MRCA was estimated. Given the low mutational burden observed in lung cancer in never smokers (LCINS), the most frequent group in our cohort, an estimated tumor acceleration rate of 1x was applied. Tumor latency, representing the time from subclonal diversification to diagnosis, was calculated as the difference between the estimated age at the MRCA and the age at tumor diagnosis. An implicit assumption is that the latency between the emergence of the last detectable subclone and the age at diagnosis is relatively short compared to the overall timeline of tumor development. This approach provides critical insights into the evolutionary dynamics of tumors, highlighting differences in clonal expansion and subclonal diversification across tumor subtypes.

### Mutational signature analysis

The detailed methods for the mutational signature analysis have been included in a separate mutational signature manuscript (Diaz-Gay et al., MedxRiv 2024). Briefly, SigProfilerMatrixGenerator^120^ was used to create the mutational matrices for all types of somatic mutations, including single base substitutions (SBS), doublet base substitutions (DBS), and indels (ID). *De novo* mutational signatures for SBS, DBS, and ID were extracted using SigProfilerExtractor^39^ (v1.1.21) with default parameters, normalization set to 10,000 mutations, and splitting the cohort between smokers, and non-smokers and unknown smoking status, in order to limit the effect of hypermutators in the signature extraction process. *De novo* signatures were extracted using the SBS-288, DBS-78, and ID-83 mutational contexts, respectively, and later decomposed into COSMICv3.4 reference signatures^121^, based on the GRCh38 reference genome, and assigned to individual samples using SigProfilerAssignment^122^ (v0.1.0). Two novel signatures were identified, one smoking-associated SBS signature, termed SBS100, extracted *de novo* in the smoker cohort and presenting similarities with prior signatures reported in Signal ^38^; and one ID signature, termed ID24, extracted *de novo* in the non-smoker cohort, and subsequently used to decompose other *de novo* extracted signatures in the smoker cohort.

### RNA-Seq data analysis

The RNA-Seq data from the Sherlock-*Lung* study were generated at the Cancer Genomics Research Laboratory (CGRL), Division of Cancer Epidemiology and Genetics, National Cancer Institute, using the Illumina HiSeq platform based on the Illumina TruSeq Stranded Total RNA-Seq protocol, which resulted in the production of 2×100bp paired-end reads. The RNA-seq data from the TCGA-LUAD study^123^ were downloaded from the GDC Data Portal^124^. The FASTQ files were aligned to the human reference genome GRCh38/hg38 using STAR (v.2.7.3a)^125^, annotated using the GENCODE v35, and processed by HTSeq (v.0.11.4)^126^ for gene quantification. The resulting expression data were corrected for batch effects using the R package ComBat-seq (v.3.48.0)^127^ and normalized to Counts Per Million (CPM), followed by log2 transformation using the R package edgeR (v.3.42.4)^128^. Only ‘expressed’ genes defined by CPM >0.1 in at least 10% QC-passed samples were included in the final quantification data. Pathway analyses were performed using Ingenuity Pathway Analysis (IPA) software and also the traditional Gene Set Enrichment Analysis (GSEA) using the hallmark gene sets from mSigDB database (https://www.gsea-msigdb.org/). The TIMER2.0 web tool^48^ was used for immune cell decomposition based on the normalized gene expression matrix.

### Differential Expression Analysis of KRAB-ZFP Target Genes

For the transcription factor ZNF695, we curated target genes from two major databases: the transposable element enrichment KRAB-ZFP database (KRABopedia: https://tronoapps.epfl.ch/web/krabopedia/) and the IntAct molecular interaction database (https://www.ebi.ac.uk/intact/home). Notably, the KRABopedia database indicated that LINE-1 elements are among the most significantly enriched peaks in ZNF695 ChIP-Seq experiments. Integration of these two sources resulted in a collection of 28 potential ZNF695 target genes. Additionally, we obtained 70 downstream target genes regulated by general KRAB-ZFPs from a previous study^83^. To confirm the link between ZNF695 expression and ID2 signatures, we examined the association between ID2 signatures and the transcriptional activity of KRAB-ZFPs, as indicated by the expression levels of these target genes. Differential expression analysis was performed using the Wilcoxon rank-sum test on the RNA-Seq data.

### Quantification of L1 RNA in bulk RNA sequencing data

L1 locus-specific RNA expression was quantified using the L1EM software^129^ based on RNA-Seq data. L1EM employs the expectation-maximization algorithm to estimate gene expression specific to L1 loci and to differentiate proper L1 expression from passive co-transcription, which includes L1 RNA but does not support retrotransposition. To quantify intact L1 RNA, we considered full-length L1 loci with no stop codon present in either ORF1 or ORF2 as expressed if they meet the following criteria: at least two read pairs per million (FPM) were assigned to that locus, and less than 10% of the RNA assigned to that locus was estimated to be result of passive co-transcription. The total expression of intact LINE-1 RNA was estimated by aggregating the FPM values for each locus.

### Analysis of *ZNF695* and L1 expression in single-cell RNA sequencing data

To investigate ZNF695 expression at the single-cell level, we queried the cellxgene database (https://cellxgene.cziscience.com/gene-expression), a comprehensive repository of curated single-cell RNA sequencing (scRNA-seq) datasets, to evaluate ZNF695 expression across diverse cell types from lung tissue studies. Expression data, including raw and scaled values, were extracted from lung-specific datasets, with cell types annotated based on the original study metadata.

Additionally, we analyzed ZNF695 and L1 expression using our previously published lung single-cell multiomics dataset^130^. Raw sequencing data in FASTQ format were downloaded from this study to quantify L1 expression. To improve the accuracy of transposable element (TE) expression profiling, particularly for L1 elements, we applied the Stellarscope pipeline^131^, a bioinformatics tool that reassigns multi-mapped reads to specific TE loci using an expectation-maximization algorithm. Stellarscope was run with default parameters to generate a per-cell L1 count matrix. This matrix, along with cell annotations, was processed using standard scRNA-seq workflows in Seurat (v5.0). Normalized L1 expression values from specific loci were aggregated to calculate overall L1 expression per cell. Cells were then classified based on detectable ZNF695 expression, determined from gene expression profiles generated in Seurat. Differential L1 expression between ZNF695-positive and ZNF695-negative cells was evaluated using a Wilcoxon rank-sum test, with statistical significance defined as P < 0.05.

### Genome-wide DNA methylation data analysis

Genome-wide DNA methylation was profiled on the Illumina HumanMethylationEPIC BeadChip (Illumina, San Diego, USA). Genomic DNA was extracted, and DNA methylation was measured according to Illumina’s standard procedure at the Cancer Genomics Research Laboratory (CGRL), Division of Cancer Epidemiology and Genetics, National Cancer Institute. Raw DNA methylation data (“.idat” files) were generated for our study, combined with TCGA LUAD^123^ methylation data collected from the GDC Data Portal^124^ (Illumina Human Methylation 450k BeadChip), and processed using RnBeads (v.2.0)^132^. Background correction (“enmix.oob”) and beta-mixture quantile normalization (“BMIQ”) were applied. Unreliable probes (Greedycut algorithm with detection P<0.05), cross-reactive probes, and probes mapping to sex chromosomes were removed. Samples with outlier intensities in 450k/EPIC array control probes were removed from the dataset described in the RnBeads vignette. Principal component analysis (PCA) of these samples was performed based on the genotyping probes and we removed subjects with Euclidean distance between matched tumor/normal pair > upper quartile + 3*inter-quartile range (IQR). In addition, we performed Combat batch correction separately for tumors and normal samples using the “sva” R package^133^ on M-values for known technical factors, including studies, collection sites, sample plates, sample wells, array types (450K and EPIC), chip ID, and positions. We used MBatch (v2.0, https://github.com/MD-Anderson-Bioinformatics/BatchEffectsPackage) to assess the presence of batch effects. The Dispersion Separability Criteria (DSC) values were <0.5 for all technical factors, suggesting that batch effects were negligible after batch correction. The corrected methylation-level beta values were used for downstream association analyses.

### Neoantigen analysis

To predict neoantigens, patient-specific HLA haplotypes were identified using HLA-HD (v.1.2.1)^134^. The software NetMCHpan4.1^135^ was then run on 9-11 neo peptides derived from all nonsynonymous mutations, taking into account the patient’s specific HLA genotypes. NetMCHpan4.1 was included in the NeoPredPipe pipeline (v.1.1)^136^. The decomposed mutation probabilities generated by the SigProfilerExtractor were used to predict the neoantigens derived from specific mutational signatures. For TCGA LUAD data, the predicted SNV neoantigen counts per sample were collected from a previous publication^137^. We used the number of peptides predicted to bind to MHC proteins as the neoantigen load for our analysis.

### Enrichment of *ZNF695* motif in the germline L1 retrotransposon insertions

The differentially methylated probe analysis between tumors exhibiting ID2 signatures and those without, was performed by the Wilcoxon rank sum testing. We identified significantly demethylated probes in ID2 positive tumors with a false discovery rate <0.05 after multiple testing corrections. Subsequently, the sequences surrounding the demethylated CpG probes of germline L1 insertions (+/-1kb) were extracted and used as input for the transcription factor enrichment analysis. This analysis was performed using the Simple Enrichment Analysis (SEA) algorithm from the MEME suite (v.5.5.4). The input motifs were selected from the Homo Sapiens motif database (CIS-BP 2.00 Single Species DNA), which includes the transcription factor *ZNF695* motif.

### Hypoxia score calculation

Hypoxia scores were calculated using a previously described method^45^. Gene markers of hypoxia identified by Buffa et al.^138^, Winter et al.^139^, Ragnum et al.^140^, Elvidge et al.^141^, and Sorensen et al^142^. were synthesized into one gene list of 286 genes, and 28 genes filtered during the RNAseq processing pipeline were removed. The median tumor expression for each gene was calculated, and each tumor was given gene scores of +1 or -1 based on whether gene expression was above or below the median for each gene. The total hypoxia score for each tumor was then calculated from the sum of its gene scores, where +258 would be the most hypoxic score and -258 would be most normoxic.

### Replication timing for mutational signature ID2

The replication timing data for two lung cancer cell lines, A549 and NCI-H460, were obtained from Repli-Seq assays available in the ENCODE database (accession IDs: ENCFF454EFB for A549 and ENCFF098QZG for NCI-H460). For each chromosome, we calculated the median replication timing using the log2 ratio of early (E) to late (L) replicating regions. Additionally, we determined the number of ID2 deletions on each chromosome. To investigate the relationship between replication timing and ID2 mutational signature, we performed Pearson correlation analyses between log2(E/L) and number of ID2 deletions across all chromosomes.

### Targeted bisulfite sequencing of the L1HS promoter on chromosome 22q12.1

#### Sodium bisulfite conversion of DNA samples

The DNA samples from tumor or normal tissues (29 tumor and normal pairs) were treated with sodium bisulfite to convert unmethylated cytosines to uracils. In addition, 6 standard human DNA samples containing 100% methylated, 66% methylated (in triplicate), 33% methylated, or 0% methylated human standard DNA from a commercial source (Human Methylated & Non-methylated DNA Set, Zymo Research) were treated along with the experimental samples in order to check the accuracy of the method in detecting the percentage of CpG DNA methylation.

Sodium bisulfite treatment was performed on 120 ng of each experimental or standard DNA sample using the EpiTect Fast DNA Bisulfite kit (Qiagen), following the manufacturer’s instructions. The reaction was conducted by adding 85 ml of bisulfite solution and 35 ml of DNA protection buffer to the DNA sample, in a total reaction volume of 140 ml. The reaction was conducted on an Applyed Biosystems 9700 thermal cycler (cycles: 95°C 5 min, 60°C 20 min, 95°C 5 min, 60°C 20 min, hold 20°C). After bisulfite treatment, the bisulfite-converted DNA was cleaned-up by MiniElute Spin columns (Qiagen), following the manufacturer instructions, and eluted in 15 ml of buffer EB.

#### Bisulfite PCR amplification of the target region

Polymerase chain reaction was performed by preparing a 60 ml reaction mix containing 6 ml of purified bisulfite converted DNA, 0.2 mM (final concentration) of each dNTP, 1.2 units (0.6 ml) of Phusion U Hot Start DNA Polymerase (Thermo Fisher Scientific), 12 ml of 5x HF Phusion Buffer, 0.5 mM (final concentration) for each of the 2 primers (respectively, Chr22_Forw and L1_rev_upper_360). The total reaction volume of 60 ml was divided into 3 vials of approximately 20 ml and the PCR reaction was performed on an Applied-Biosystems 9700 thermal cycler (cycles: 98° 1min 30s; 42-44 cycles of 98° 30s, 59° 20s, 68° 1min 30s; final extension 68° 7 min). The sequence and target specificity of the forward and reverse primers are listed in **Supplementary Table 7**.

The PCR amplified samples were then purified using Magtivio MagSi-NGSPREP Plus magnetic beads (with bead ratio 1.8x the input sample volume). After binding the DNA to the beads, the beads were washed twice with 70% EtOH (500 ml). The purified DNA samples were eluted in 50 ml of Low EDTA TE buffer (10 mM Tris, 0.1 mM EDTA, pH 8). Purification was performed in 1.5 ml Eppendorf LoBind tubes. The size and concentration of the amplicons were checked by automated electroforetic analysis on an Agilent Tapestation instrument, D1000 ScreenTape Assay.

#### Targeted next-generation bisulfite-sequencing

Approximately 70 ng of PCR amplified and purified DNA sample was used for NGS library preparation using the Ion Plus Fragment Library Kit (Thermo Fisher Scientific). The manufacturer’s instructions for the preparation of amplicon libraries were followed. In summary, the DNA sample was subjected to an end-prep reaction in a total volume of 50 ml for 25 min at room temperature, then to a cleanup with 1.8x MagSi beads (as reported above) with final elution in 12.5 ml of IDTE. Finally, a ligation reaction was performed by adding 1 ml of Ion Torrent P1 Adapter, 1 ml of Ion Xpress Barcode (different for each sample) and 35.5 ml of a ligation mix (containing 5ml of 10x ligase buffer, 1ml dNTPs, 1ul ligase enzyme, 4ml Nick repair polymerase and 24.5 ml H_2_O) to the purified and pre-purified DNA sample (12.5 ml). The reaction was performed in an Applied Biosystems 9700 thermal cycler (hold: 25°C 15 min, 72°C 5 min, 4°C up to 1 h). After adapter/index ligation, the library was purified from the unligated adapters by a double round of purification: the first round in MagSi beads 1x (50 ml of sample and 50 ml of beads), two washes in 300 ml EtOh 70%, elution in 50 ml IDTE; the second purification in SPRI select beads 0.9x (50 ml of DNA sample + 45 ml of beads), with three washes in 300 ml EtOh 70%, elution in 20 ml Low EDTA TE buffer.

After checking the size and concentration of each library on the D1000 HS Screen Tape Assay (Agilent), a pool of 455 pM of each library was prepared. The library pool was then treated with a semi-nested PCR assay of few cycles, to produce shorter amplicons and to increase that target specificity. The PCR reaction was performed by using a forward primer specific for the target sequence and fused with a Ion Torrent P1 adapter (trP1_Alu_Met_Forw_3), and a reverse primer specific for the external part of the indexed adapter (trA). The sequence of the forward and reverse primers are reported in **Supplementary Table 7**.

The reaction was prepared by mixing 76.9 ml of Platinum SuperMix High Fidelity (Thermo Fisher Scientific), 0.5 mM forward primer and 0.5 mM reverse primer. The final volume of 100 ml was divided into two tubes (50 ml each) and the reaction was run in an Applied Biosystems 9700 thermal cycler (hold 95°C 5 min, 5 cycles at 95°C 15 sec, 55° for 30 sec, 70°C 1 min, hold 4°.

The riamplified library pool was than purified with 1.5x MagSi beads (2 washes with 70% EtOh, elution in 13 ml low TE). After checking the size and concentration of the riamplified library pool using the D1000 HS Screen Tape Assay (Agilent), it was diluted to 65 pM for loading on the Ion Chef instrument. Template preparation and chip loading were performed using the reagents included in the Ion 510™ & Ion 520™ & Ion 530™ - Chef Kit and an Ion 510 Chip (Thermo Fisher Scientific). The chip was loaded onto an Ion S5 instrument for NGS sequencing (Thermo Fisher Scientific).

After sequencing (850 run flows) and raw data processing, the sequences of each sample were mapped to the target region (bisulfite converted sequence) on the hg19 human genome sequence.

### Statistical analyses

All statistical analyses were performed using the R software (v4.1.2) (https://www.r-project.org/). In general, the Mann-Whitney (Wilcoxon rank sum) test was used for two-group continuous variables, and the two-sided Fisher exact test was applied for the enrichment analysis of two categorical variables. P <0.05 was considered statistically significant. If multiple testing was required, we applied the FDR correction based on the Benjamini–Hochberg method. For the survival analyses, a proportional-hazards model was used to investigate the associations between the presence of ID2 mutational signature and overall survival, adjusting for age, sex, and tumor stage.

## DATA AVAILABILITY

Normal and tumor-paired CRAM files and the methylation raw data (intensity idat files) for the 542 WGS subjects of the Sherlock-*Lung* study have been deposited in dbGaP under the accession numbers phs001697.v1.p1 and phs002992.v1.p1. For RNA-Seq data, the FASTQ files for the same subjects can be accessed through dbGaP under the accession number phs002346.v1.p1. Detailed access information for the publicly available multi-omics datasets, including 153 subjects, can be found in **Supplementary Table 1**.

## CODE AVAILABILITY

The WGS bioinformatics pipelines can be accessed at https://github.com/xtmgah/Sherlock-Lung. Battenberg SCNA calling algorithm can be found at https://github.com/Wedge-lab/battenberg. Dirichlet process-based method for the subclonal reconstruction of tumors can be found at https://github.com/Wedge-lab/dpclust. The bioinformatic pipeline for identifying transposable element insertion is available at https://gitlab.com/mobilegenomesgroup/TraFiC.

## ACKNOWLEDGEMENTS

This work was supported by the Intramural Research Program of the National Cancer Institute, US National Institute of Health (NIH) (project ZIACP101231 to MTL); by the NIH grants R01ES032547-01, R01CA269919-01, and 1U01CA290479-01 to LBA as well as by LBA’s Packard Fellowship for Science and Engineering. The research performed in LBA’s lab was also supported by UC San Diego Sanford Stem Cell Institute. The funders had no roles in study design, data collection and analysis, decision to publish, or preparation of the manuscript. MD-G fellowship, within the “Generación D” initiative, Red.es, Ministerio para la Transformación Digital y de la Función Pública, for talent attraction (C005/24-ED CV1), is funded by the European Union NextGenerationEU funds, through PRTR. The computational analyses reported in this manuscript have utilized the NIH high-performance Biowulf Cluster and the Triton Shared Computing Cluster at the San Diego Supercomputer Center of UC San Diego. We thank the study participants, Dr. Peter Kraft for his reviewing of the manuscript and insightful comments, and the staff at Westat Inc. for their assistance in collecting samples and corresponding clinical data.

Where authors are identified as personnel of the International Agency for Research on Cancer/World Health Organization, the authors alone are responsible for the views expressed in this article and they do not necessarily represent the decisions, policy or views of the International Agency for Research on Cancer /World Health Organization.

## AUTHOR CONTRIBUTIONS

Conceptualization, MTL, TZ; Methodology, TZ, LY, BZ, JS, LBA, DCW, MTL; Formal Analysis, TZ, WZ, CW, MD-G, PHH, JFMS, JPM, AK, AK, CH, LY, BZ, JS ; Findings’ validation work: JY, MCA, MCE, FM, KB, JC, KMJ; Pathology work: CL, MKB, WDT, LMS, PJ, RH, S-RY; Resources, MPW, KCL, CAH, C-YC, NEC, ACP, DC, ESE, JMS, MBS, SSY, MMa, JL, BS, AM, OS, DZ, IH, VJ, DM, SM, MS, YB, BEGR, DCC, VG, PB, GL, PH, NR, QL, MTL, SJC; Data Curation, TZ, JR, MMi, FC-M, MS, OL; Writing – Original Draft, TZ, MTL; Writing – Review & Editing, LY, BZ, JS, JC, TZ, MAN, DCW, SJC, LBA, MTL; Visualization: TZ, MTL; Supervision, MTL.

## SUPPLEMENTARY TABLE LEGENDS

**Supplementary Table 1**: Demographics, clinical and tumor characteristics of 524 LUADs included in the Sherlock-Lung study.

**Supplementary Table 2**: Proportion of point mutations assigned to each early and late clone clusters estimated by MutationTimeR.

**Supplementary Table 3**: Predicted tumor latency with 90% confidence interval and MRCA age.

**Supplementary Table 4**: Annotation of nonsynonymous mutations in predicted driver genes.

**Supplementary Table 5**: Result of mutational signature deconvolution using SigProfileExtractor.

**Supplementary Table 6**: Summary of transposable element insertions with retrotransposition source information for each sample.

**Supplementary Table 7.** List of primer oligonucleotides used for library preparation for targeted bisulfite NGS.

## FIGURE LEGENDS

**Supplementary Fig. 1**: **Power analysis for detecting a diverse clonal architecture. a**) The scatter plot illustrates the relationship between the number of reads per chromosome copy (NRPCC) and the total detected single nucleotide variants (SNVs). Our ability to detect subclones relies not on the number of identified SNVs, but on the number of reads per tumor chromosomal copy. NRPCC accounts for tumor purity, ploidy, and sequencing coverage. **b**) The minimum cancer cell fraction (CCF) of the detected clusters in each tumor is plotted against NRPCC. To mitigate biases, we exclusively considered tumors with NRPCC ≥ 10. In these tumors, our analysis is sufficiently powered to identify a subclone with a CCF ≥ 30%. The suggested NRPCC threshold is denoted by the dashed line.

**Supplementary Fig. 2: Detailed sample and data information for LUAD tumor evolution analysis. a**) Summary of multi-omics data stratified by group information. **b**) proportion of samples within each group. **c**) Ancestry inference based on comparisons with samples from the 1000 Genome Project.

**Supplementary Fig. 3: Detection of whole genome doubling (WGD) events**. a) Tumor samples with and without WGD (nWGD) are distinguished based on their ploidy and the fraction of the genome exhibiting loss of heterozygosity (LOH). The initial demarcation line between WGD and nWGD tumors, established by the PCAWG study^26^, is represented as y = 2.9 − 2×. **b**) The fraction of the autosomal genome with a major copy number (MCN) of two or greater displays a bimodal distribution, highlighting the distinct separation between WGD and nWGD tumors.

**Supplementary Fig. 4: Overview of the molecular timing distribution of copy number gains per chromosome**. a) Pie charts depict the distribution of the inferred mutation time for a given copy number gain in a group. Green denotes early clonal gains, with a gradient to purple for late gains. The size of each pie chart is proportional to the frequency of recurrence of each event. **b)** Comparison of the molecular timing of copy number gains on chromosomes 21 and 22 versus other chromosomes across all three groups. **c)** Comparison of the molecular timing of copy number gains on chromosome X versus other chromosomes in tumors from the EU_S group. Two-sided Wilcoxon test p-values are shown above the box plots.

**Supplementary Fig. 5: Timing of point mutations reveals that most recurrent driver gene mutations occur early. a**) Driver mutations are categorized based on their mutation frequency, highlighting their distribution across clonality features. **b**) Driver mutations are presented as proportions, indicating their prevalence across clonality features.

**Supplementary Fig. 6**: **Contribution of mutational signatures to major driver genes. a**) Single base substitution (SBS) mutational patterns for the driver mutations of EGFR, KRAS, and TP53. **b**) Distribution of all driver mutations and hotspot mutations (recurrence > 5) in EGFR, KRAS, and TP53 attributed to each mutational signature.

**Supplementary Fig. 7: Mutually exclusive analysis of top mutated driver genes and exploration of the EGFR → TP53 evolutionary trajectory based on cancer cell fraction (CCF). a)** Oncoplot illustrating the mutational landscape across four major tumor groups (AS_N, EU_N, EU_S, and others) with frequencies displayed on the right side. Each colored band in the oncoplot represents a specific mutation type. **b)** Heatmaps summarize pairwise co-occurrence or mutual exclusivity (red gradient) relationships between driver gene pairs across the four tumor groups (AS_N, EU_N, EU_S, and others) and combined (ALL), assessed using Fisher’s exact test. The upper left triangle shows significant p-values on a -log10 scale, while the lower right triangle displays odds ratios (purple gradient, mutual exclusivity: OR < 1; co-occurrence: OR > 1). **c)** Examples of CCF distributions with marked locations of *TP53* and *EGFR* mutations. A barplot highlights the enriched *EGFR* → *TP53* evolutionary trajectory in EU_N tumors compared to AS_N (P = 0.05, Fisher’s exact test). To validate the *EGFR* → *TP53* trajectory difference between AS_N and EU_N, CCF were compared in tumors with detectable subclones (CCF < 0.8), approximating mutation order. An *EGFR* → *TP53* trajectory was defined when: (i) CCFEGFR - CCFTP53 > 0.2, or (ii) *EGFR* resides on the right side of the clonal peak (higher CCF) and *TP53* on the left side of the clonal peak or subclonal peak (lower CCF).

**Supplementary Fig. 8: APOBEC mutation clonality**. Fold changes between relative proportions of subclonal and clonal mutations attributed to APOBEC mutational signatures. Box plots demarcate the first and third quartiles of the distribution, with the median shown in the center and whiskers covering data within 1.5× the IQR from the box. Echo dot symbolizes a tumor sample, with its color indicating the APOBEC mutational ratio (calculated as APOBEC mutations divided by total mutations).

**Supplementary Fig. 9: Dynamic SBS mutational processes during clonal and subclonal tumor evolution across ancestry and smoking group**. Fold changes between relative proportions of clonal and subclonal mutations attributed to individual SBS mutational signatures. Points are coloured by mutational signature. P-values from the Wilcoxon rank-sum test are displayed on the bottom of the boxplots.

**Supplementary Fig. 10: Dynamic SBS mutational processes during early and late tumor evolution.** Fold changes between relative proportions of early and late clonal mutations attributed to individual SBS mutational signatures. Points are coloured by mutational signature. P-values from the Wilcoxon rank-sum test are displayed on the bottom of the boxplots.

**Supplementary Fig. 11**: Distribution of mutations assigned to different mutational signatures for each mutation clonality group.

**Supplementary Fig. 12: Dynamic indel mutational processes during tumor evolution.** Fold changes between relative proportions of clonal and subclonal mutations attributed to individual indel mutational signatures. Points are coloured by mutational signature. P-values from the Wilcoxon rank-sum test are displayed on the bottom of the boxplots.

**Supplementary Fig. 13: Illustration of tumor latency between the most recent common ancestor (MRCA) and the age at diagnosis, based on the canonical tumor evolution model.** This figure depicts the timespan (orange double-sided arrow) between the emergence of a tumor’s progenitor cells, originating from a single most recent common ancestor (MRCA), and the age at diagnosis. Tumor latency is characterized by the diversification of distinct subclones (colored teardrop shapes), driven by the accumulation of driver mutations in immediate progenitor cells. This process is influenced by competition among subclones under endogenous and exogenous selective pressures, leading to branching evolutionary trajectories that may result in cancer relapse or metastasis. Adapted from Yates and Campbell, *Nature Reviews Genetics*, 2012^143^.

**Supplementary Fig. 14: Association between the copy number and ITH-adjusted CpG>TpG mutation burden and age at diagnosis with loess fitting curves. a)** Overall; **b)** Stratified by ancestry and tobacco smoking groups.

**Supplementary Fig. 15: Chronological timing inference based on whole genome sequencing data. a**) Median latency periods between the most recent common ancestor (MRCA) and the last detectable subclone before diagnosis, categorized by various CpG>TpG mutation rate changes in AS_N, EU_N and EU_S. **b**) Comparison of estimated tumor latency based on a 1x acceleration rate among AS_N, EU_N and EU_S. **c**) Comparison of age at diagnosis among AS_N, EU_N and EU_S. **d**) Estimated Most Recent Common Ancestor (MRCA) age based on a 1x acceleration rate in AS_N, EU_N, EU_S, and “Others”. P-values from the Wilcoxon rank-sum test are presented above the respective boxplots.

**Supplementary Fig. 16: Associations between tumor latency and *EGFR* mutation status (a), *KRAS* mutation status (b) and sex (c), stratified by ancestry and tobacco smoking group.**

**Supplementary Fig. 17: Associations between tumor latency and the presence of mutational signatures (SBS, ID, and DBS).** Mutational signatures are arranged based on the median tumor latency difference between tumors with and without each mutational signature. The bar plot at the bottom illustrates the significance of these associations determined by the Wilcoxon rank-sum test.

**Supplementary Fig. 18: Pearson correlation between the estimated tumor latency and the number of deletions attributed to mutational signature ID2.** Pearson correlation coefficients and corresponding p-values are displayed above the plots.

**Supplementary Fig. 19: Numbers of deletions per megabase attributed to the mutational signature ID2 across ancestry and tobacco smoking groups.** Each dot represents an individual tumor sample and only samples where ID2 signature is found are shown.

**Supplementary Fig. 20: Enrichment of mutational signature ID2 with late replication time.** The scatter plots illustrate the Pearson correlation between the number of deletions attributed to mutational signature ID2 on each chromosome and the median DNA replication time on the same chromosome, derived from two lung cancer cell line experiments (left: A549; right: NCI-H460). Pearson correlation coefficients and corresponding p-values are presented above the plots.

**Supplementary Fig. 21: Up-Regulation of cell proliferation marker genes in tumors and normal lung tissue exhibiting mutational signature ID2.** Boxplots depict the comparison of gene expression levels for each marker between samples with ID2 signatures and samples without, derived from both normal (top) and tumor (bottom) RNA-Seq data.

**Supplementary Fig. 22: Pearson correlation analysis of gene expression levels for cell proliferation markers and deletions attributed to mutational signature ID2 in RNA-Seq data from normal lung tissue.**

**Supplementary Fig. 23**: **Association between tumors with mutational signature ID2 and increased tumor proliferation. a**) Ingenuity Pathway Analysis (IPA) of differentially expressed genes in tumors with and without mutational signature ID2. The x-axis represents the Z-score, and the y-axis indicates the corresponding canonical pathways. Suppressed pathways are shown in blue, while activated pathways are shown in orange. Circle size reflects the level of significance. **b–d**) Gene Set Enrichment Analysis (GSEA) highlights key proliferation-related pathways significantly associated with tumors harboring mutational signature ID2, including Hallmark Mitotic Spindle (**b**), Hallmark MYC Targets V2 (**c**), and Hallmark G2M Checkpoint (**d**). The normalized enrichment score (NES) and FDR-adjusted q-value are displayed at the bottom of each GSEA plot. **e-f**) Enrichment of tumor metastasis in tumors with mutational signature ID2, stratified by *TP53* mutation status. Odds ratios and p-values from the Fisher’s exact test are shown above the plot. **g-h**) Comparison of ID2 deletion burden between tumors with and without metastasis, stratified by *TP53* mutation status (**g**) or across all tumors (**h**).

**Supplementary Fig. 24: Association between mutational signature ID2 with *TP53* driver mutations. a)** Enrichment for the presence of mutational signature ID2 in tumors with *TP53* mutations across AS_N, EU_N, and EU_S, and “Others” groups. Odds ratios and p-values, calculated using Fisher’s exact test, are shown above the plots. **b)** Elevated levels of deletions attributed to mutational signature ID2 in tumors with *TP53* mutations. The significance of these associations is determined by the Wilcoxon rank-sum test.

**Supplementary Fig. 25: Association between mutational signature ID2 and genome instability.** Increased percentage of the genome altered by copy numbers in tumors exhibiting the ID2 signature.

**Supplementary Fig. 26: Box plot illustrating neoantigen predictions across all mutational signatures.** Neoantigen burden is normalized as the number of neoantigens per 1000 mutations.

**Supplementary Fig. 27: Enrichment of immune cell types in tumors exhibiting mutational signature ID2.** Volcano **(a)** and Box **(b)** plots illustrating the difference in immune cell score estimated by multiple RNA-seq-based methods for immune cell decomposition between tumors with and without the ID2 signature.

**Supplementary Fig. 28: Gene Set Enrichment Analysis (GSEA) of L1-regulated gene sets in tumors with and without the ID2 mutational signature.** This figure presents GSEA comparing tumors with and without the ID2 mutational signature for L1-regulated gene sets identified from two genome-wide CRISPR–Cas9 screens: **a)** PMID: 29211708 and **b)** PMID: 38849613. For context, enrichment levels were benchmarked against 50 curated hallmark pathways from the MSigDB gene sets and 658 pathways from the KEGG MEDICUS database. Normalized enrichment scores (NES) and FDR-adjusted q-values are displayed at the bottom of each GSEA enrichment plot.

**Supplementary Fig. 29: Enrichments of the presence of each mutational signature in tumors with somatic source L1 insertions.** The horizontal line indicates the significance threshold FDR < 0.05 in orange and <0.01 in red.

**Supplementary Fig. 30. Validation of the relationship between mutational signature ID2, L1 retrotransposition, and ZNF695 in an independent TCGA LUAD dataset. a**) Pearson correlation between the number of L1 element insertions and the number of mutations attributed to mutational signature ID2. **b**) Enrichment of *ZNF695* RNA expression in tumors with detected ID2 signature or L1 insertions. P-values from the Wilcoxon rank-sum test and fold change are shown on the top of the boxplot.

**Supplementary Fig. 31: Enrichment of mutational signature ID2 presence and total L1 insertions (a), transposed L1 insertions from germline source (b), and transposed L1 insertions from somatic source (c).** Odds ratios and p-values, calculated using Fisher’s exact test, are shown above the plots.

**Supplementary Fig. 32: Comparison of tumor latency between tumors with and without detection of L1 insertions, stratified by the source of L1 insertions (ALL tumors, germline source, and somatic source).** P-values, calculated using Wilcoxon-sum rank test, are shown above the plot.

**Supplementary Fig. 33: Analysis of differentially methylated CpG probes in tumors with and without mutational signature ID2.** Horizontal dashed lines indicate significance thresholds (FDR < 0.05 in orange and FDR < 0.01 in red).

**Supplementary Fig. 34: Association between DNA demethylation in the promoter of L1 from germline source in Chr22q12.1 and mutational signature ID2. a**) Comparison of DNA methylation levels between normal and tumor samples across the CpG islands in the L1 promoter. **b)** Comparison of DNA methylation levels between tumors with and without the ID2 signature, across the CpG islands in the L1 promoter. P-values, calculated using the Wilcoxon rank-sum test, are displayed at the bottom of the plots.

**Supplementary Fig. 35: Heatmap of locus-specific L1 RNA expression detected using RNA-Seq data from normal (a) and tumor samples (b).**

**Supplementary Fig. 36: Association between smoking status and total L1 RNA expression detected estimated from RNA-Seq data. a)** Between smokers and never smokers; **b)** Across current, former and never smokers.

**Supplementary Fig. 37: Pairwised Pearson correlation of the expression of 390 human KZFP protein-coding genes in LUAD from RNA-Seq data.**

**Supplementary Fig. 38: Analysis of differentially expressed KZFP protein coding genes between normal and tumor samples (a), and between smokers and never smokers in tumors (b).** Horizontal dashed lines represent significance thresholds (FDR < 0.05 in orange and FDR < 0.01 in red). The top 20 significant genes are labeled by gene name. **c**) RNA expression difference between tumor and normal samples for *ZNF483*, *ZNF658* and *ZNF695*. **d)** *ZNF695* RNA expression in normal and tumor tissues across the four groups.

**Supplementary Fig. 39: Multivariable analyses confirm *ZNF695* expression as the most significant factor associated with mutational signature ID2. a)** Logistic regression analysis assessing the association between *ZNF695* expression and mutational signature ID2 status (binary outcome: presence vs. absence), adjusted for potential confounders, including sex, *EGFR* mutation status, age, *TP53* mutation status, smoking history, and *KRAS* mutation status. **b)** Linear regression analysis evaluating the association between *ZNF695* expression and the number of mutations attributed to ID2, adjusted for the same covariates. Statistically significant associations (P < 0.05) are highlighted in red. Horizontal bars represent 95% confidence intervals.

**Supplementary Fig. 40: *ZNF695* upregulation in tumors and its association with L1 retrotransposition activity. a**) Analysis of differentially expressed KZFP genes between tumors with and without L1 insertions. Horizontal dashed lines represent significance thresholds (FDR < 0.05 in orange and FDR < 0.01 in red). The top 20 significant genes are annotated with gene names. **b**) Box plots illustrate the differential expression of *ZNF695* among normal tissue samples, tumors without somatic L1 insertions, and tumors with somatic L1 insertions. **c**) Pearson correlations between KZFP gene expression and number of somatic L1 insertions. Horizontal dashed lines indicate significance thresholds (FDR < 0.05 in orange and FDR < 0.01 in red). The top 20 significant genes are annotated with gene names. **d**) Correlation between *ZNF695* expression and the number of somatic L1 insertions. Pearson correlation coefficients and corresponding p-values are displayed above the plot.

**Supplementary Fig. 41: Pearson correlation between gene expression levels of different cell proliferation markers and *ZNF695* using tumor RNA-Seq data.** Pearson correlation coefficients and corresponding p-values are displayed above the plots.

**Supplementary Fig. 42: *ZNF695* expression in single cell RNA-Seq studies. a**) Identification of lung cell types with detectable *ZNF695* expression across multiple single cell types from RNA-Seq data in the CZ CELLxGENE database. The numbers on the right y-axis represent the number of cells identified from the single-cell studies. **b**) Enrichment of *ZNF695* expression in alveolar type 2 proliferating (AT2pro) cells from a normal lung single-nucleus multiome dataset. Dot plot visualizes the normalized RNA expression of *ZNF695* by cell type. The color and size of each dot correspond to the scaled average expression level and fraction of expressing cells, respectively. **c)** Enrichment of L1 RNA expression in cells with detectable *ZNF695* expression. P-values from the Wilcoxon rank-sum test are shown on the top of the boxplot.

**Supplementary Fig. 43: Differential gene expression analysis of 70 downstream target genes regulated by KRAB-ZFPs in tumors with and without signature ID2.** Of the 70 genes regulated by KRAB-ZFPs previously identified^83^, 47 genes (67%) were significantly differentially expressed. Two dashed horizontal lines represent the false discovery rate (FDR) thresholds: 0.05 (orange) and 0.01 (red).

**Supplementary Fig. 44. Isoform-specific expression of *ZNF695* and its association with mutational signature ID2. a**) IGV Sashimi plot for *ZNF695*, illustrating the expression profile in four distinct groups, including tumor with and without ID2 and normal lung tissue with and without ID2. The plot aggregates all sequencing reads mapped to *ZNF695* from samples within each group. Junction reads with less than 10% of the maximum exon sequencing coverage within *ZNF695* are filtered out. **b**) Splicing junction ratio estimation for *ZNF695* across all four groups. The junction ratio, calculated as the ratio of junction reads to the maximum exon depth, is depicted. Junction ratios less than 0.1 are indicated using a dashed line. A zoom-in view of sequencing coverage of the last exon of the canonical *ZNF695* isoform is shown in the bottom right. **c**) Summary information of *ZNF695* isoforms, providing an overview of the different isoforms and their respective characteristics. The canonical *ZNF695* isoform is highlighted in bold.

**Supplementary Fig. 45: Schematic model of canonical and non-canonical ZNF695 function in L1 regulation.** This schematic, adapted from Rosspopoff and Didier, *Trends Genet.*, 2023^84^, illustrates ZNF695-mediated regulation of L1 retrotransposons. In normal cells, canonical KZFP transcripts encode proteins with an N-terminal KRAB domain and a C-terminal zinc finger array. The zinc finger domain binds transposable elements (TEs), while the KRAB domain recruits a heterochromatin-inducing complex—including TRIM28 (KAP1), SETDB1, the NuRD complex, HP1, and DNMTs. This complex mediates H3K9 trimethylation and directs DNMTs to methylate CpG sites in L1 promoter regions, repressing L1 retrotransposition. In tumor cells, however, non-canonical *ZNF695* transcripts, lacking the zinc finger domain, predominate. This impairs TE binding and DNMT recruitment, leading to L1 promoter hypomethylation and subsequent L1 activation. Between the first and second strand synthesis by activated ORF2p (Baldwin et al., *Nature*, 2024)^67^, intermediates may be cleaved and integrated into the genome. Coupled with DNA repair mechanisms, this process may generate single-base-pair indels (e.g., ID1/ID2). This suggests a dominant-negative effect, where the balance between canonical and non-canonical transcripts governs L1 methylation status.

