## Supplementary Figures for "Deciphering lung adenocarcinoma evolution and the role of LINE-1 retrotransposition"

#### Supplementary Fig. 1

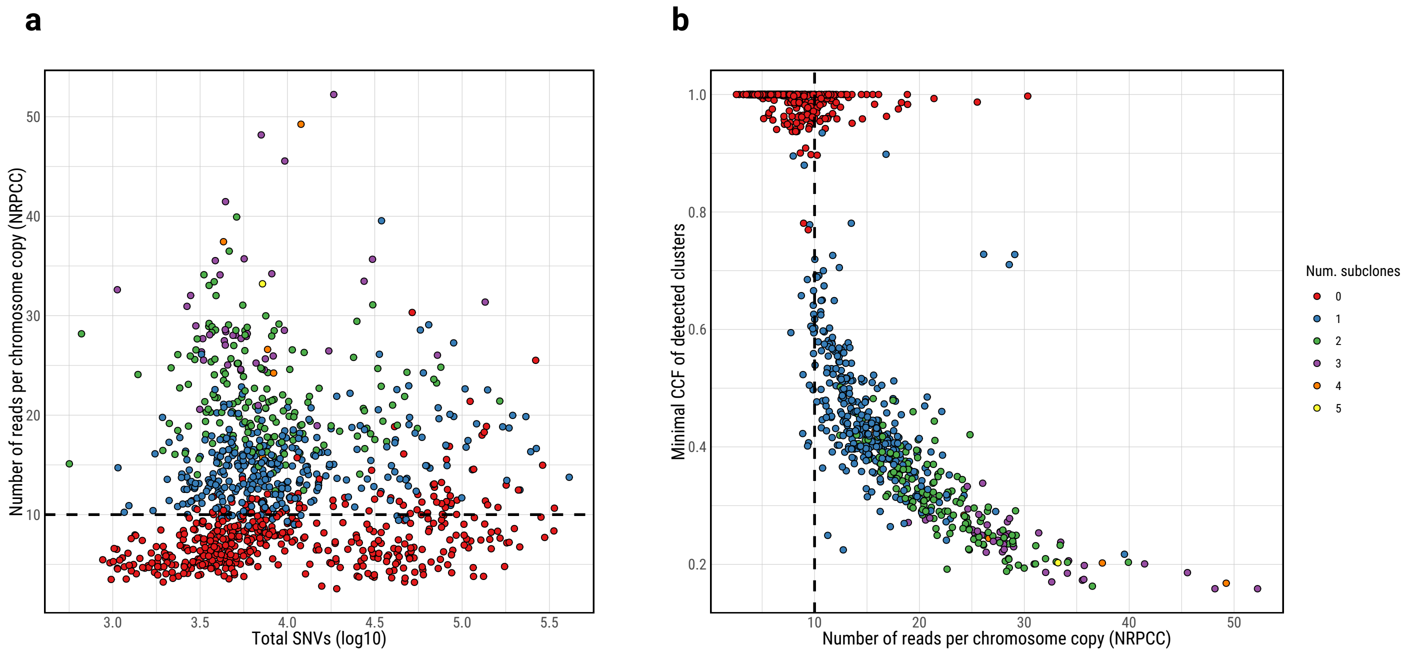

#### Supplementary Fig. 2

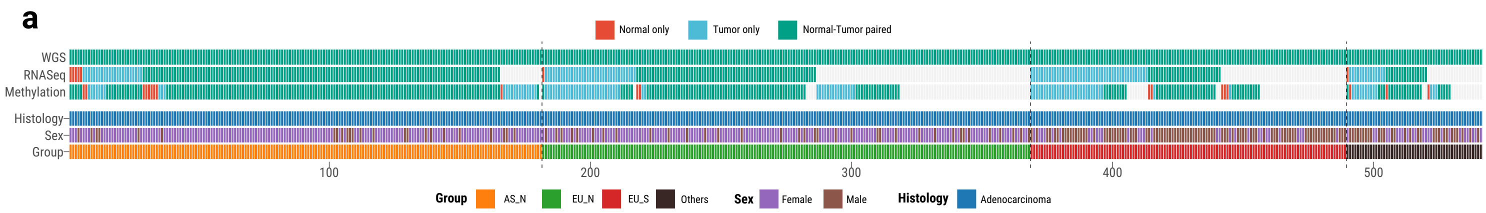


**b**

| Group | n | percent |
|---|---|---|
| AS_N | 180 | 33.2% |
| EU_N | 184 | 34.0% |
| EU_S | 120 | 22.1% |
| Others | 58 | 10.7% |

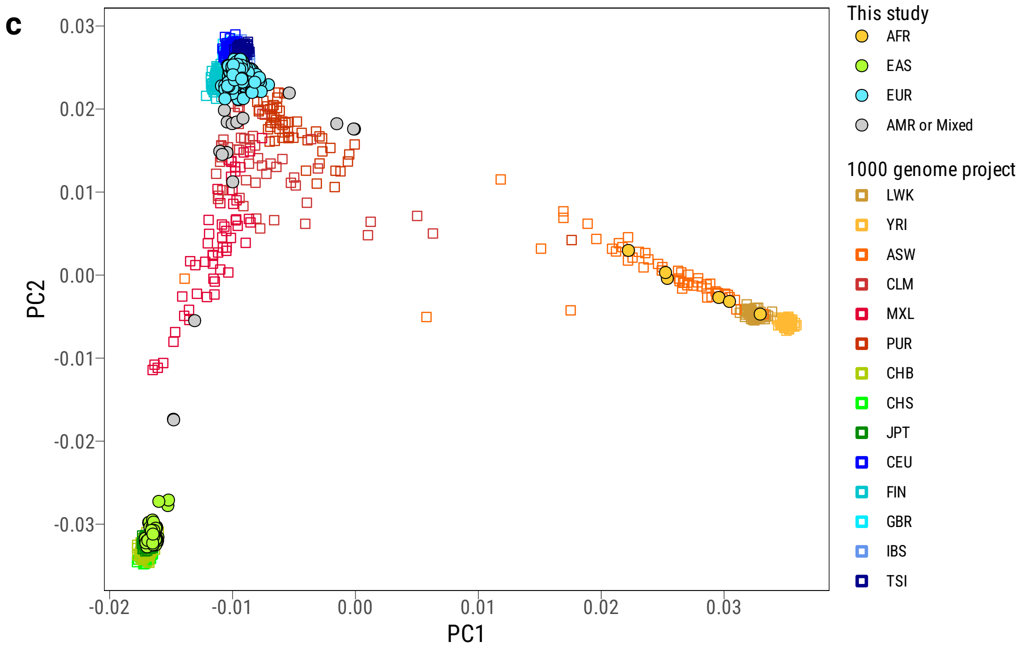

#### Supplementary Fig. 3

a

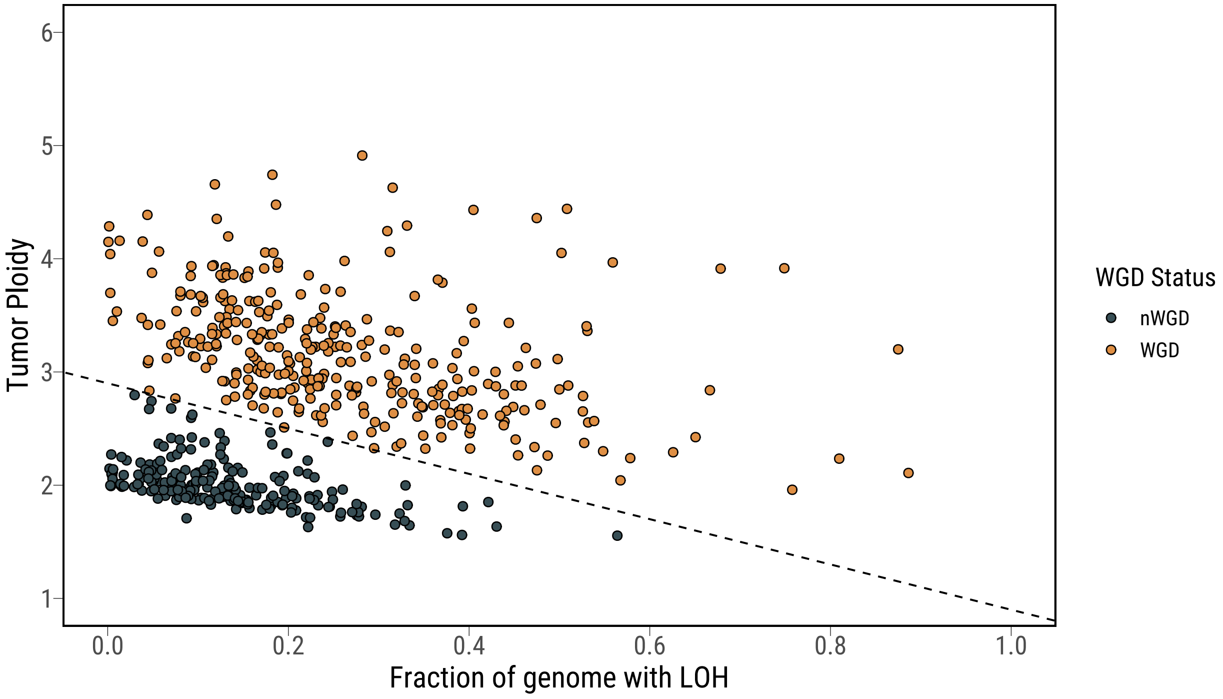


b

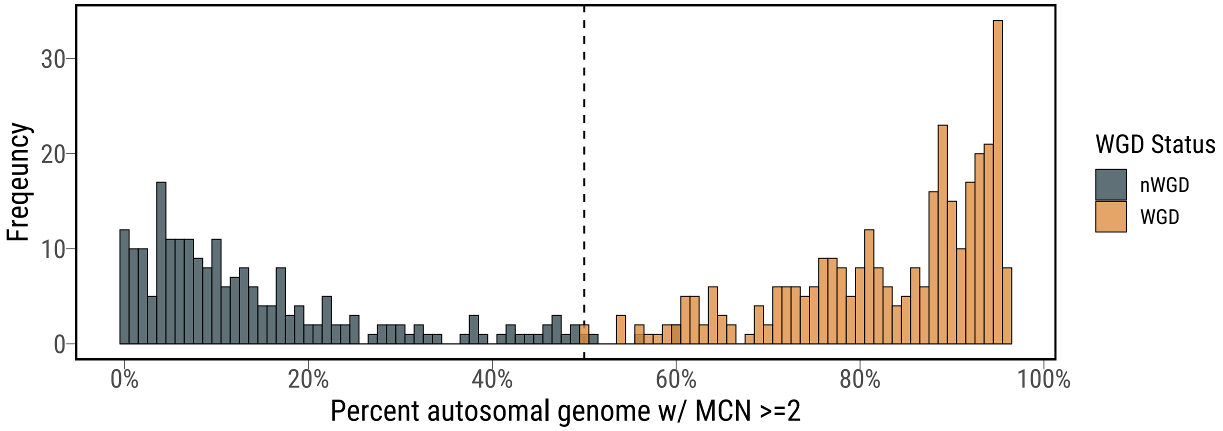

**Supplementary Fig. 4**

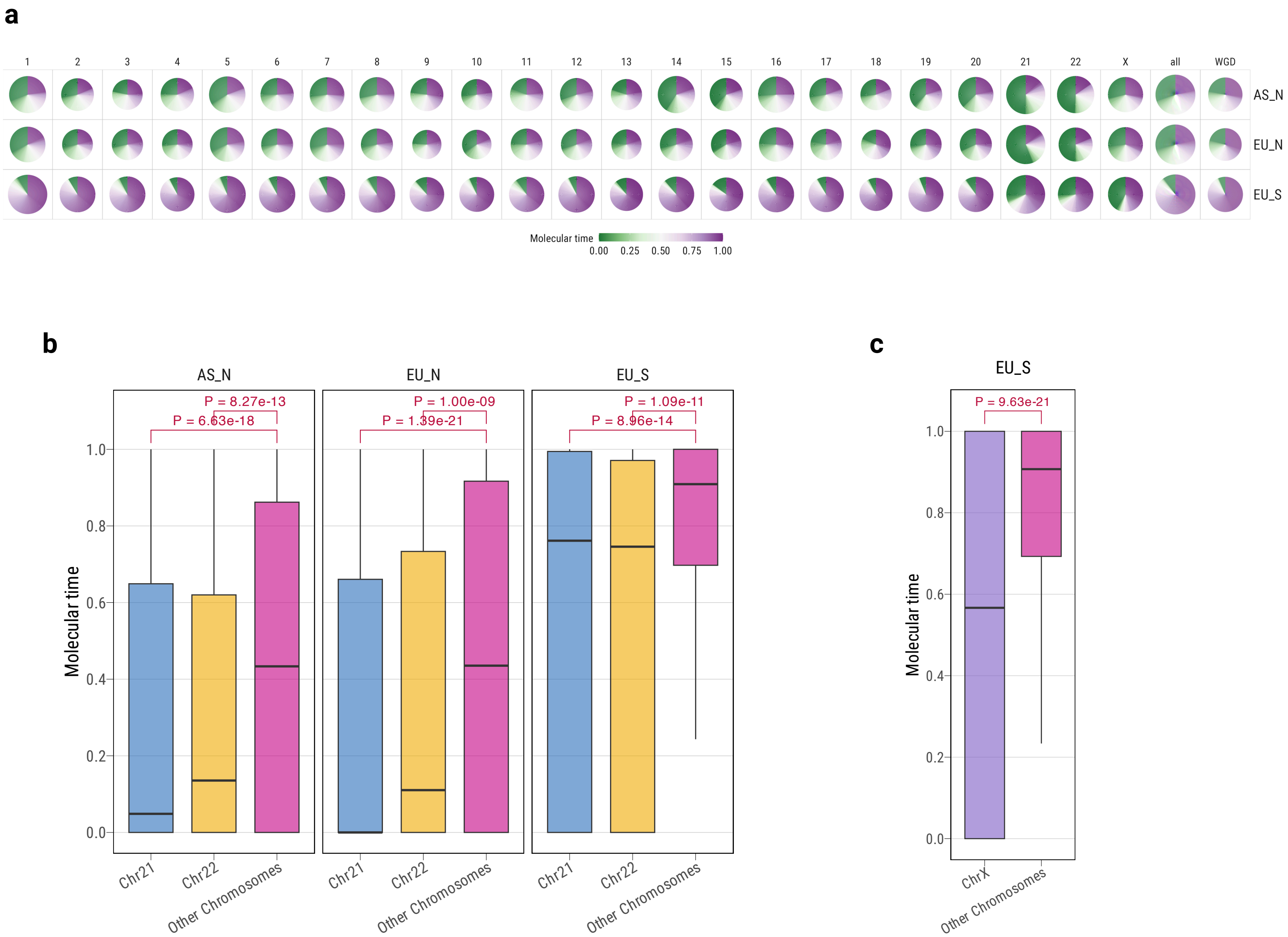

### Supplementary Fig. 5

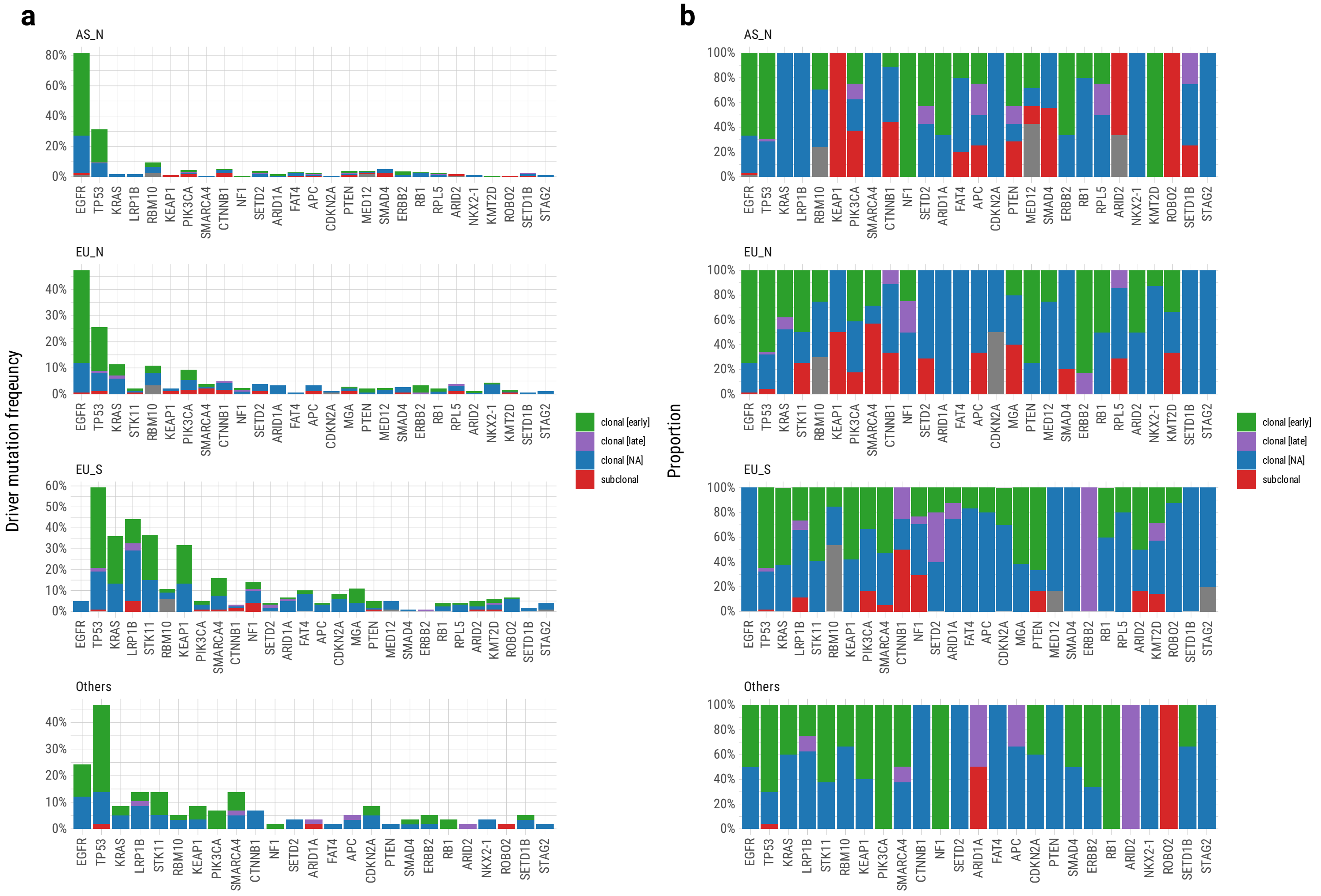

### Supplementary Fig. 6

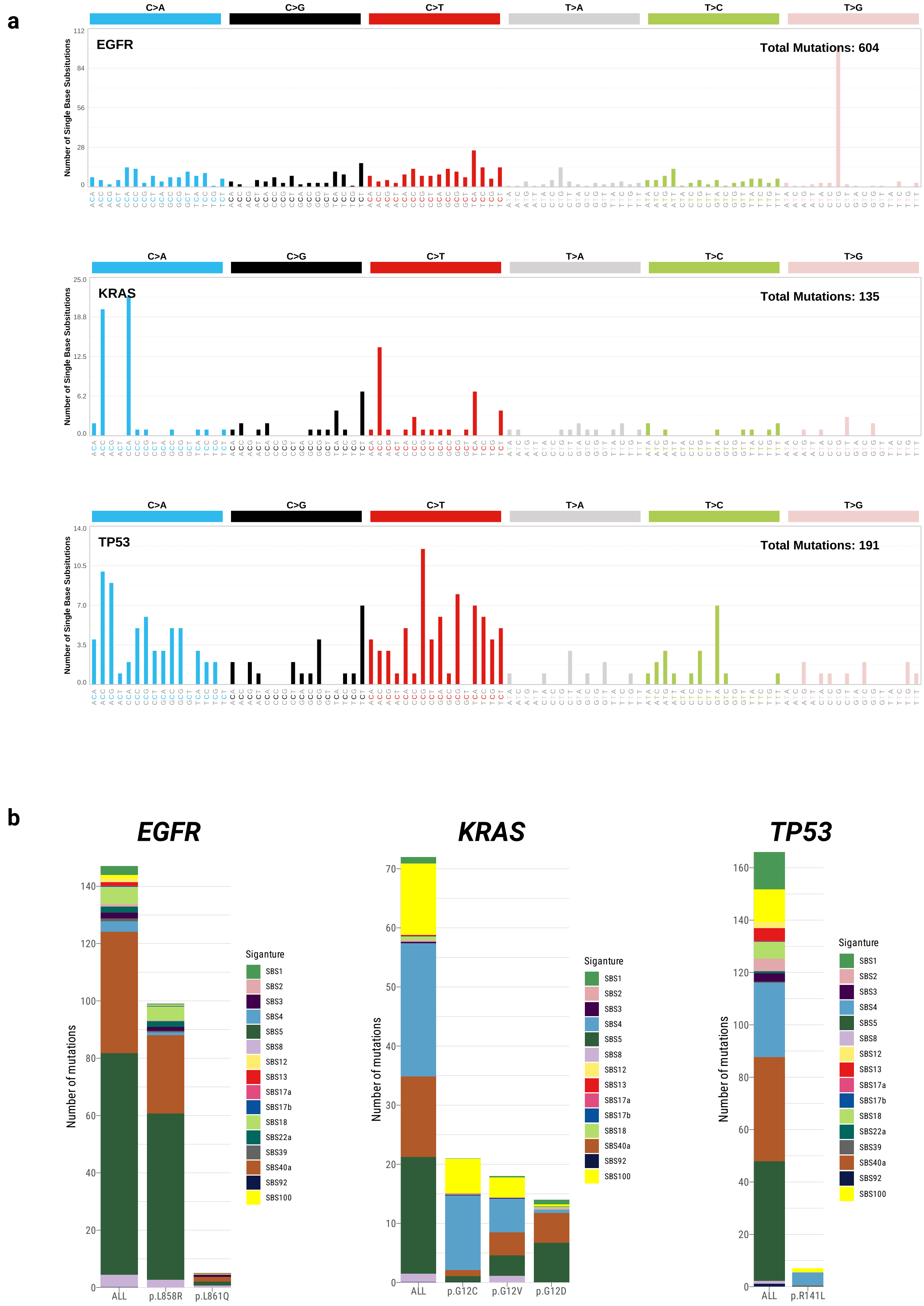

### Supplementary Fig. 7

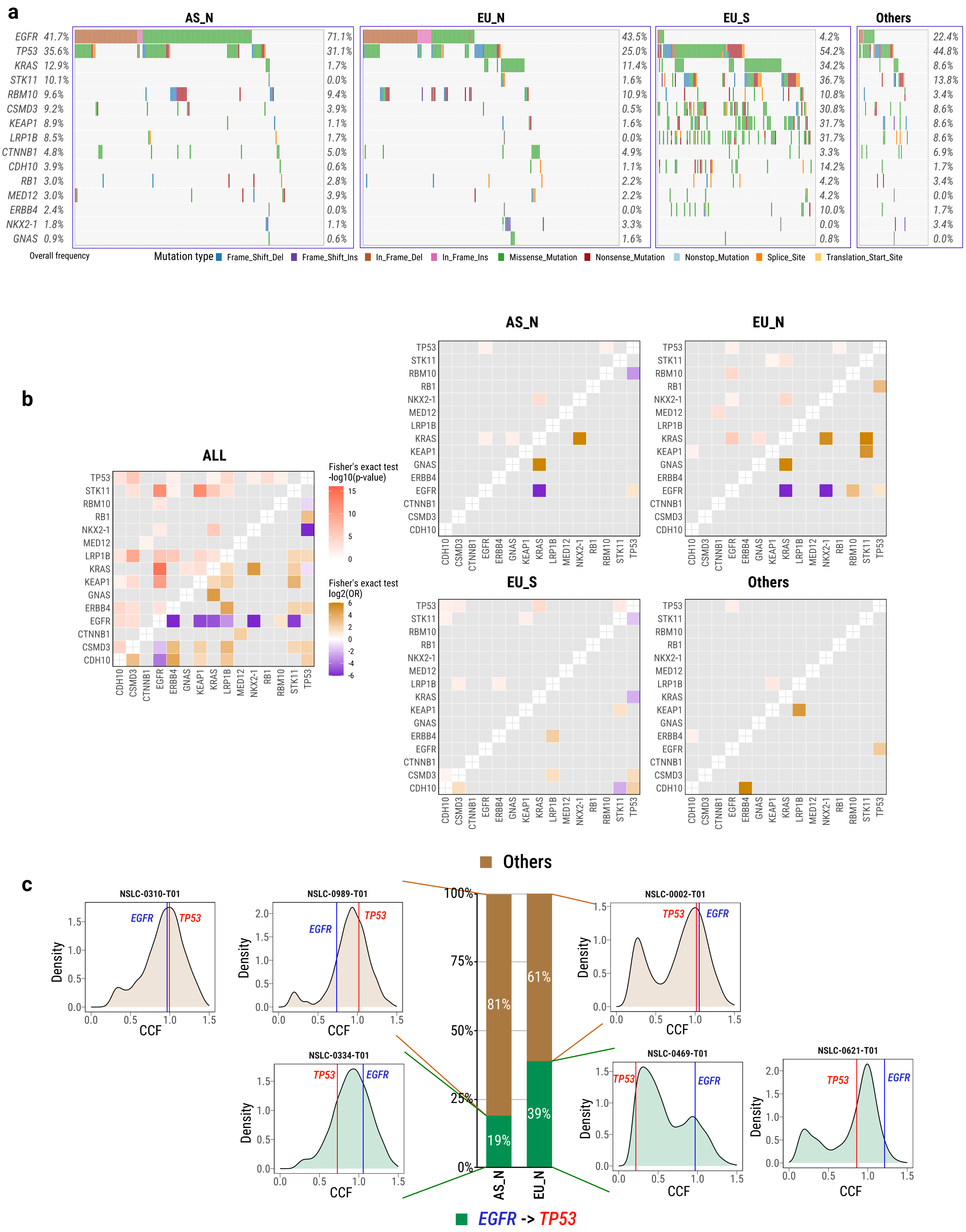

**Supplementary Fig. 8**

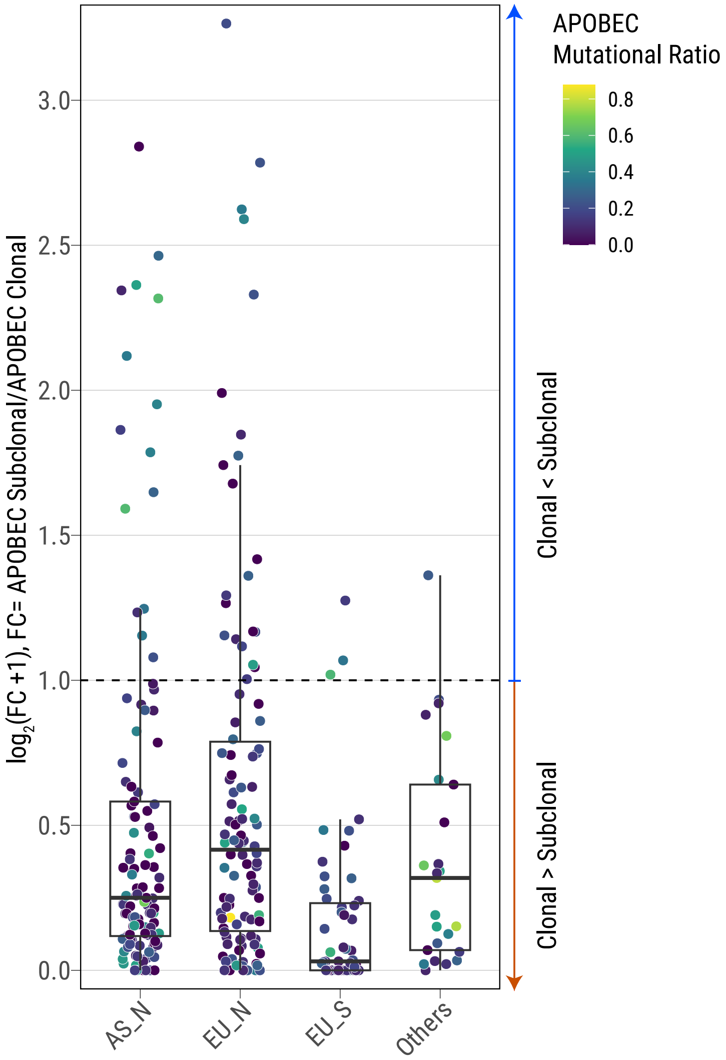

#### Supplementary Fig. 9

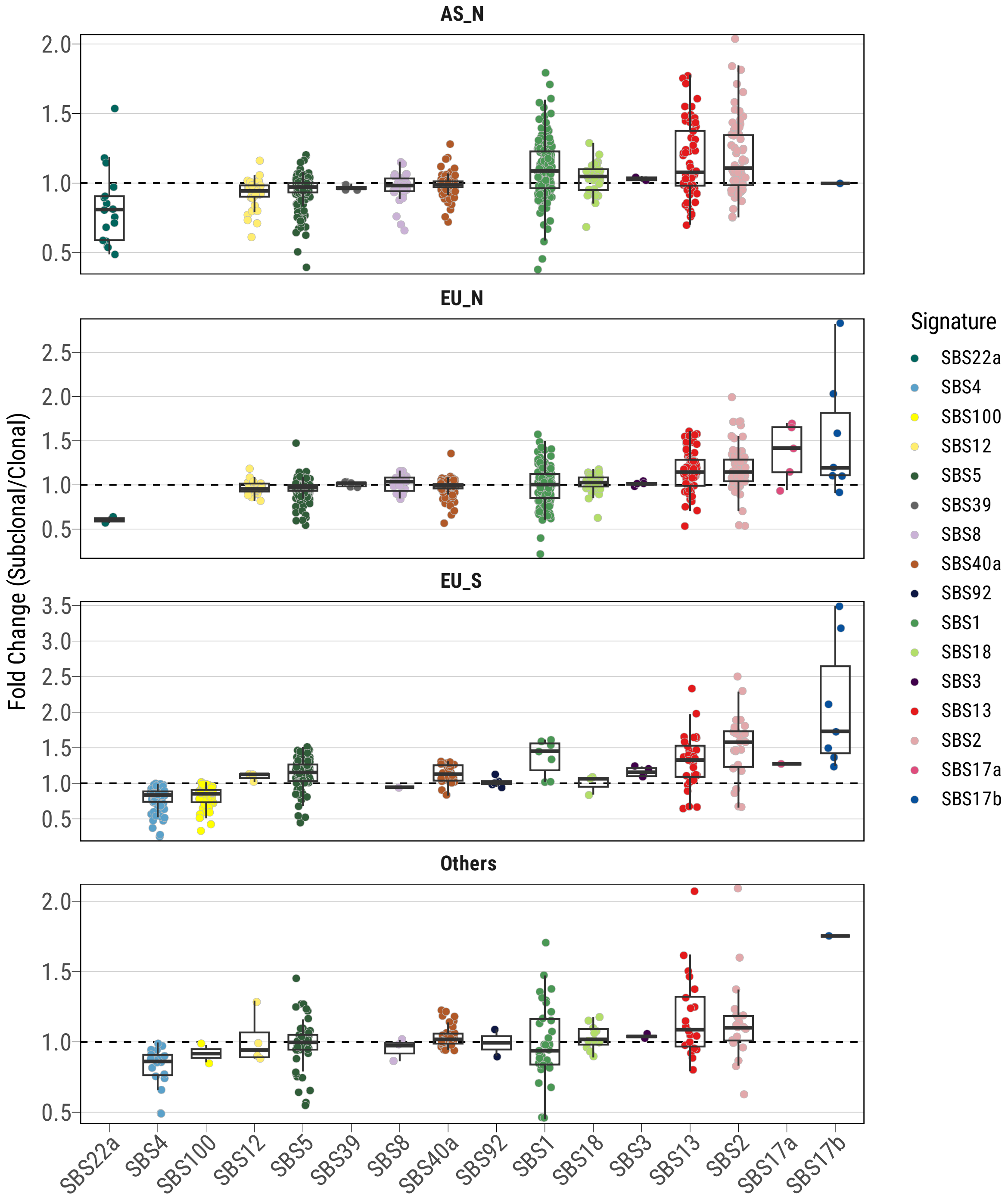

**Supplementary Fig. 10**

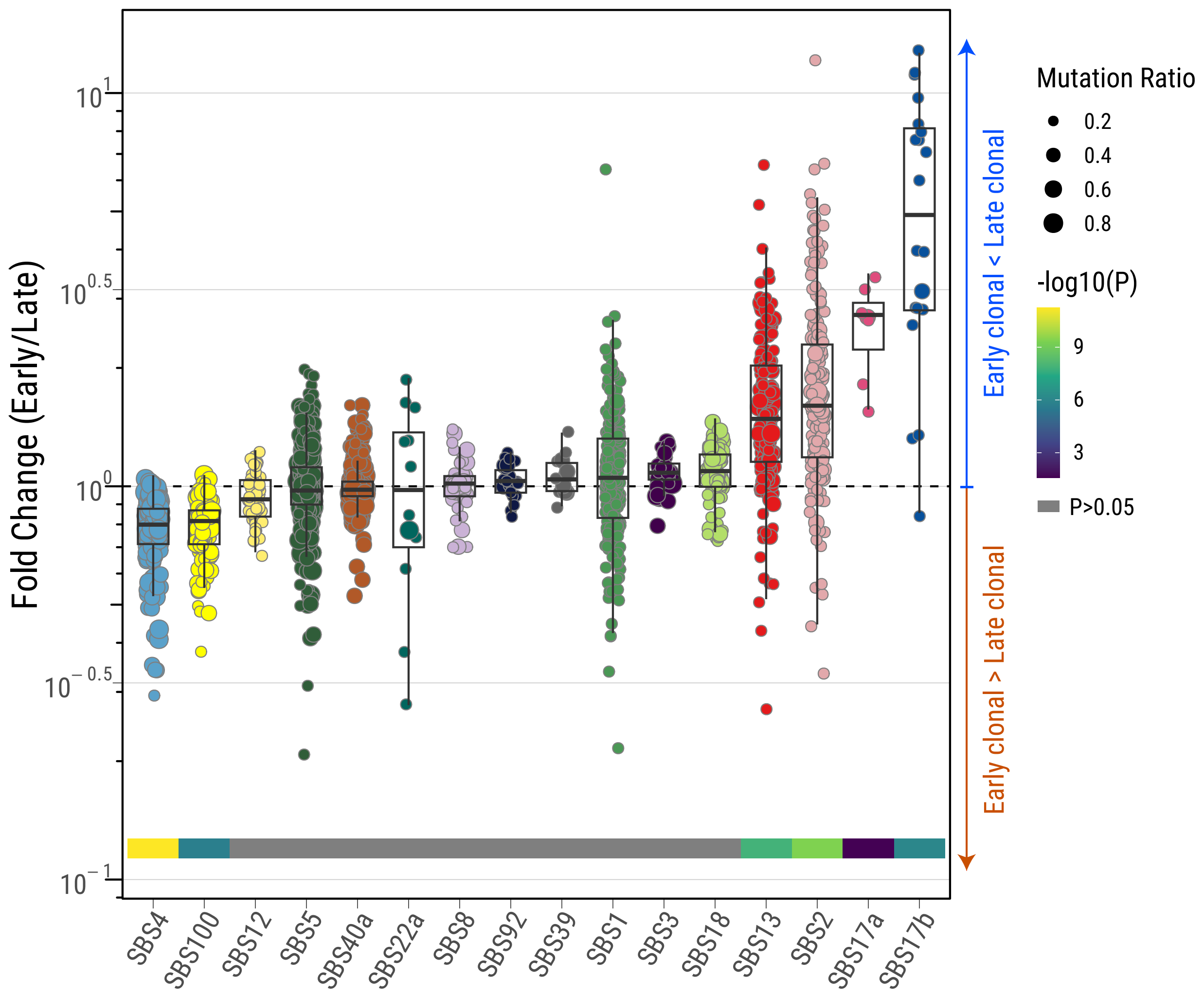

**Supplementary Fig. 11**

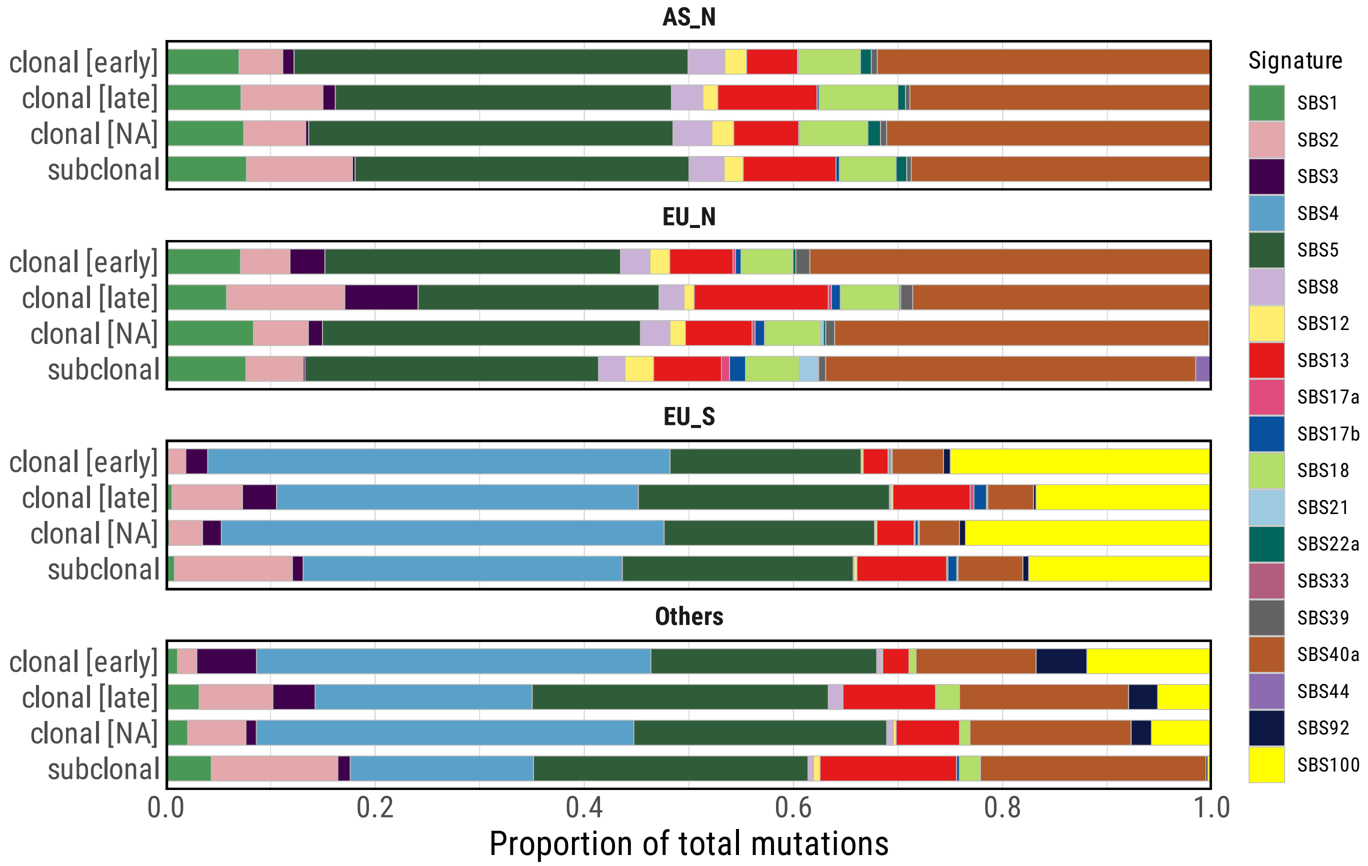

**Supplementary Fig. 12**

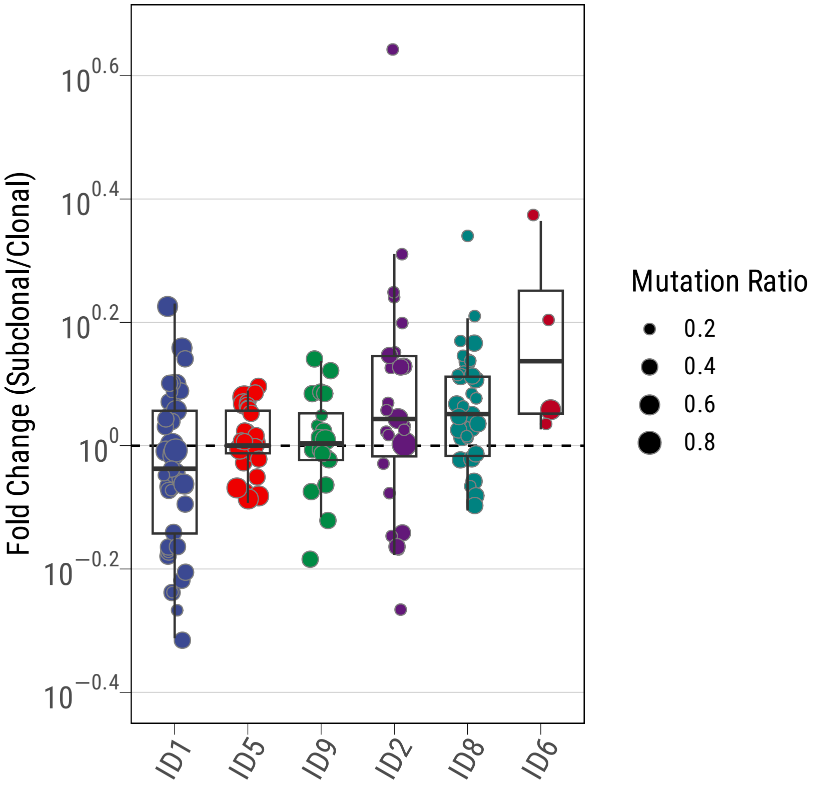

**Supplementary Fig. 13**

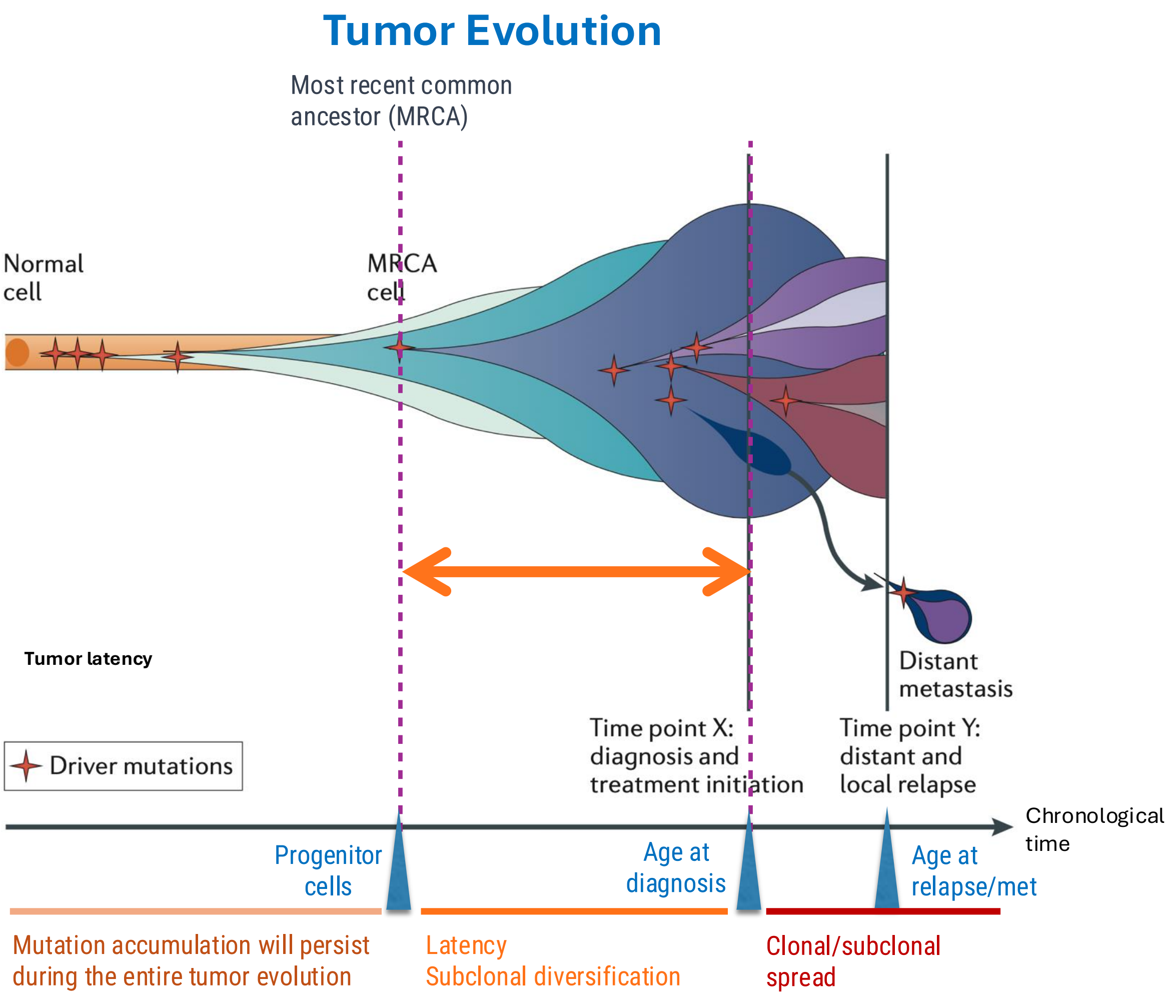


Figure adapted from Yates and Campbell, *Nature Reviews Genetics*, 2012

**Supplementary Fig. 14**

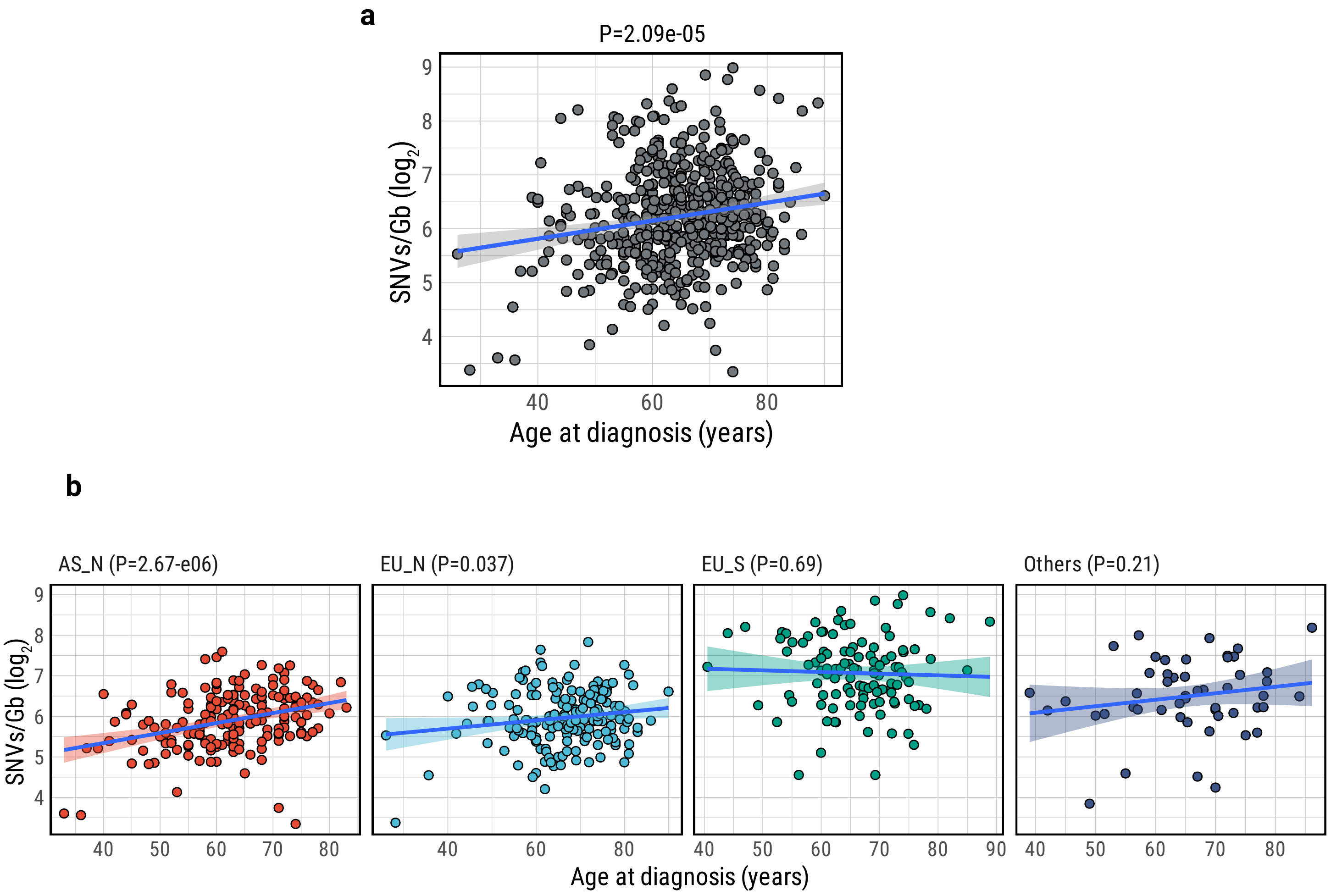

**Supplementary Fig. 15**

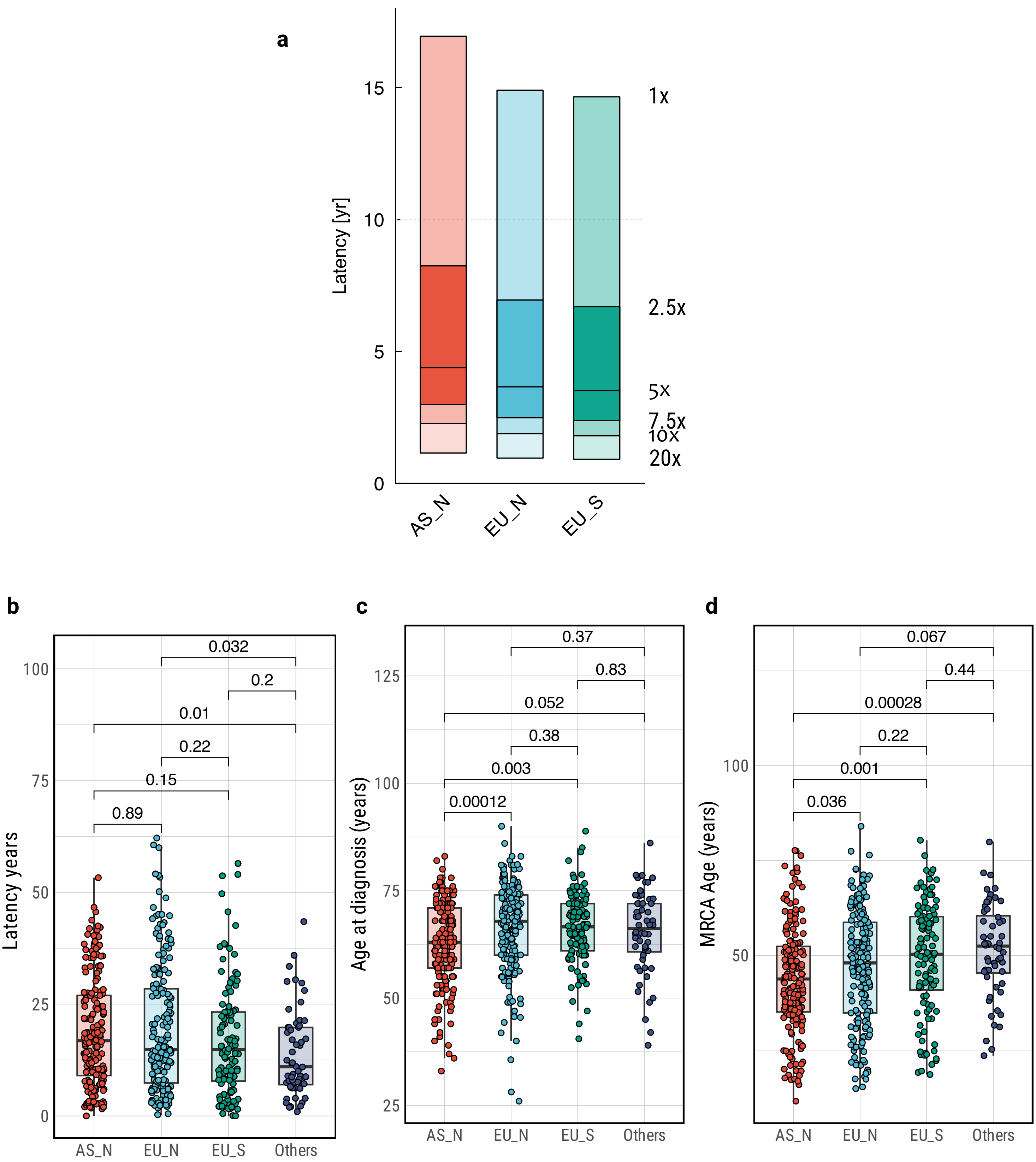

**Supplementary Fig. 16**

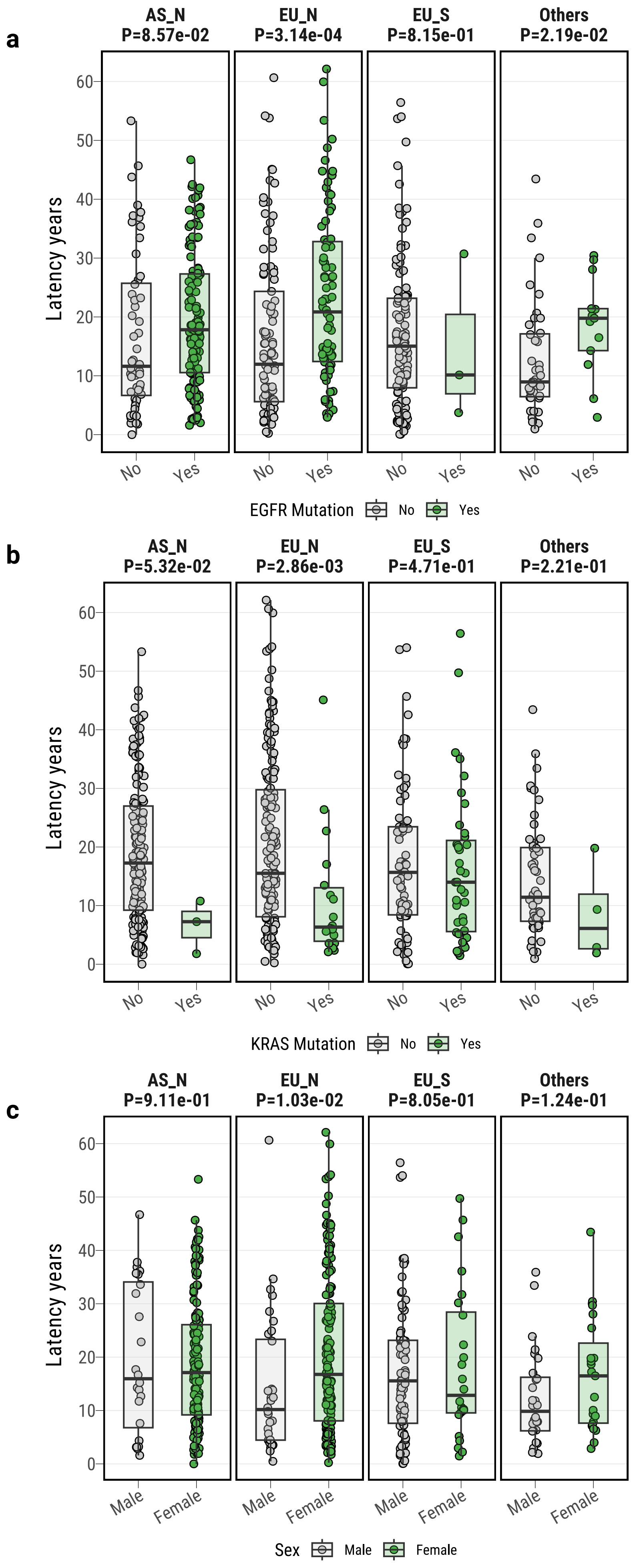

#### Supplementary Fig. 17

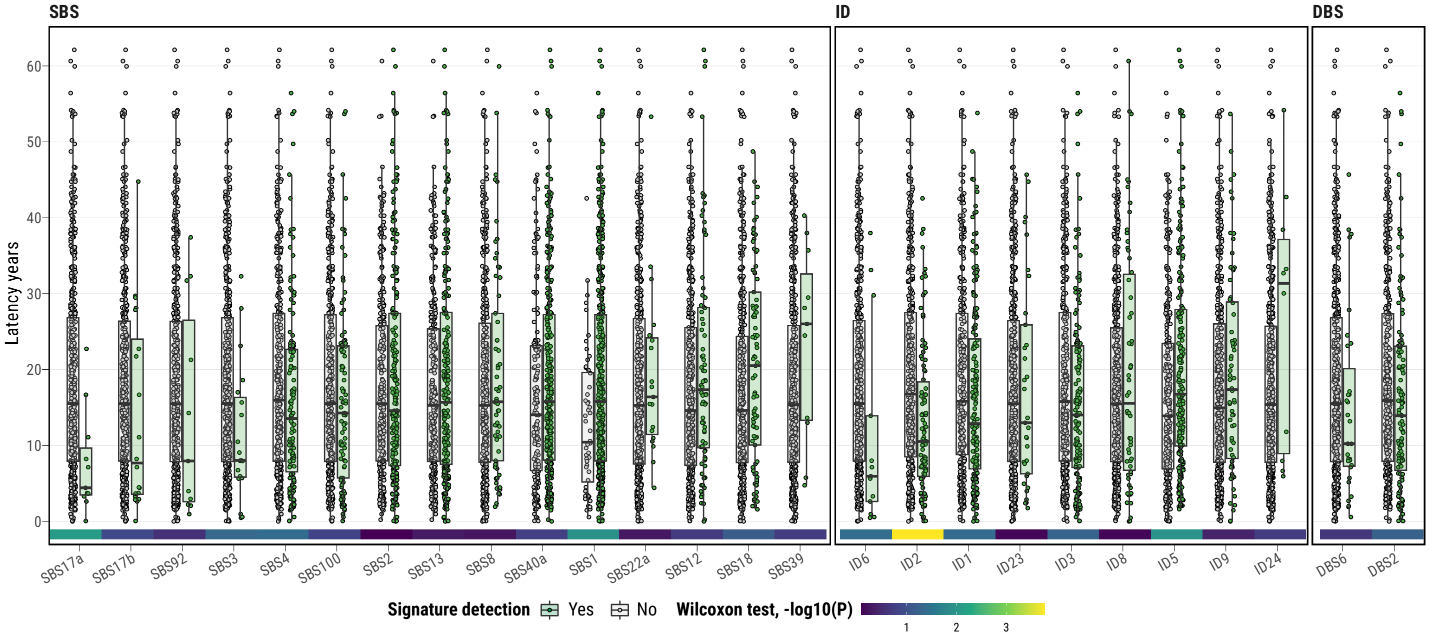

**Supplementary Fig. 18**

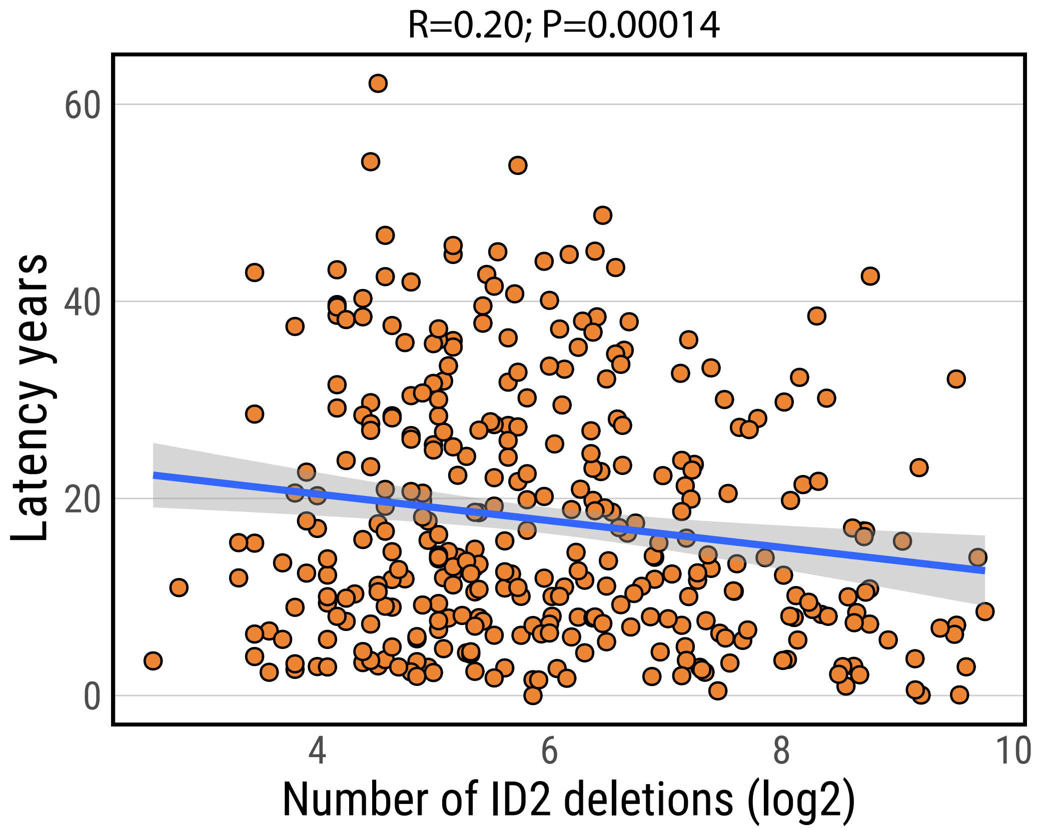

**Supplementary Fig. 19**

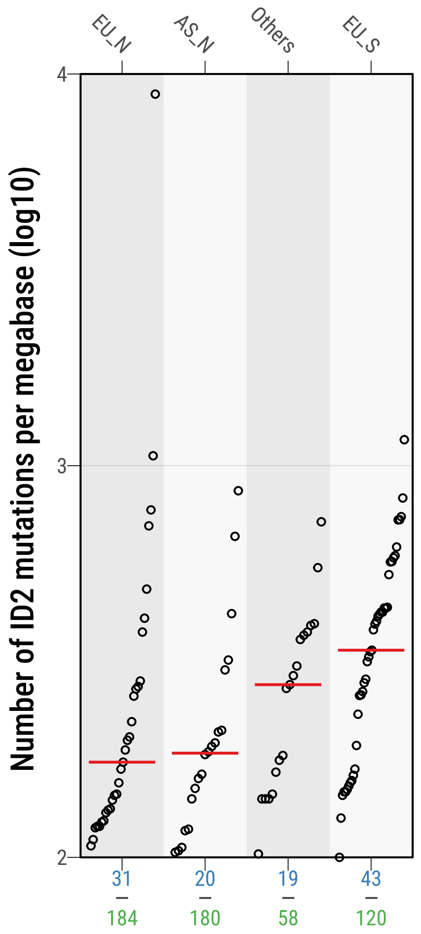

**Supplementary Fig. 20**

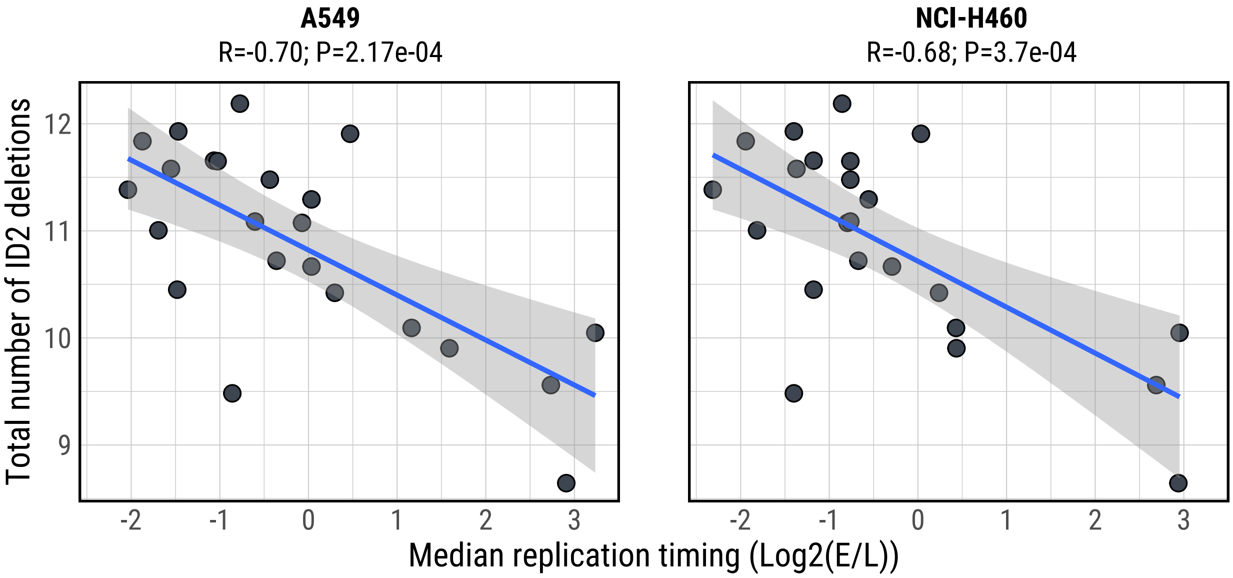

**Supplementary Fig. 21**

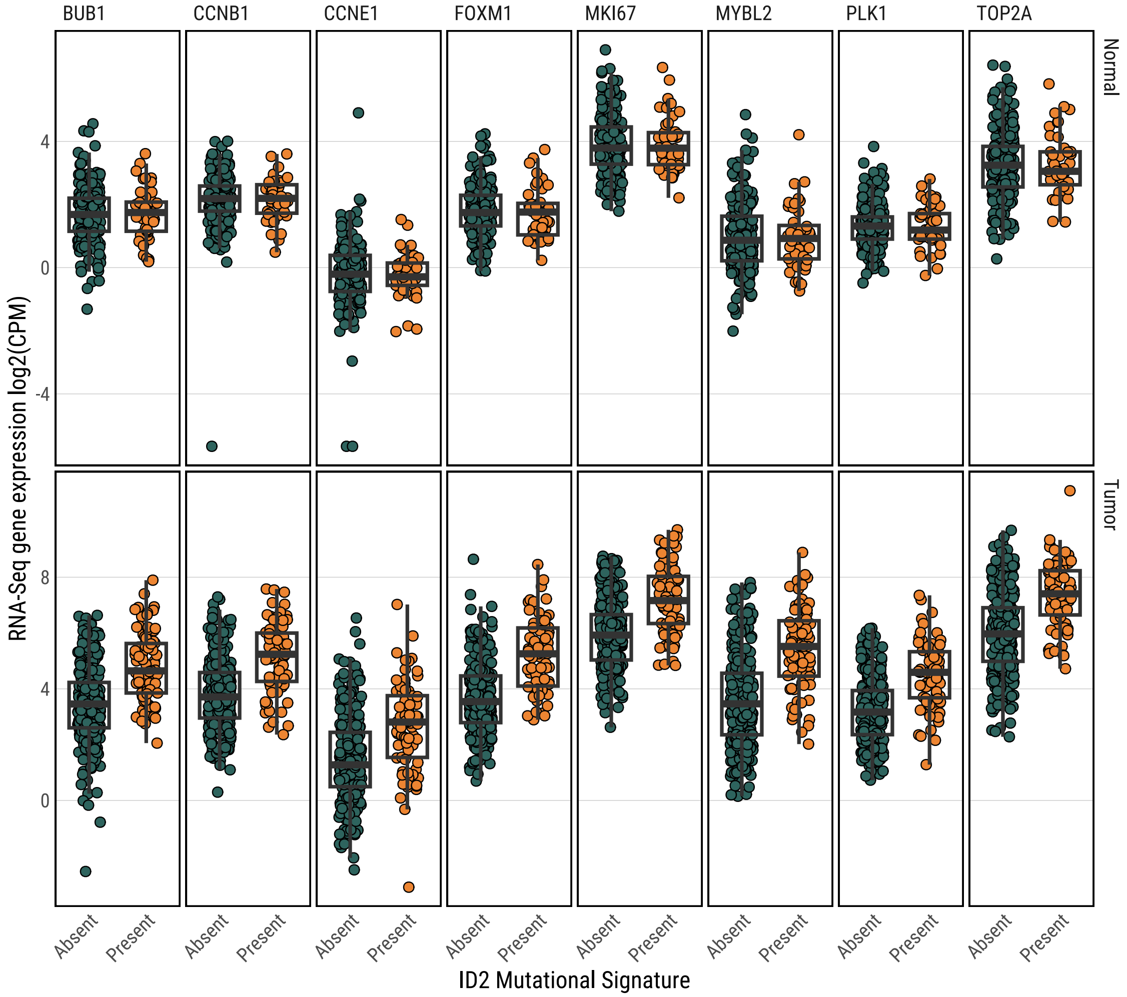

**Supplementary Fig. 22**

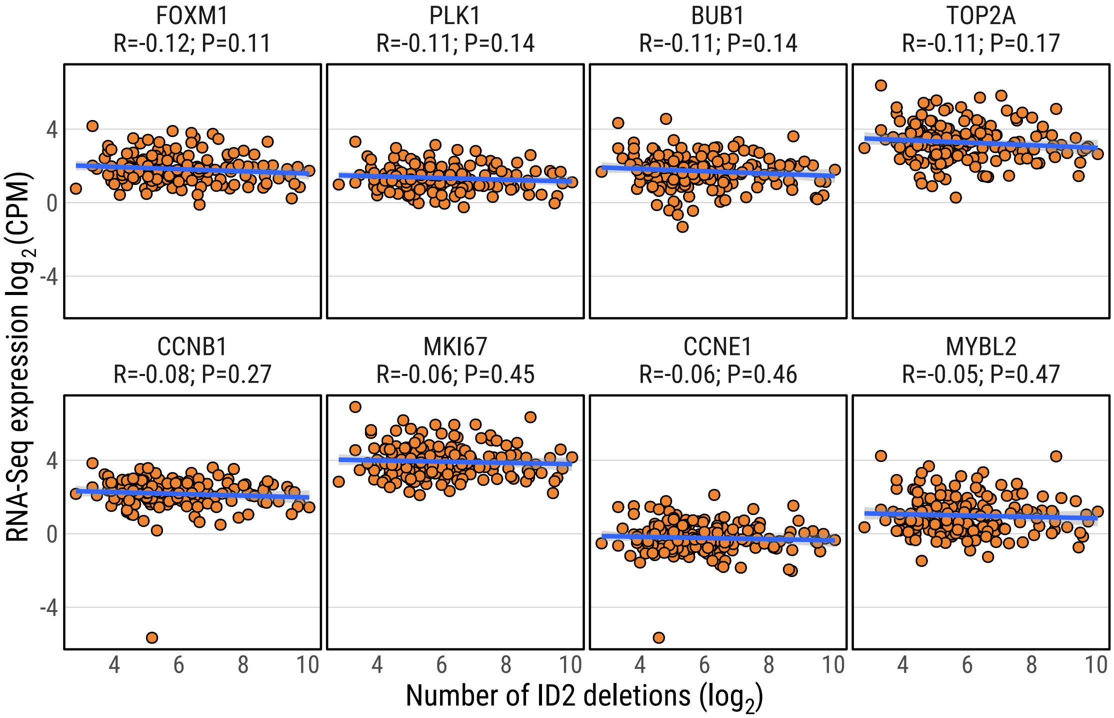

#### Supplementary Fig. 23

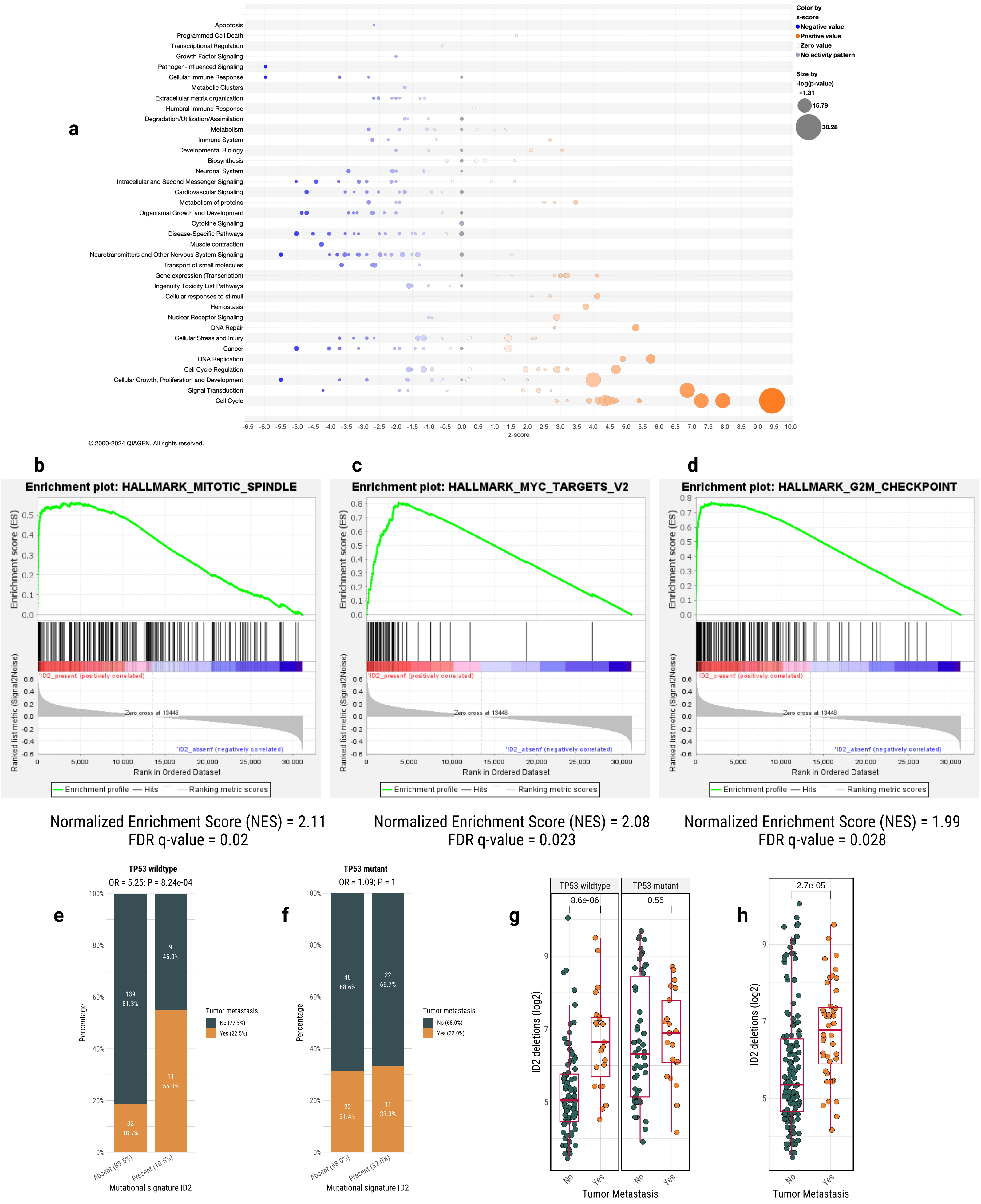

Supplementary Fig. 24

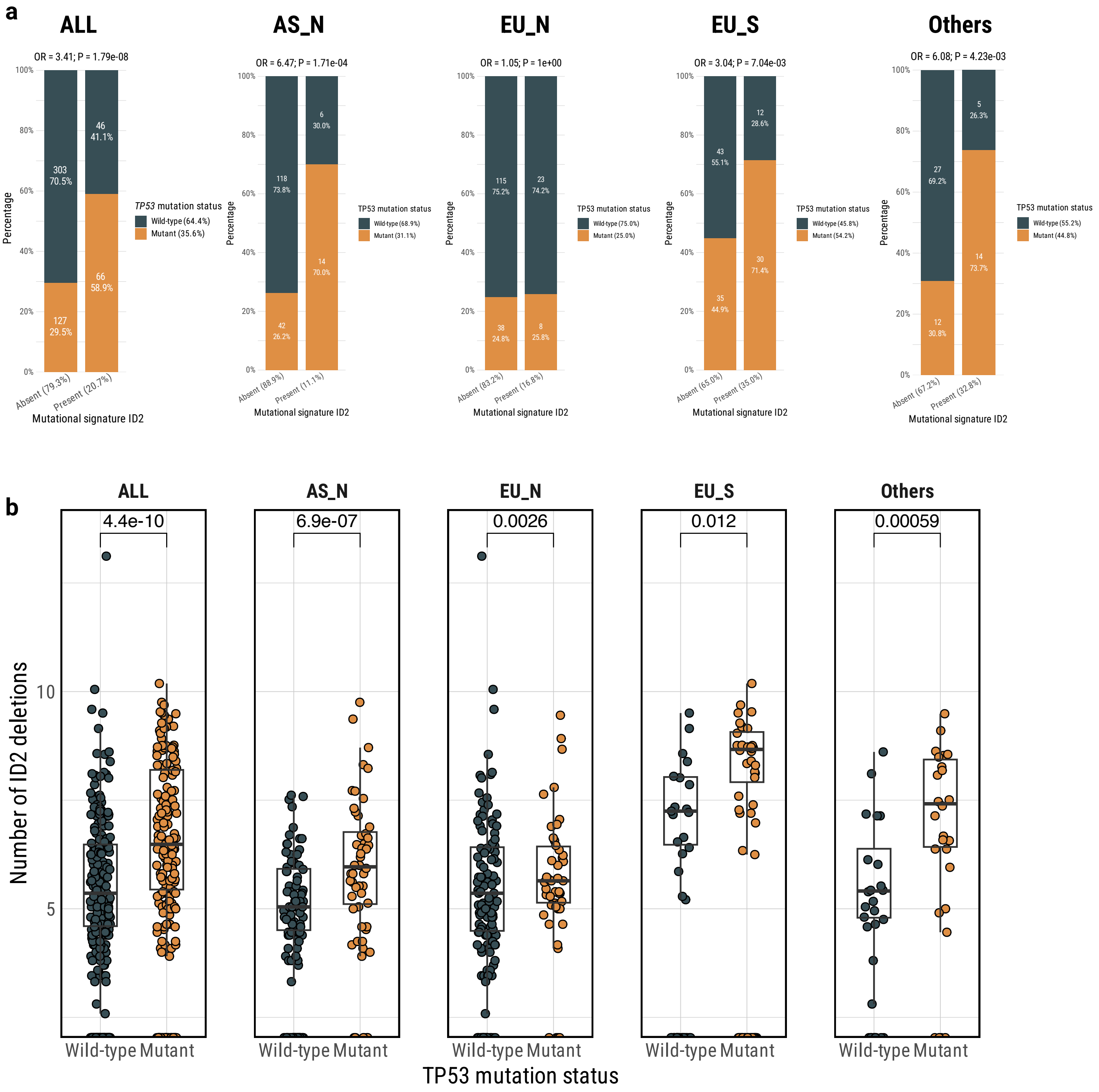

**Supplementary Fig. 25**

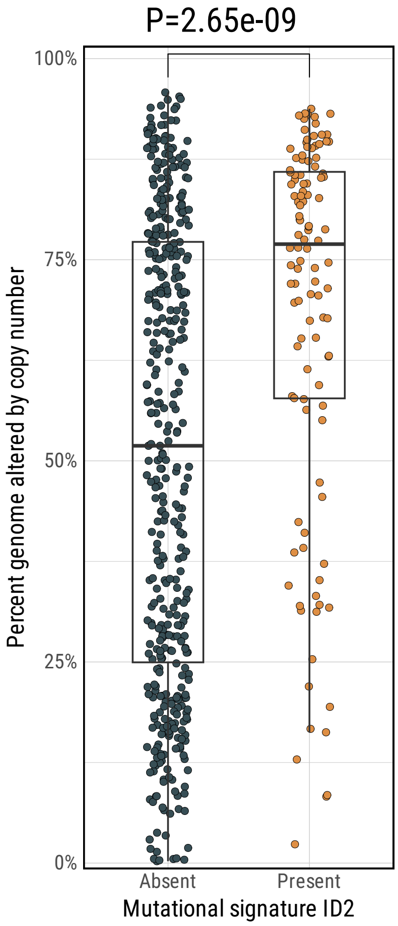

#### Supplementary Fig. 26

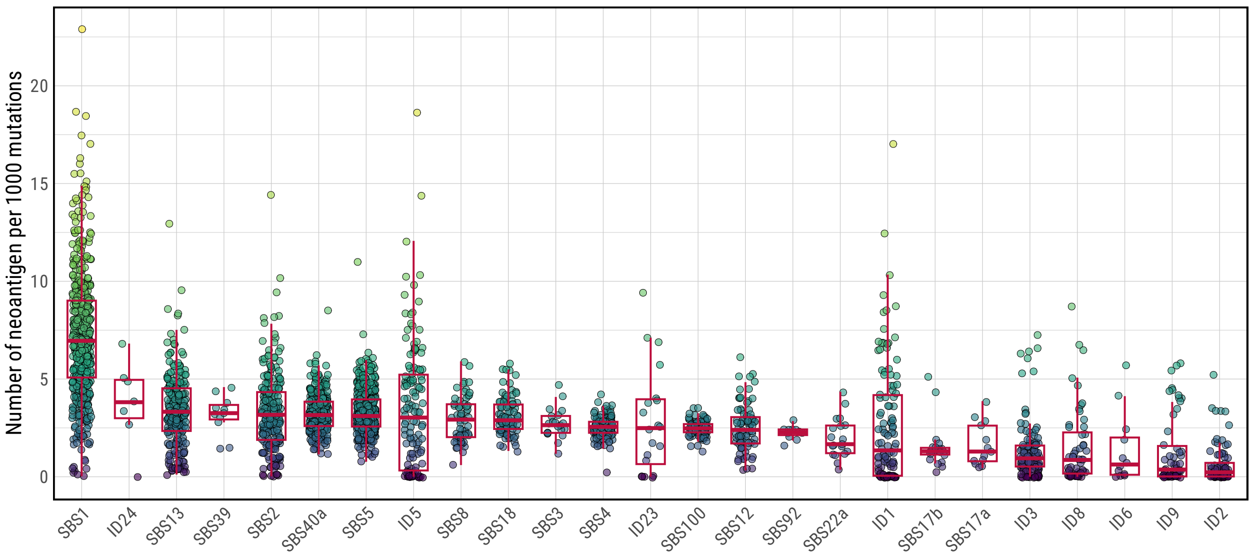

#### Supplementary Fig. 27

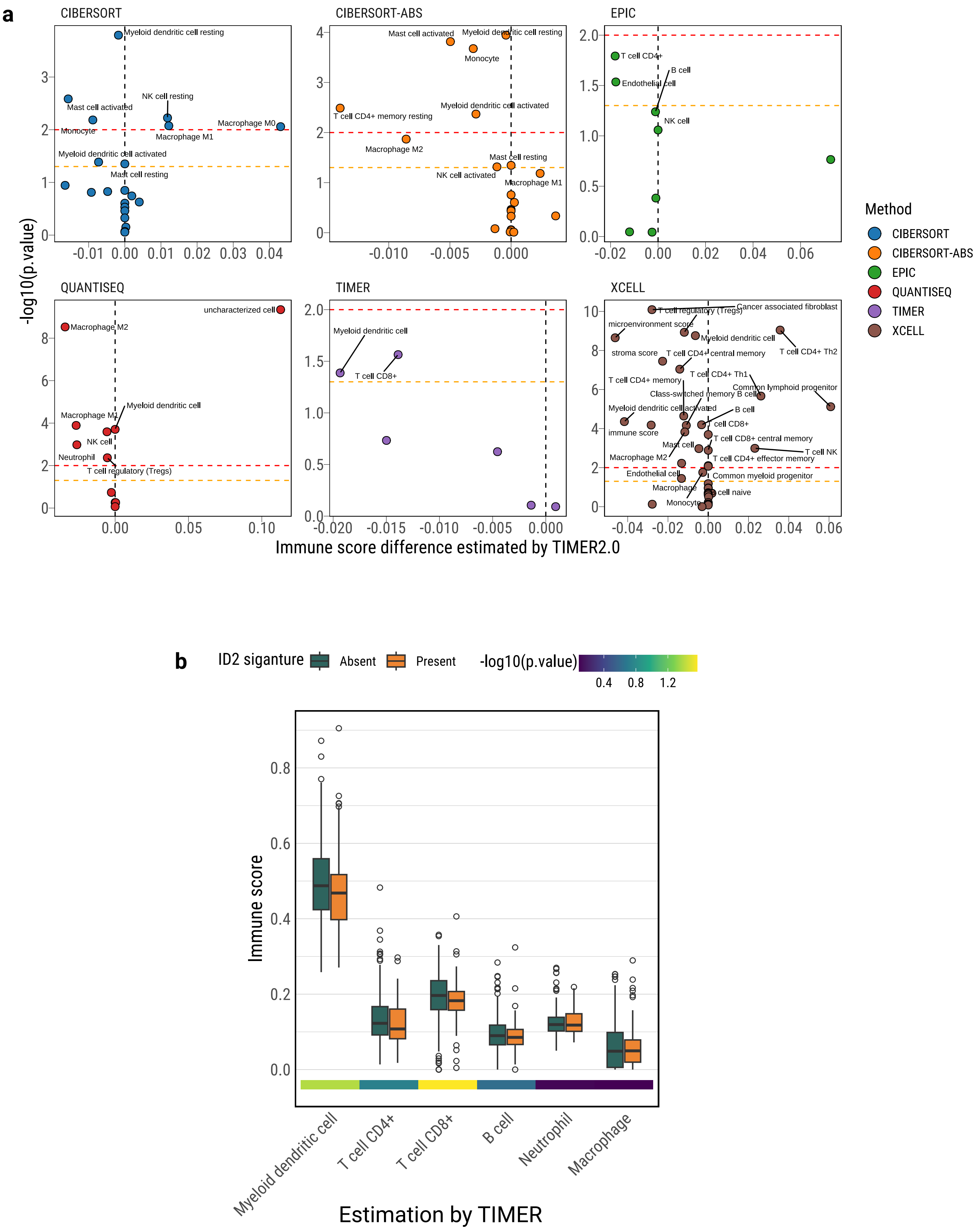

**Supplementary Fig. 28**

**a**

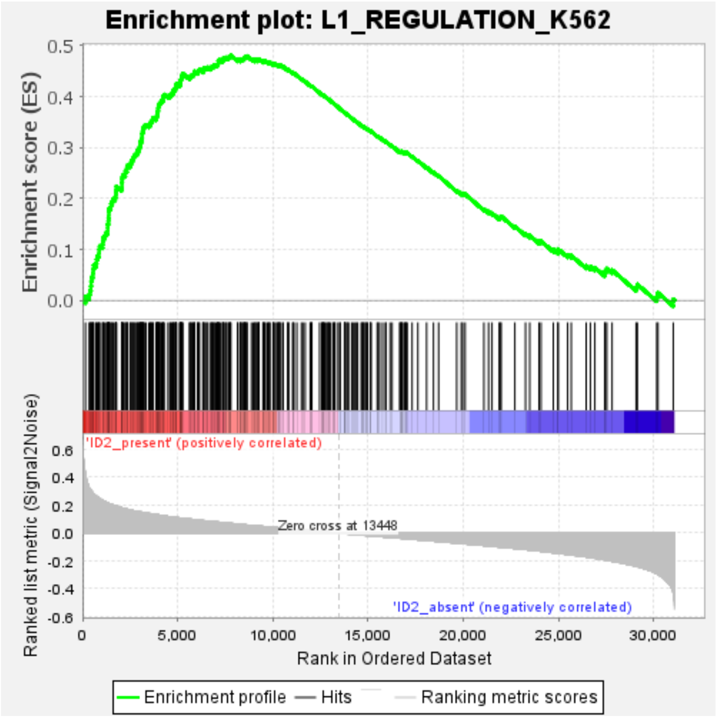


Normalized Enrichment Score (NES) = 2.02
FDR q-value = 0.025

**b**

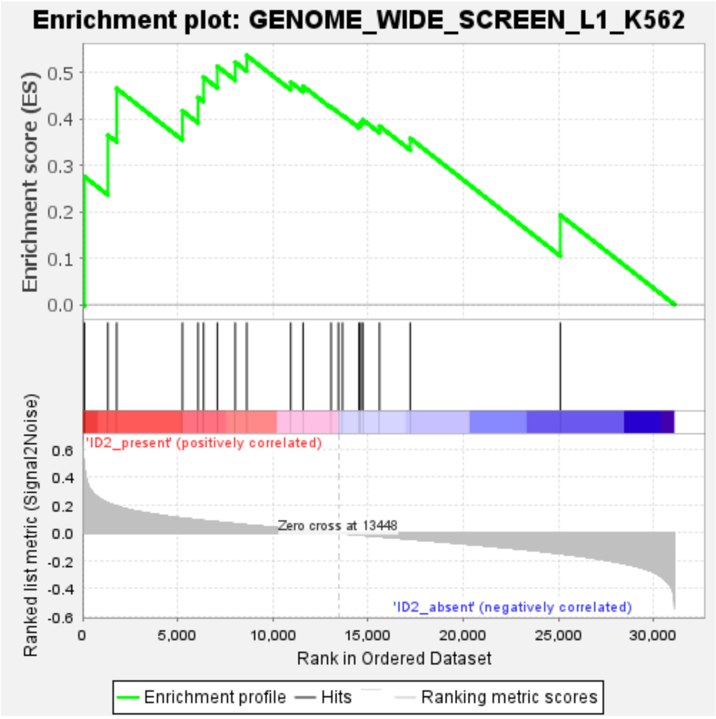


Normalized Enrichment Score (NES) = 1.80
FDR q-value = 0.042

**Supplementary Fig. 29**

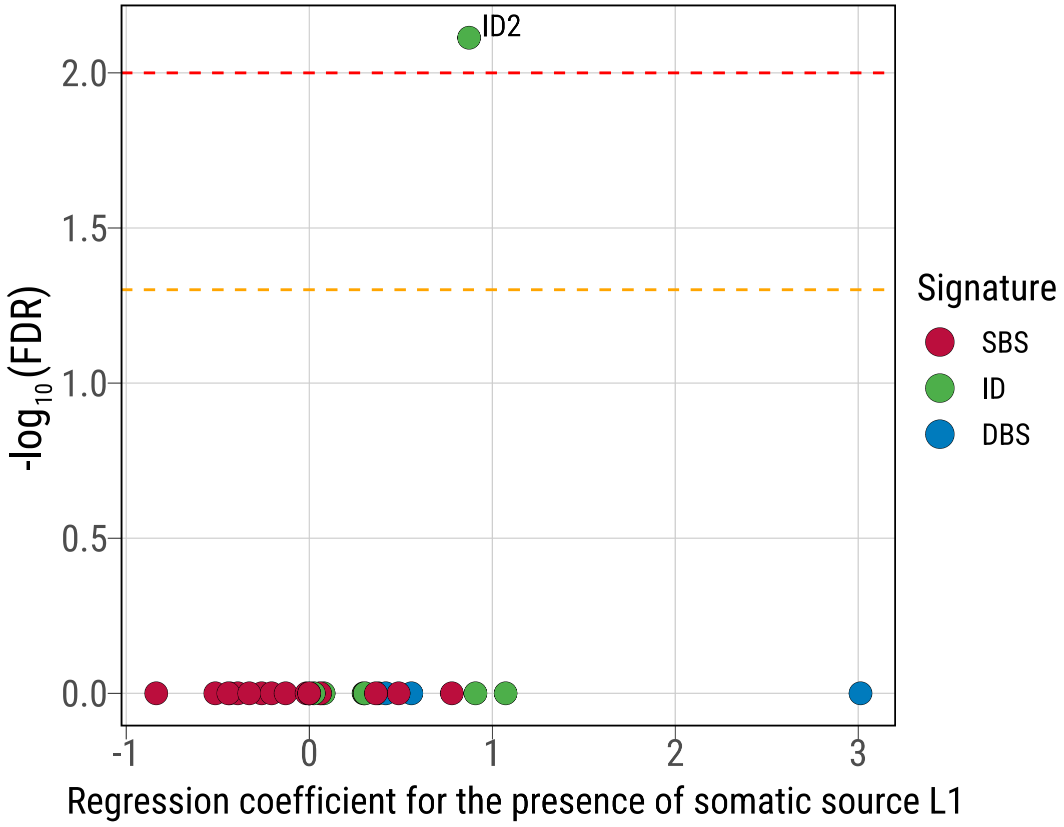

**Supplementary Fig. 30**

a

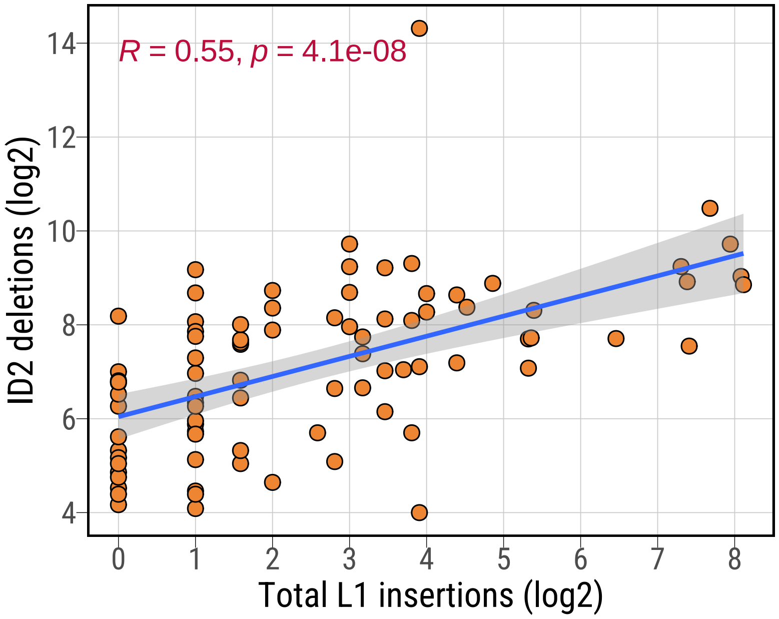


b

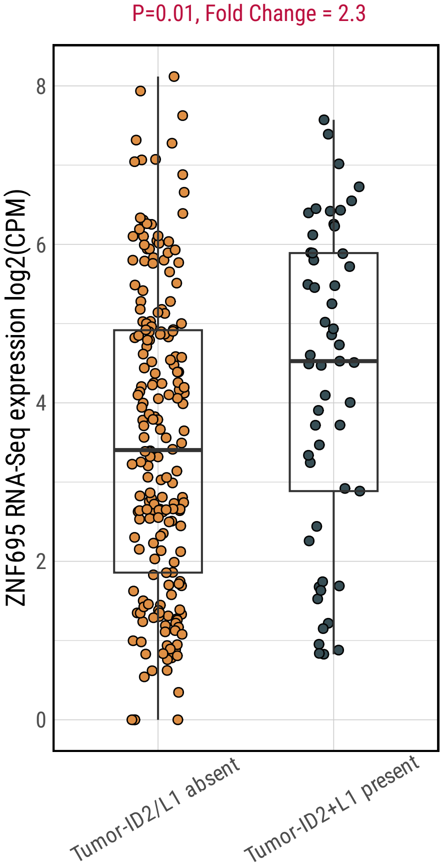

#### Supplementary Fig. 31

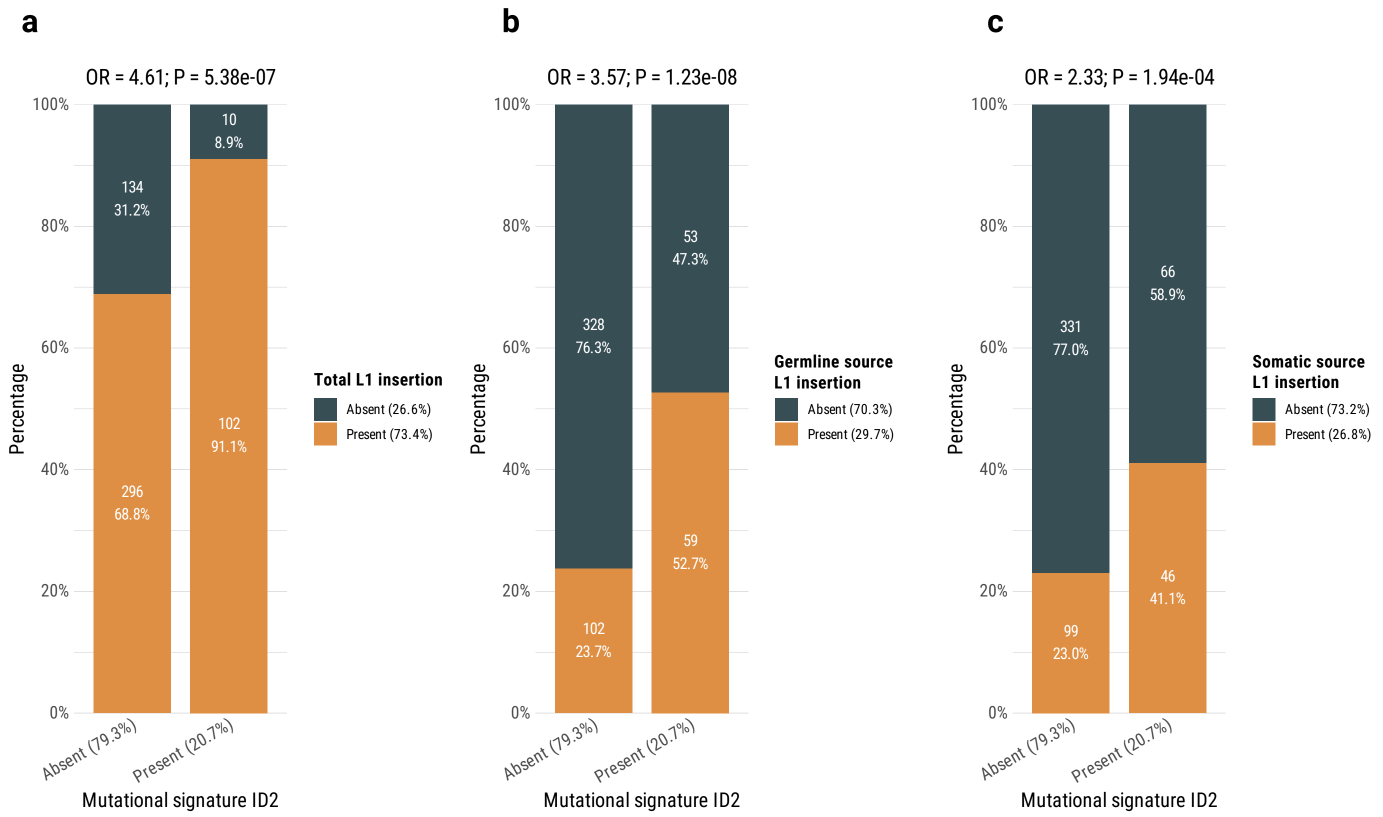

**Supplementary Fig. 32**

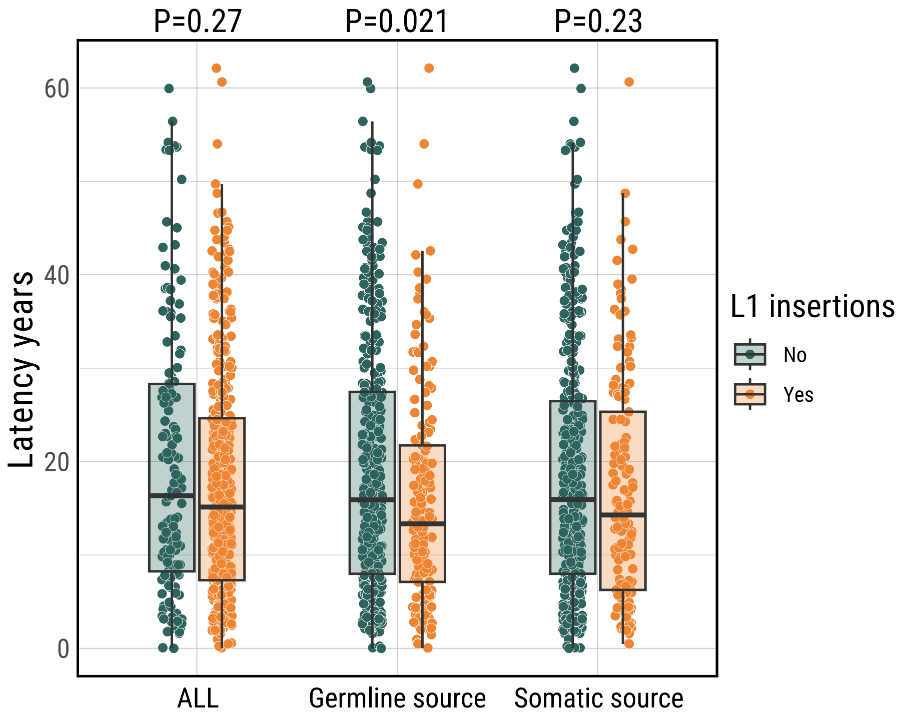

#### Supplementary Fig. 33

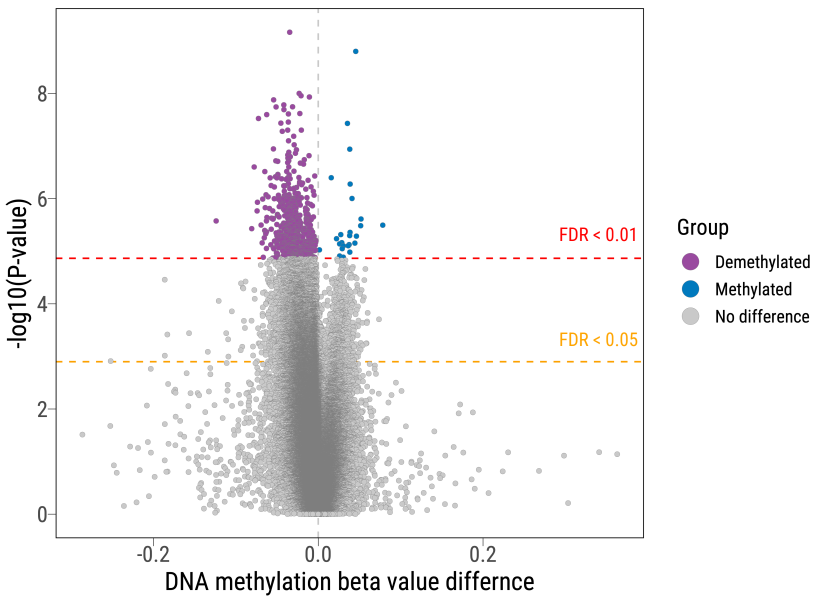

#### Supplementary Fig. 34

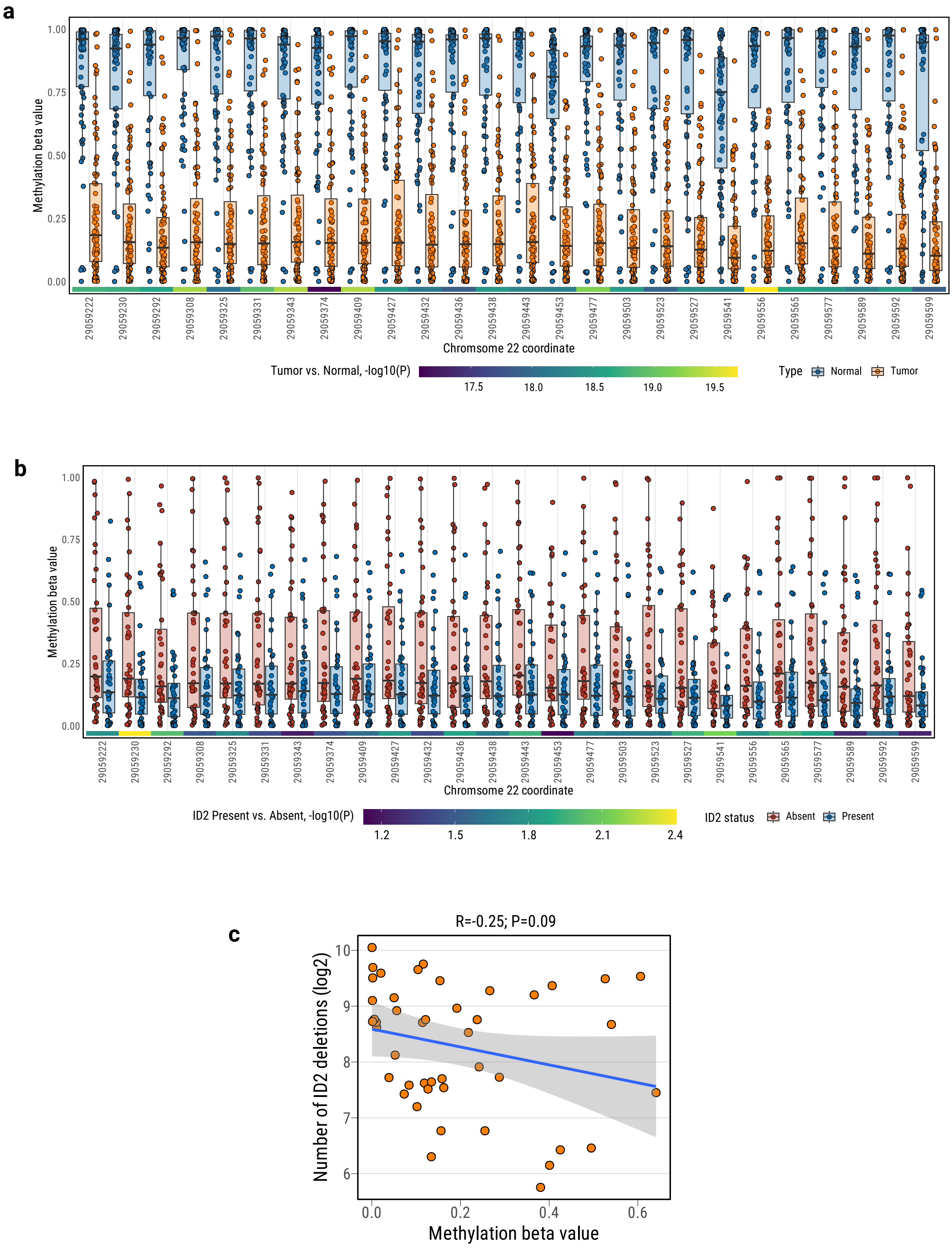

#### Supplementary Fig. 35

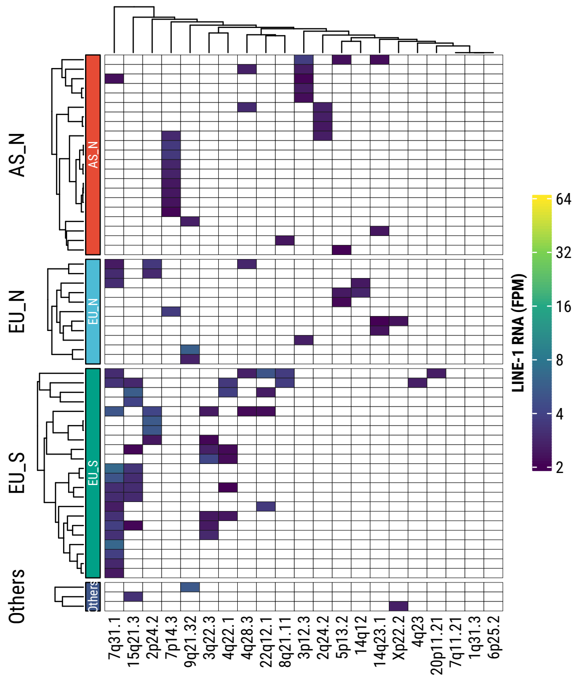


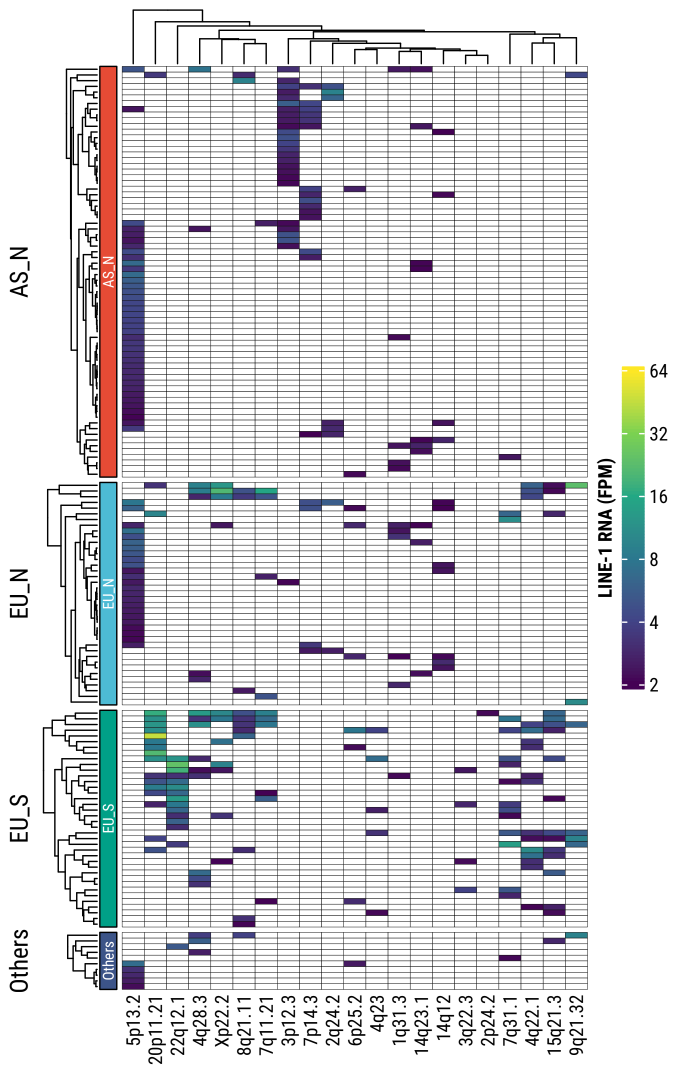

**Supplementary Fig. 36**

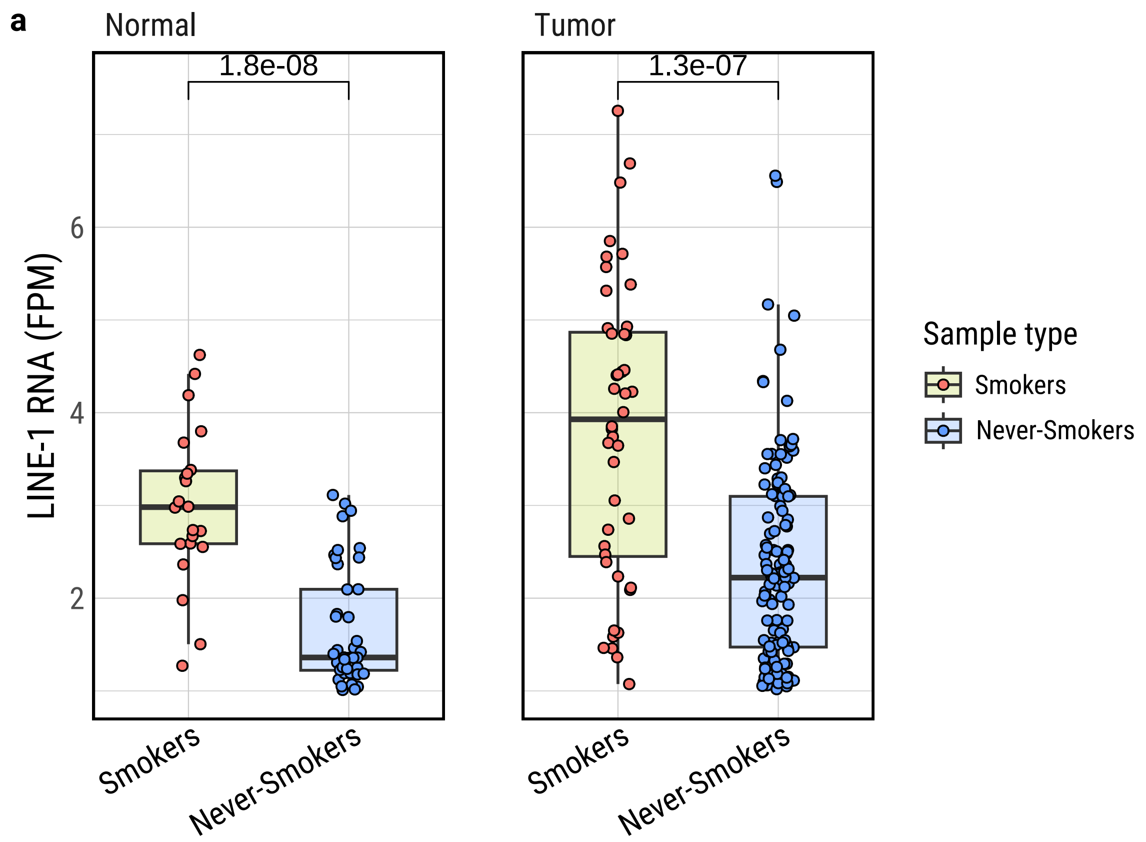


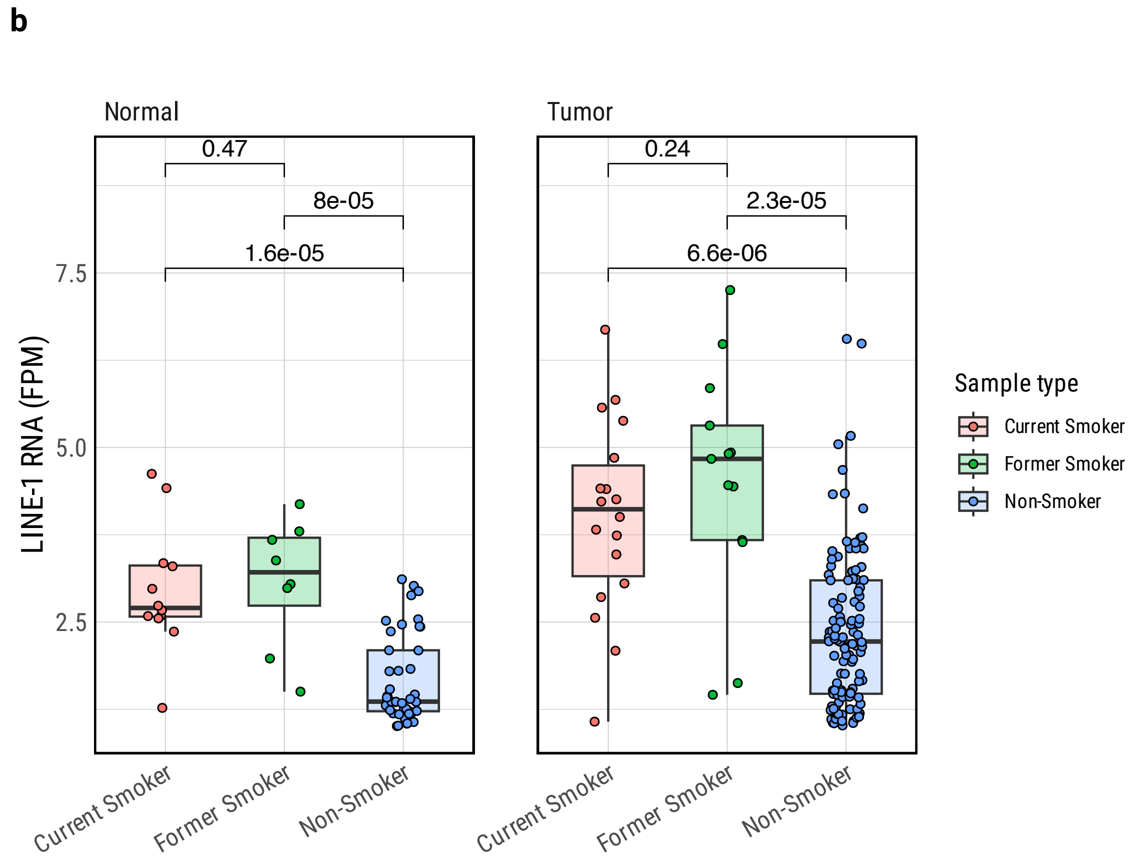

**Supplementary Fig. 37**

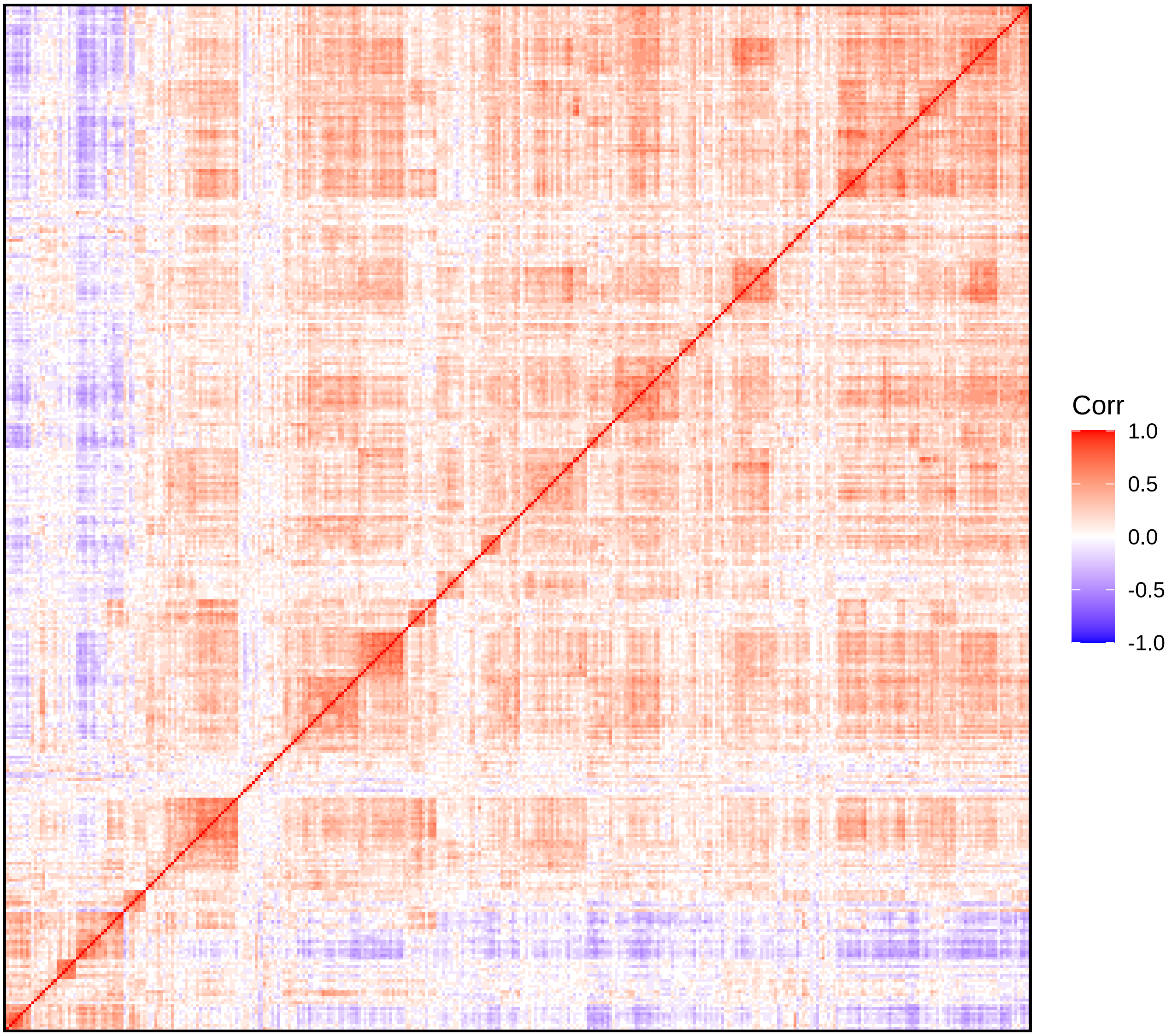

#### Supplementary Fig. 38

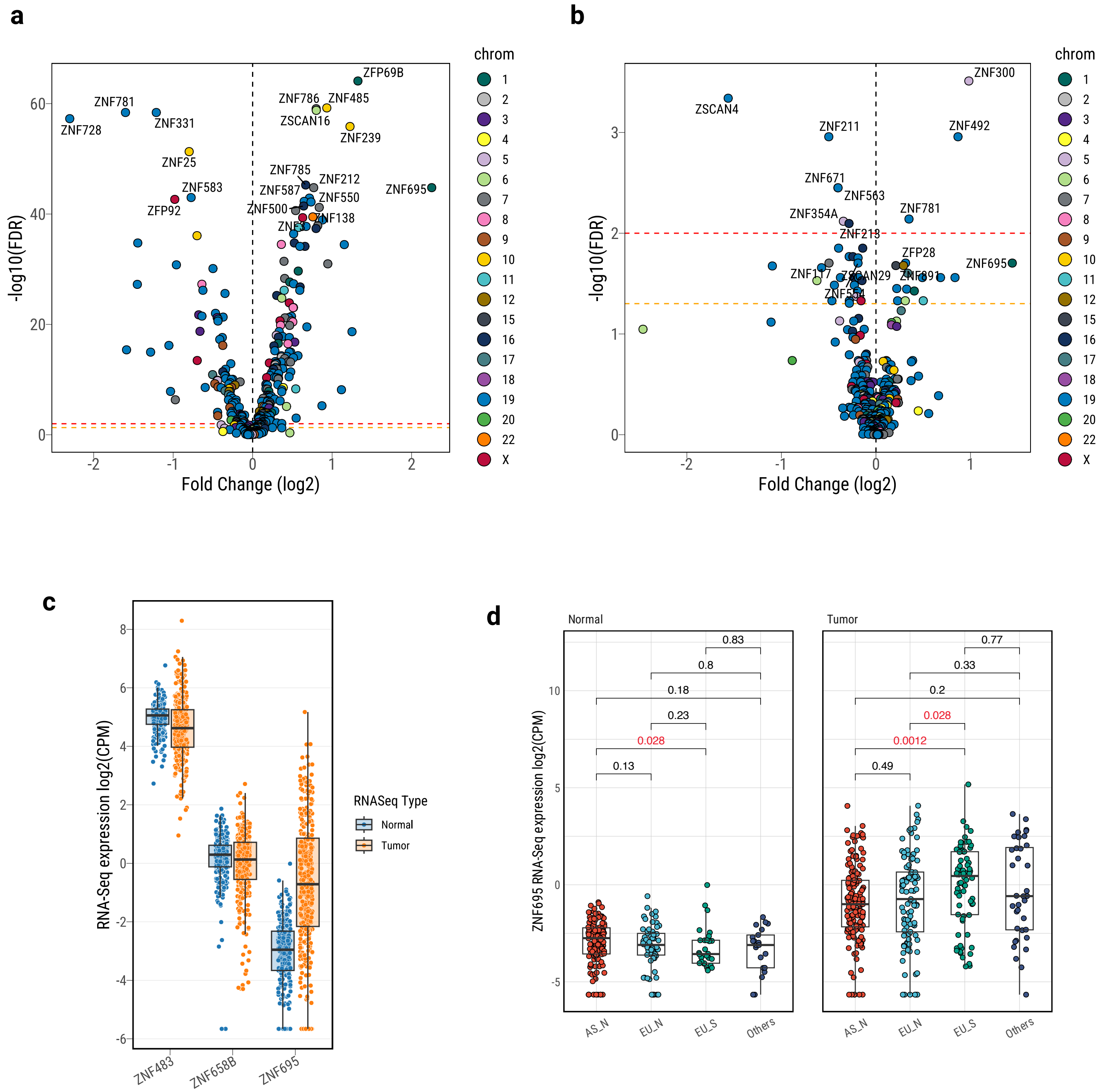

**Supplementary Fig. 39**

a

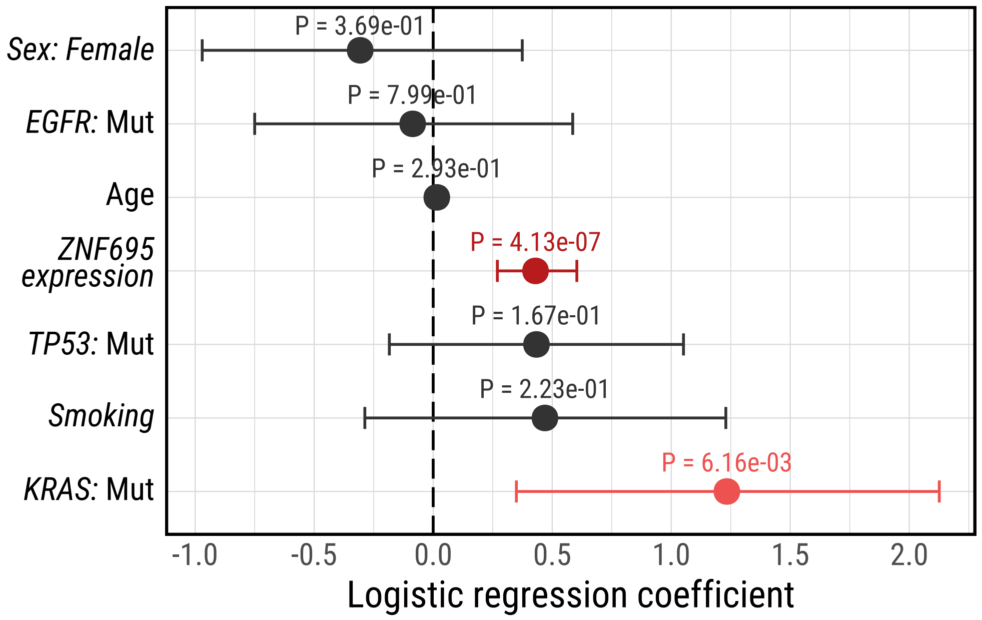


b

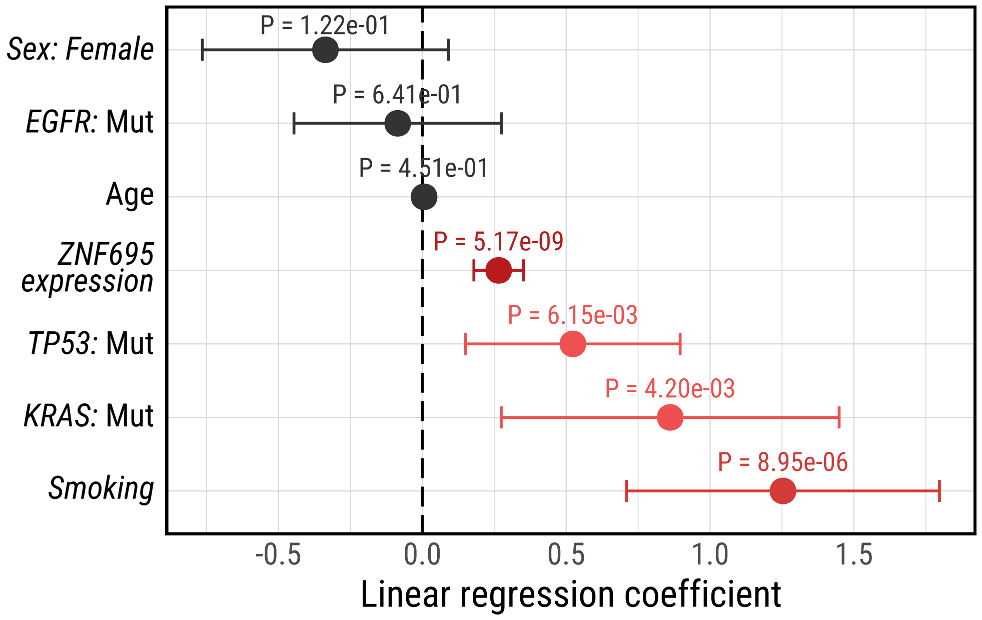

#### Supplementary Fig. 40

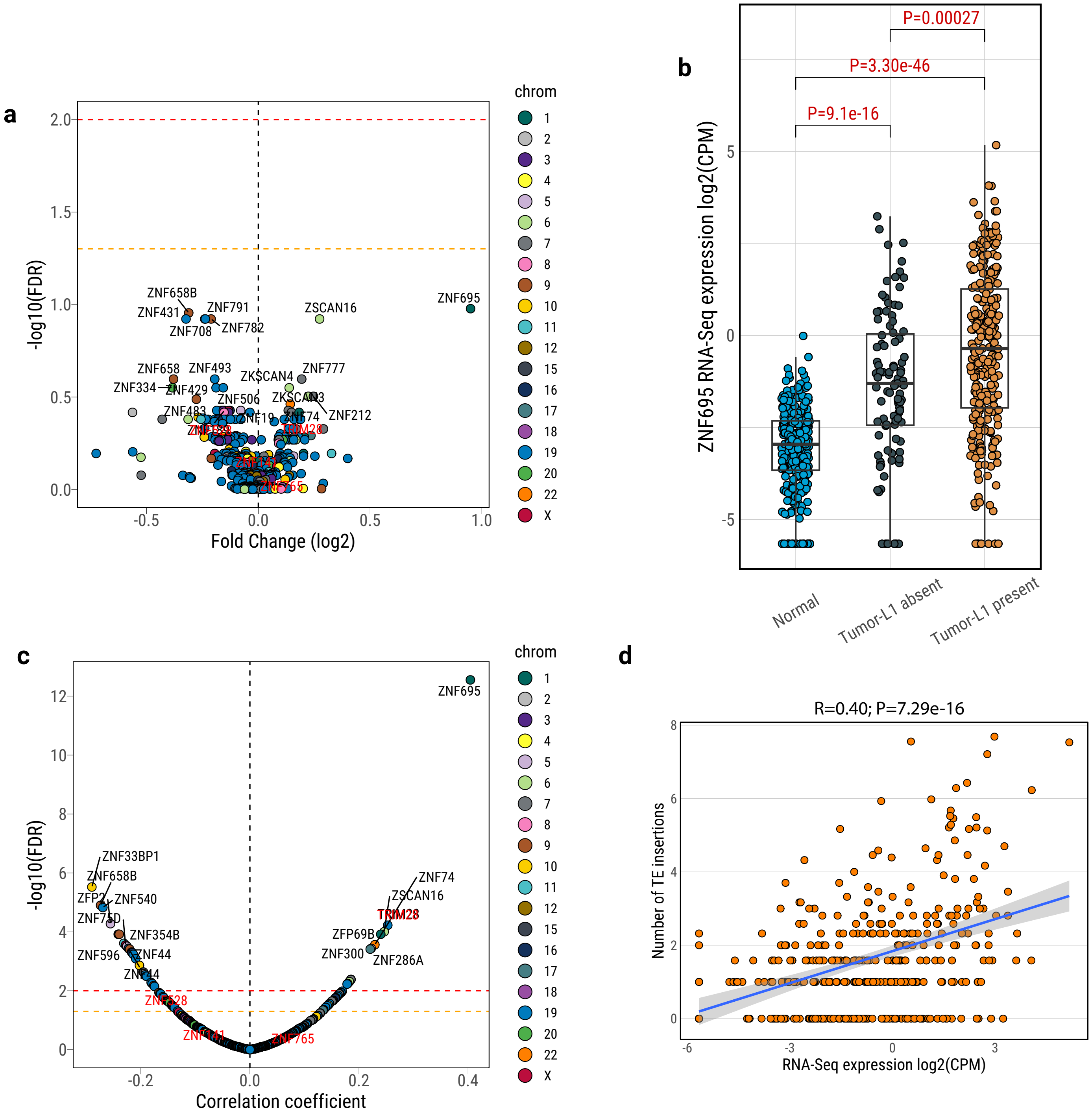

**Supplementary Fig. 41**

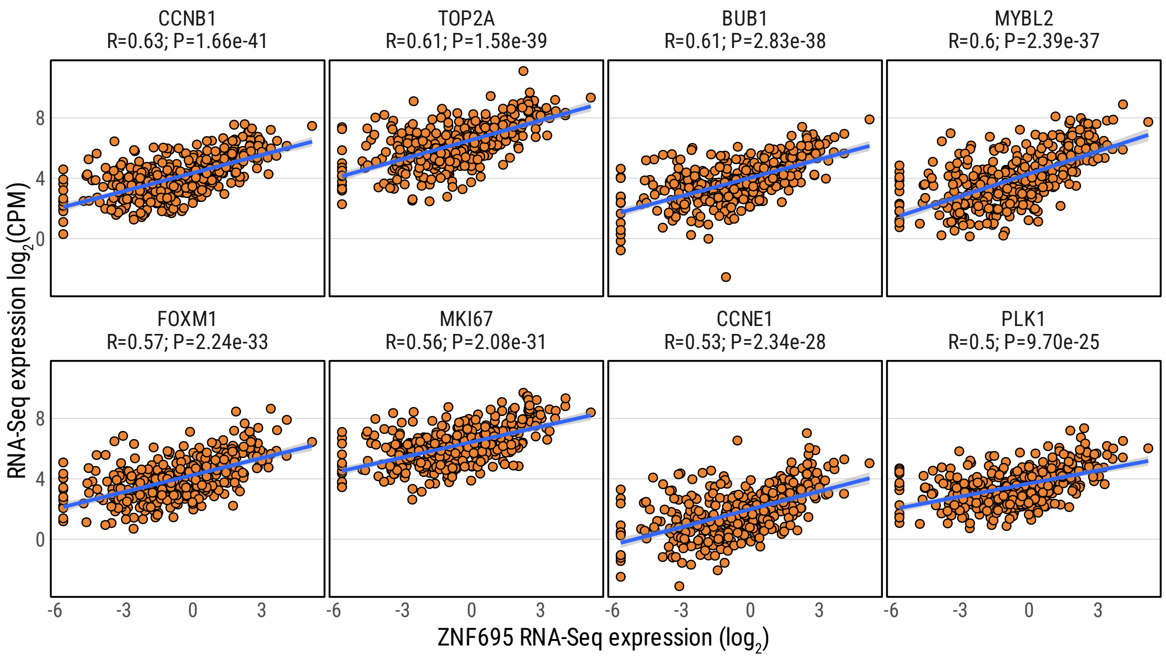

#### Supplementary Fig. 42

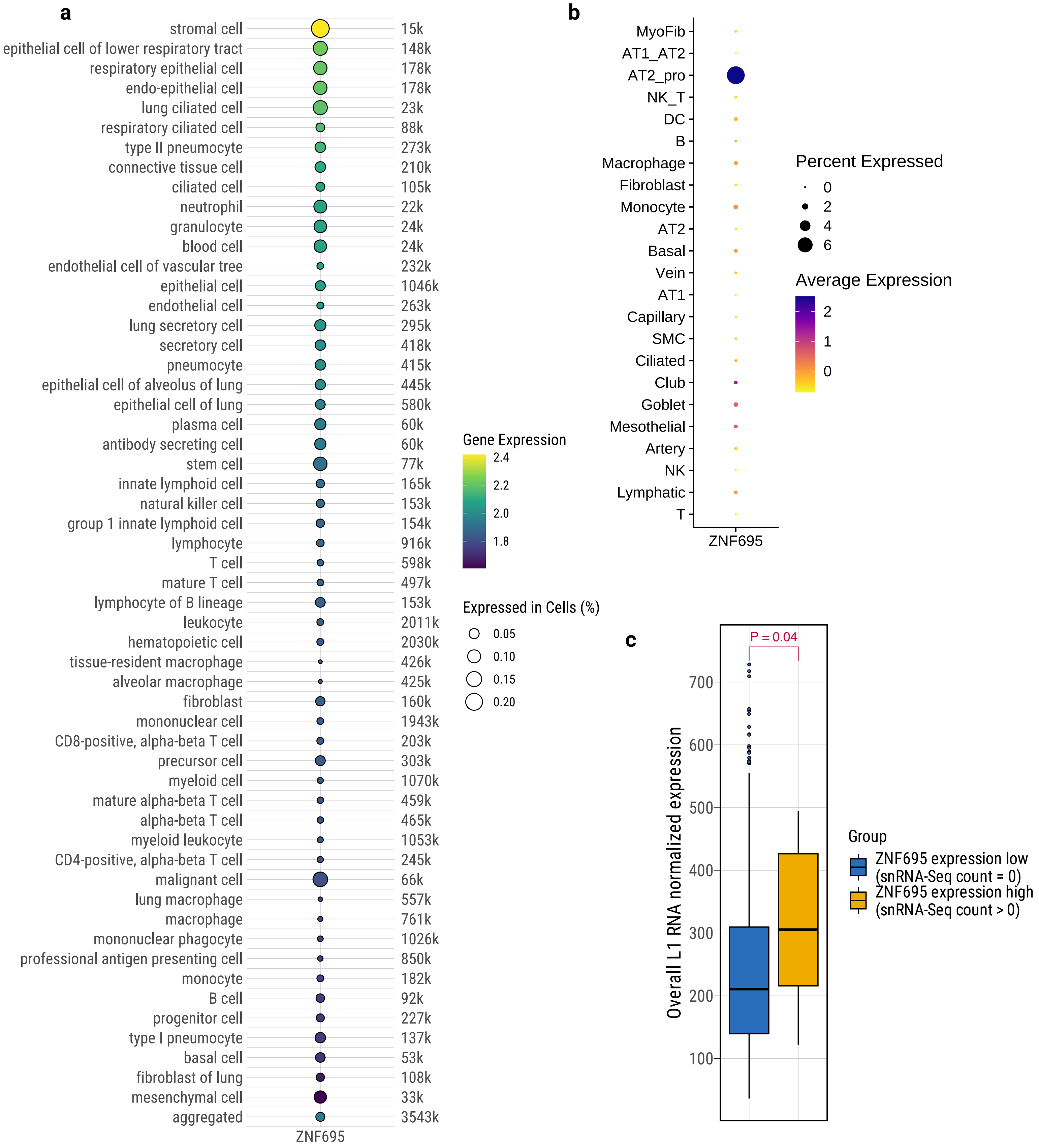

**Supplementary Fig. 43**

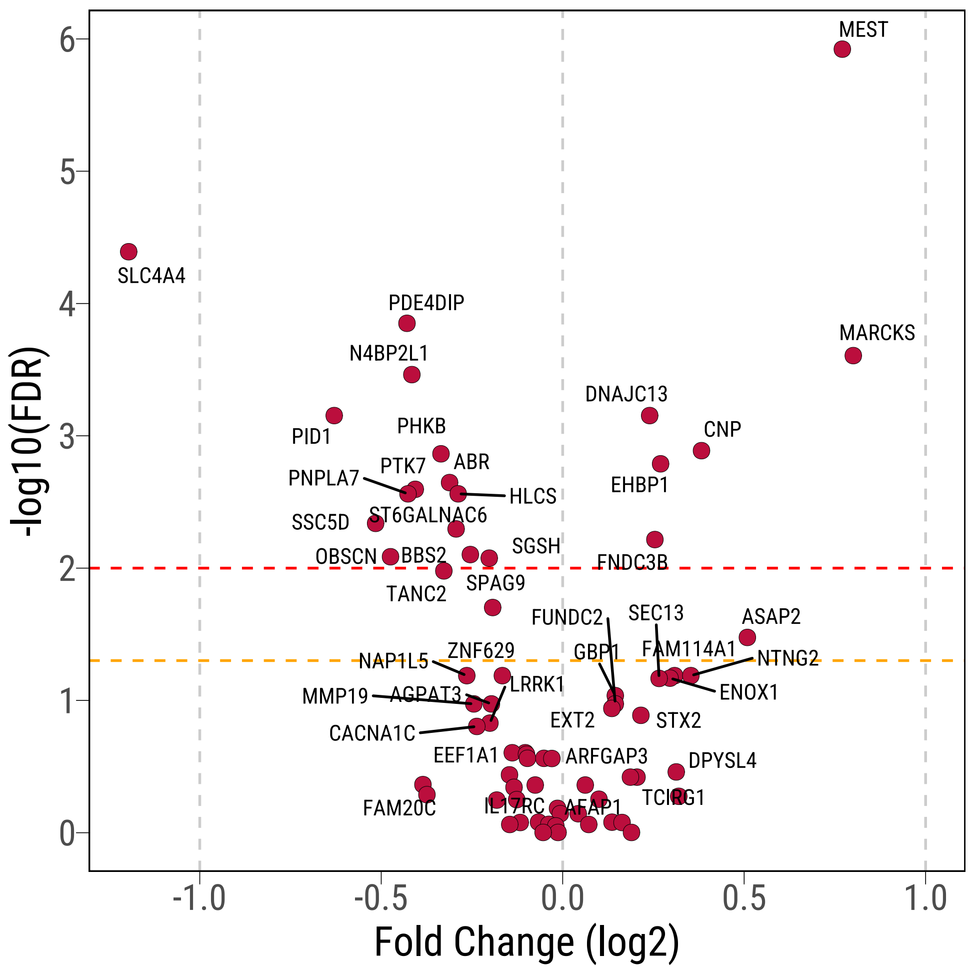

#### Supplementary Fig. 44

Only show junctions with depth >10% of max sequencing depth

**a**

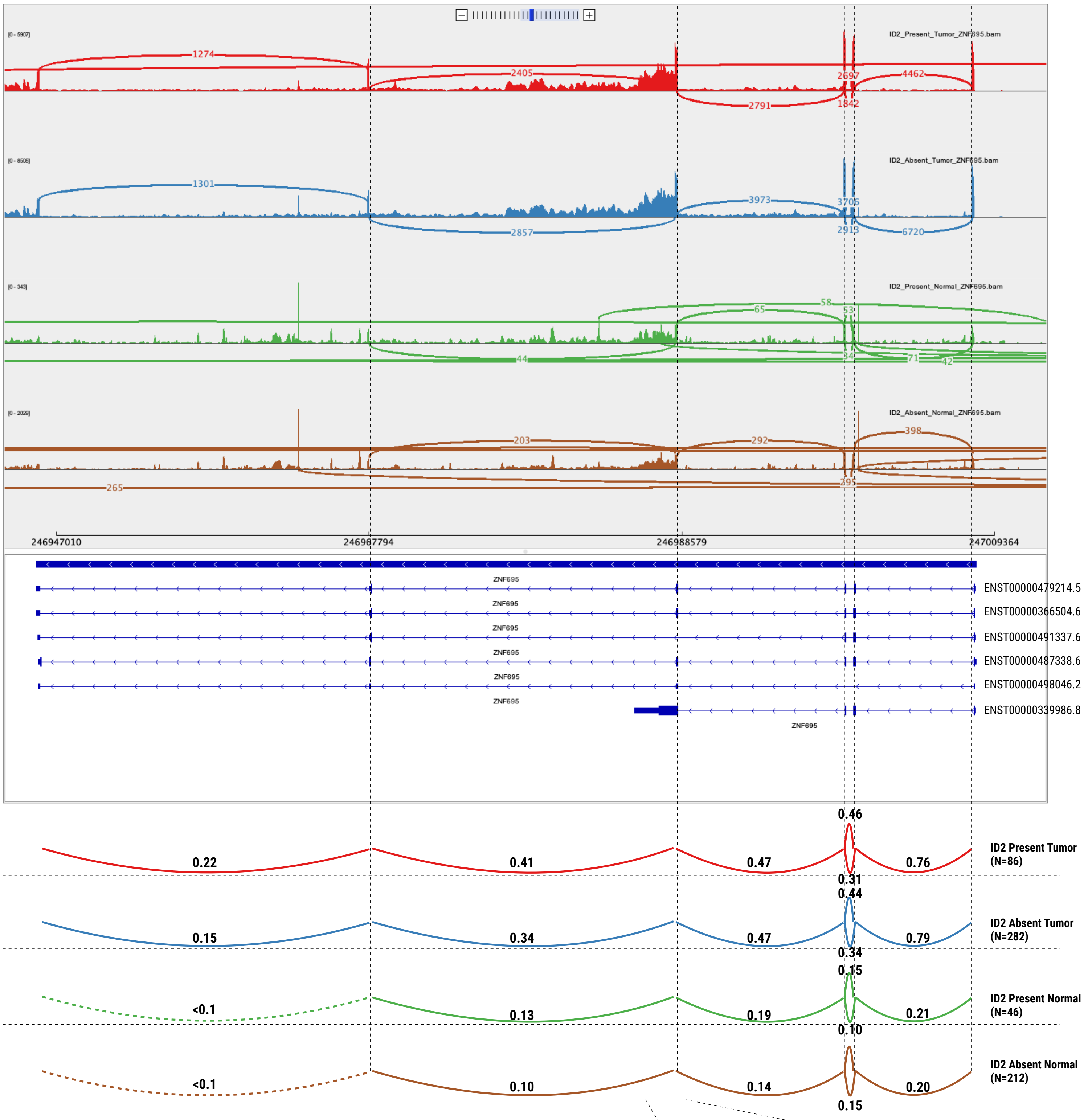


**b**

**c**

| Transcript ID | Transcript Name | Refseq | Amino Acide | Protein Domain | Protein Type |
|---|---|---|---|---|---|
| ENST00000479214.5 | ZNF695-203 | NR_037894 | 121 | KRAB | nonsense_mediated_decay |
| ENST00000366504.6 | ZNF695-202 | - | 133 | KRAB | nonsense_mediated_decay |
| ENST00000491337.6 | ZNF695-205 | - | 88 | KRAB | nonsense_mediated_decay |
| ENST00000487338.6 | ZNF695-204 | NM_001204221 | 172 | KRAB | protein_coding |
| ENST00000498046.2 | ZNF695-206 | | - | - | processed_transcript |
| **ENST00000339986.8** | **ZNF695-201** | NM_020394 | **515** | KRAB and Zinc-finger | **protein_coding** |

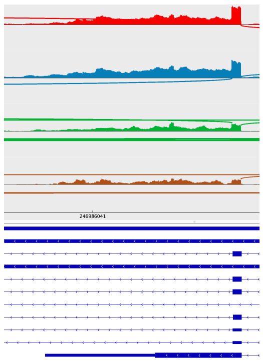

#### Supplementary Fig. 45

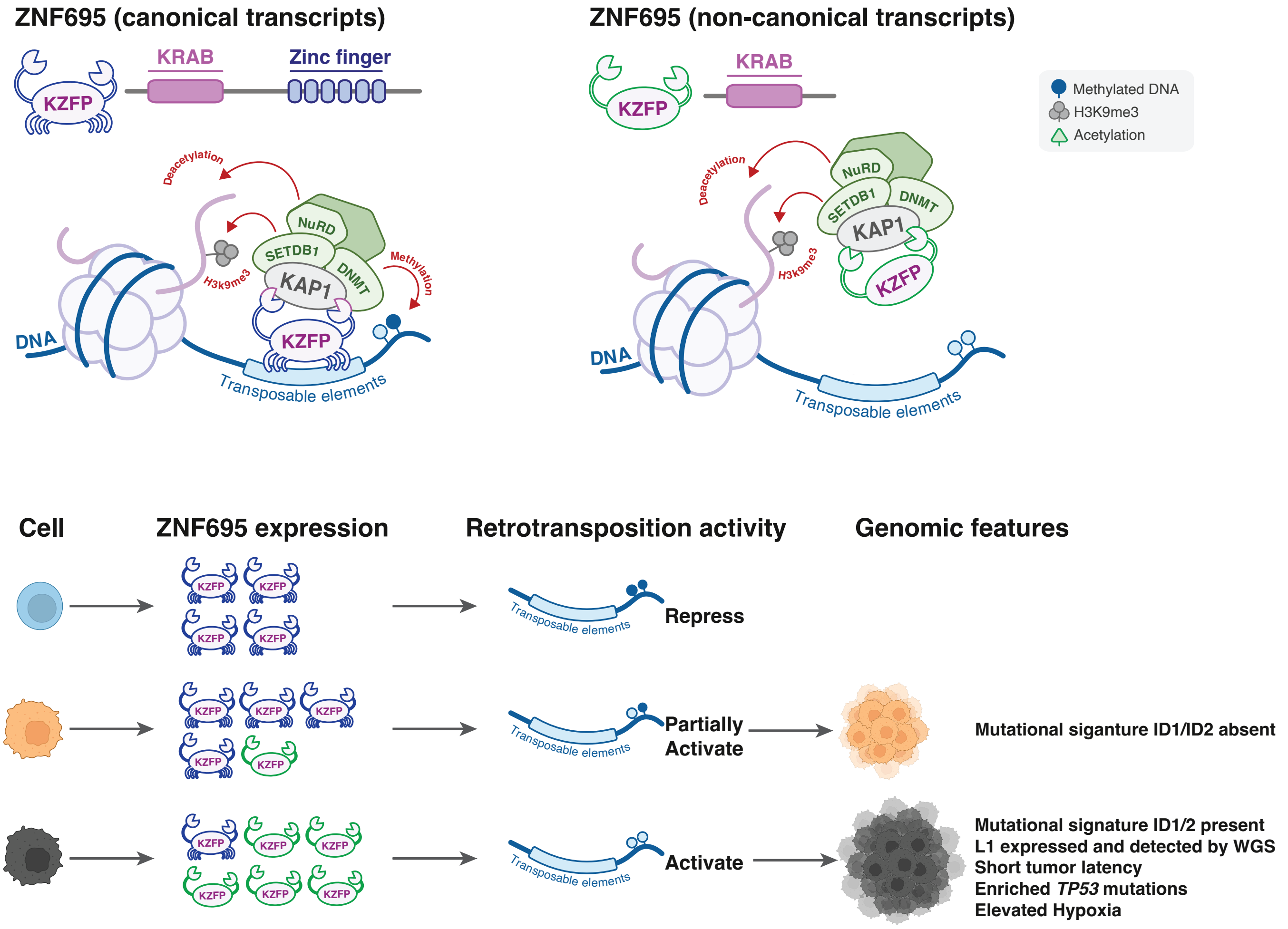
